# An integrated genomic approach to dissect the genetic landscape regulating the cell-to-cell transfer of a-synuclein

**DOI:** 10.1101/2019.12.23.886838

**Authors:** Eleanna Kara, Alessandro Crimi, Anne Wiedmer, Marc Emmenegger, Claudia Manzoni, Sara Bandres-Ciga, Karishma D’Sa, Regina H Reynolds, Juan A Botía, Marco Losa, Veronika Lysenko, Manfredi Carta, Daniel Heinzer, Merve Avar, Andra Chincisan, Cornelis Blauwendraat, Sonia Garcia Ruiz, Daniel Pease, Lorene Mottier, Alessandra Carrella, Dezirae Schneider, Andreia Magalhaes, Caroline Aemisegger, Alexandre P A Theocharides, Zhanyun Fan, Jordan D Marks, Sarah C Hopp, Patrick Lewis, Mina Ryten, John Hardy, Bradley T Hyman, Adriano Aguzzi

## Abstract

Neuropathological and experimental evidence suggests that the cell-to-cell transfer of a-synuclein has an important role in the pathogenesis of Parkinson’s disease (PD). However, the mechanism underlying this phenomenon is not fully understood. We undertook an siRNA, genome-wide high throughput screen to identify genes regulating the cell-to-cell transfer of a-synuclein. We transiently transfected HEK cells stably overexpressing a-synuclein with a construct encoding GFP-2a-aSynuclein-RFP. The cells expressing a-synuclein-RFP through transfection were double positive for GFP and RFP fluorescence, whereas the cells receiving it through transfer were positive only for RFP fluorescence. The amount of a-synuclein transfer was quantified by high content microscopy. A series of unbiased screens confirmed the involvement of 38 genes in the regulation of a-synuclein-RFP transfer. One of those hits was *ITGA8*, a candidate gene recently identified through a large PD genome wide association study (GWAS). Weighted gene co-expression network analysis (WGCNA) and weighted protein-protein network interaction analysis (WPPNIA) showed that the hits clustered in networks that included known PD Mendelian and GWAS risk genes more frequently than expected than random chance. Given the genetic overlap between a-synuclein transfer and PD, those findings provide supporting evidence for the importance of the cell-to-cell transfer of a-synuclein in the pathogenesis of PD, and expand our understanding of the mechanism of a-synuclein spread.

## Introduction

The cell-to-cell transfer of a-synuclein is thought to be an important event in the pathogenesis of Parkinson’s disease (PD). The first evidence suggesting a possible role of this phenomenon came from post mortem analyses of brains from patients with PD who had received neuronal grafts in the midbrain, and showed that those healthy neurons had developed Lewy bodies (Kordower et al., 2008; Li et al., 2008). In addition, patients with PD develop a-synuclein pathology in the brain that follows a consistent pattern starting from the brain stem and olfactory bulb and moving up to the cortex (Braak et al., 2003), which is consistent with the spread of an “agent” that is thought to be misfolded a-synuclein. Those observations have been supported by experimental evidence generated through studies on mouse models and tissue culture systems (Luk et al., 2016; Luk et al., 2012; Luk and Lee, 2014; Luk et al., 2009; Rey et al., 2018; Rey et al., 2016; Thakur et al., 2017; Volpicelli-Daley et al., 2014; Volpicelli-Daley et al., 2011). However, despite the accumulation of evidence supporting the fact that a-synuclein spread does occur, the importance of this phenomenon in the pathogenesis of PD remains controversial. A substantial proportion of PD cases do not follow the Braak staging, and the severity of the clinical presentation often does not correlate with the Braak stage (Surmeier et al., 2017). Therefore, an alternative model that has been suggested is the selective vulnerability hypothesis, according to which the progressive development of pathology is not because of the spread of a pathogenic species of a-synuclein, but because of the differential vulnerability of various brain regions to the disease process (Alegre-Abarrategui et al., 2019; Fu et al., 2018; Hardy, 2016).

The mechanism of cell-to-cell transfer of a-synuclein is incompletely understood. The species of a-synuclein that undergoes transfer, along with the mechanism of the transfer, are unknown. Until recently, it had been thought that the fibrillar form of a-synuclein undergoes cell-to-cell transfer in the disease, and it is capable of inducing seeding and misfolding of the endogenous a-synuclein (Aguzzi et al., 2007; Aguzzi and Rajendran, 2009; Guo and Lee, 2014). However, recent evidence has challenged this hypothesis. It has been shown that oligomeric and monomeric a-synuclein can also undergo transfer and induce the formation of pathology after intracerebral injections in mice through mechanisms other than seeding of endogenous template (Rey et al., 2013; Sacino et al., 2013). Cell-to-cell transfer can also occur without the presence of endogenous template (in the case of tau, another protein whose behavior is thought to be similar to that of a-synuclein) (Wegmann et al., 2015) and can cross species barriers (Luk et al., 2012), therefore suggesting a clear separation between true prions and prionoids.

Therefore, there are several critical questions regarding the spread of a-synuclein. First, what are the genetic underpinnings and mechanisms underlying the a-synuclein spread, do those relate to known genetic causes of PD, and if yes, how? Second, what is the importance of this phenomenon in the pathogenesis of PD?

To address those questions, we performed a high throughput, siRNA genome-wide screen to find genes regulating the cell-to-cell transfer of a-synuclein. Given the uncertainty regarding the nature of the transferred species of a-synuclein, we used a novel genetically encoded reporter to identify the cells that received the transferred a-synuclein, which does not make any assumptions regarding the aggregation state of the transferred protein. The screen identified 38 genes regulating this process, one of which was *ITGA8*, a PD risk gene recently identified through GWAS (Nalls et al., 2019). Genes involved in cell signaling, protein transport and protein catabolism, among others, were enriched within the cohort. Weighted gene coexpression network analysis (WGCNA) and weighted protein-protein network interaction analysis (WPPNIA) showed a significant overlap between the networks formed by the 38 genes and the networks formed by PD Mendelian and risk genes. Collectively, those findings provide insight into the mechanisms regulating the cell-to-cell transfer of a-synuclein. Our findings also suggest an overlap between the molecular underpinnings of a-synuclein transfer and the genetic causes of PD, indicating that the cell-to-cell transfer of a-synuclein probably has a crucial role in the pathogenesis of PD.

## STAR Methods

### KEY RESOURCES TABLE

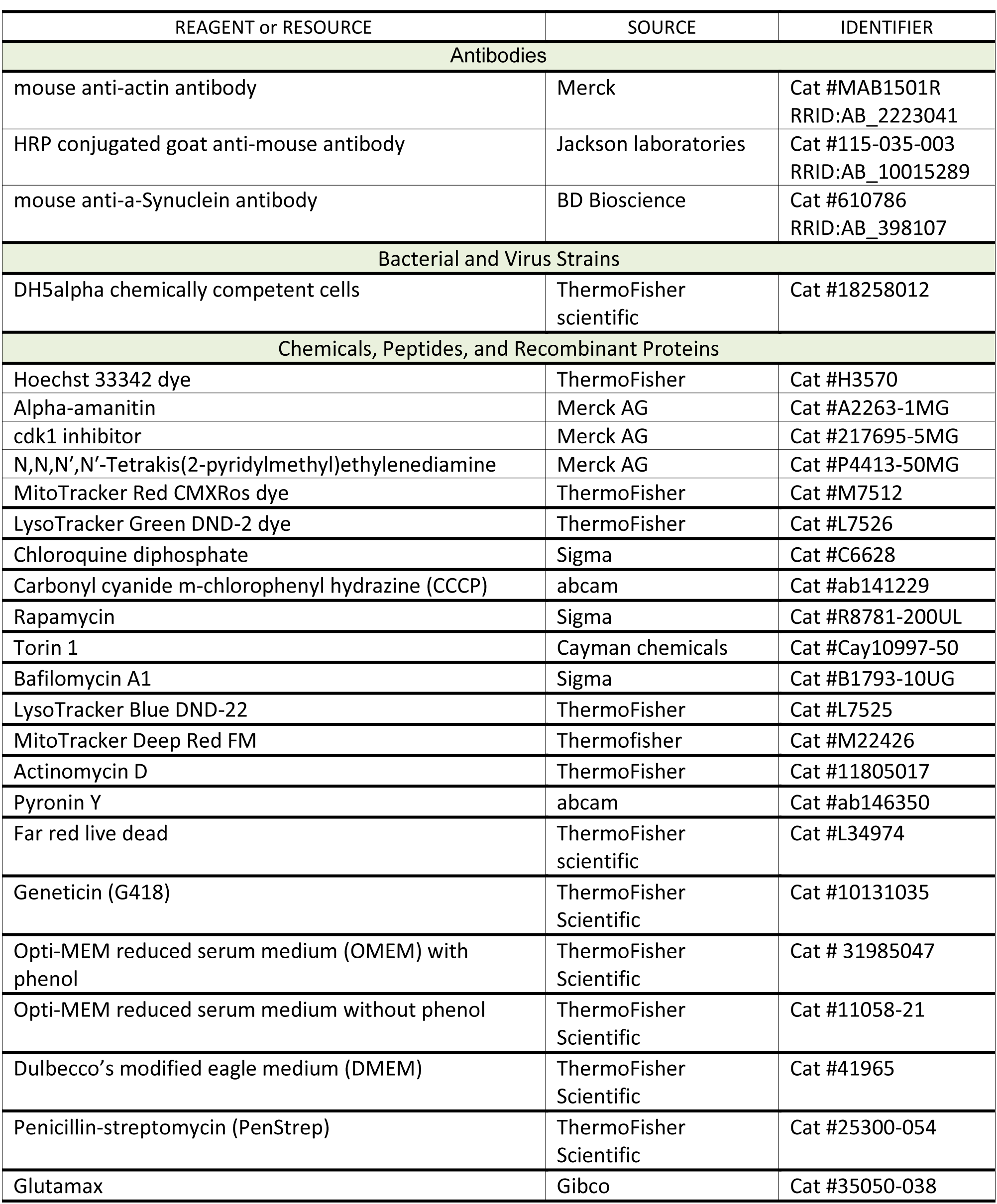
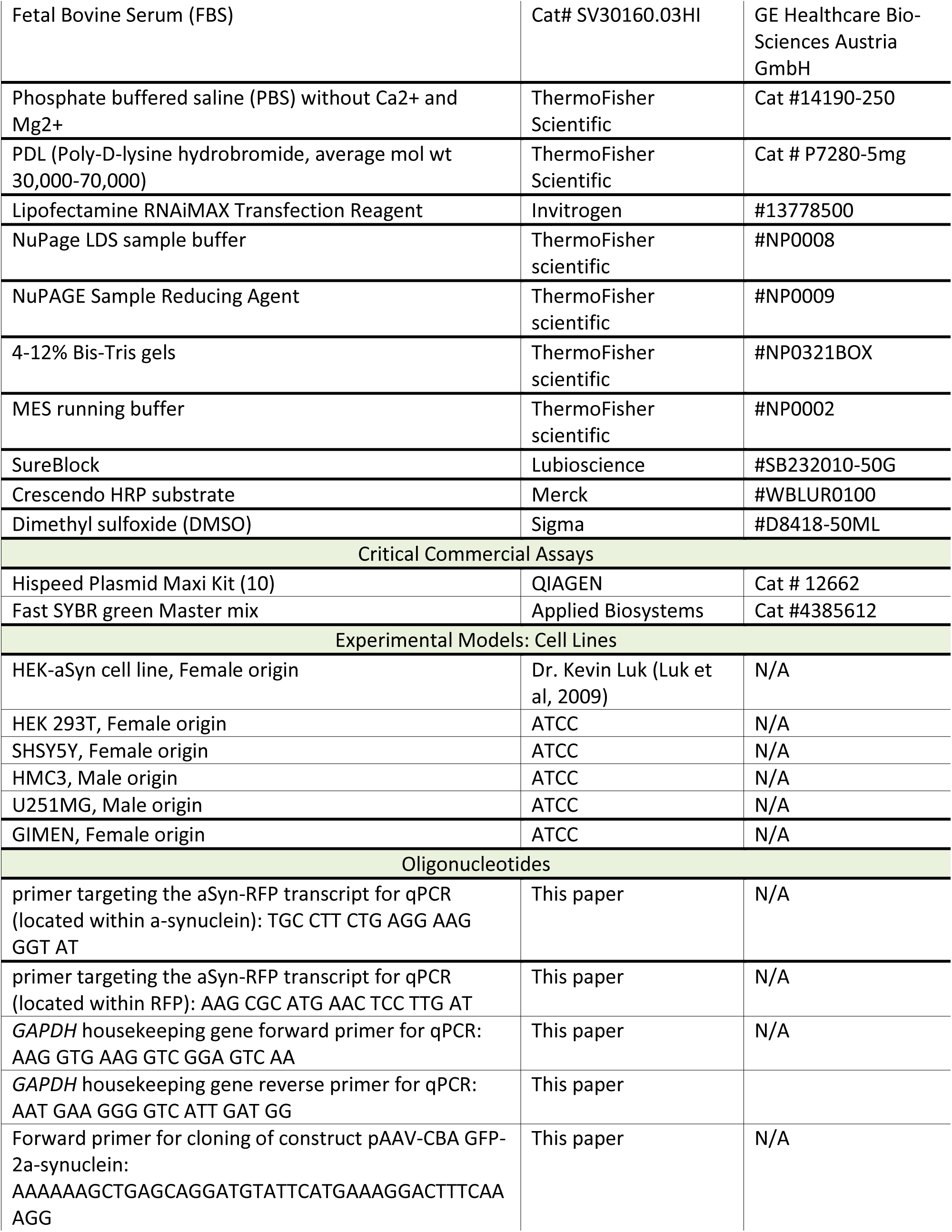
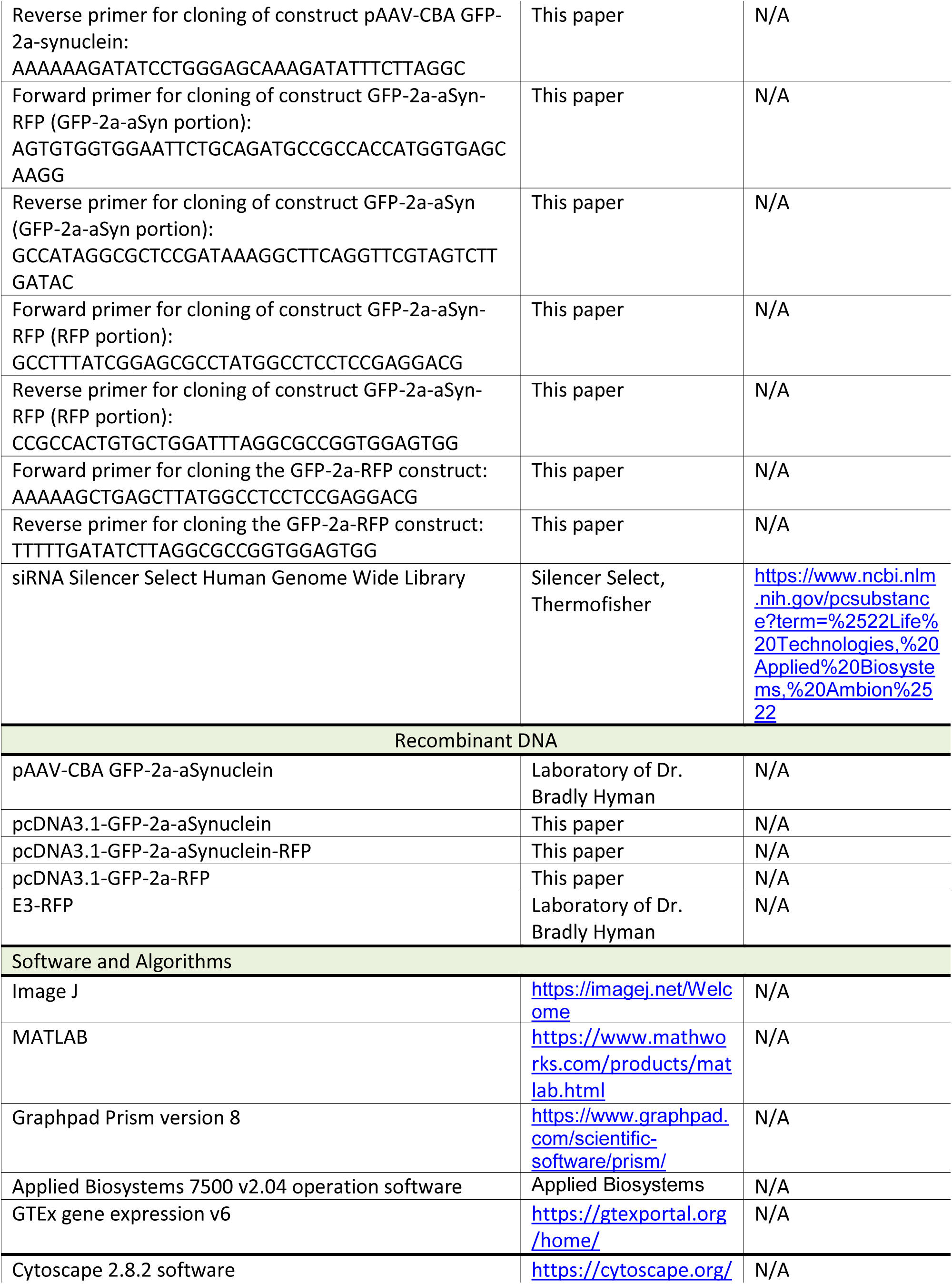
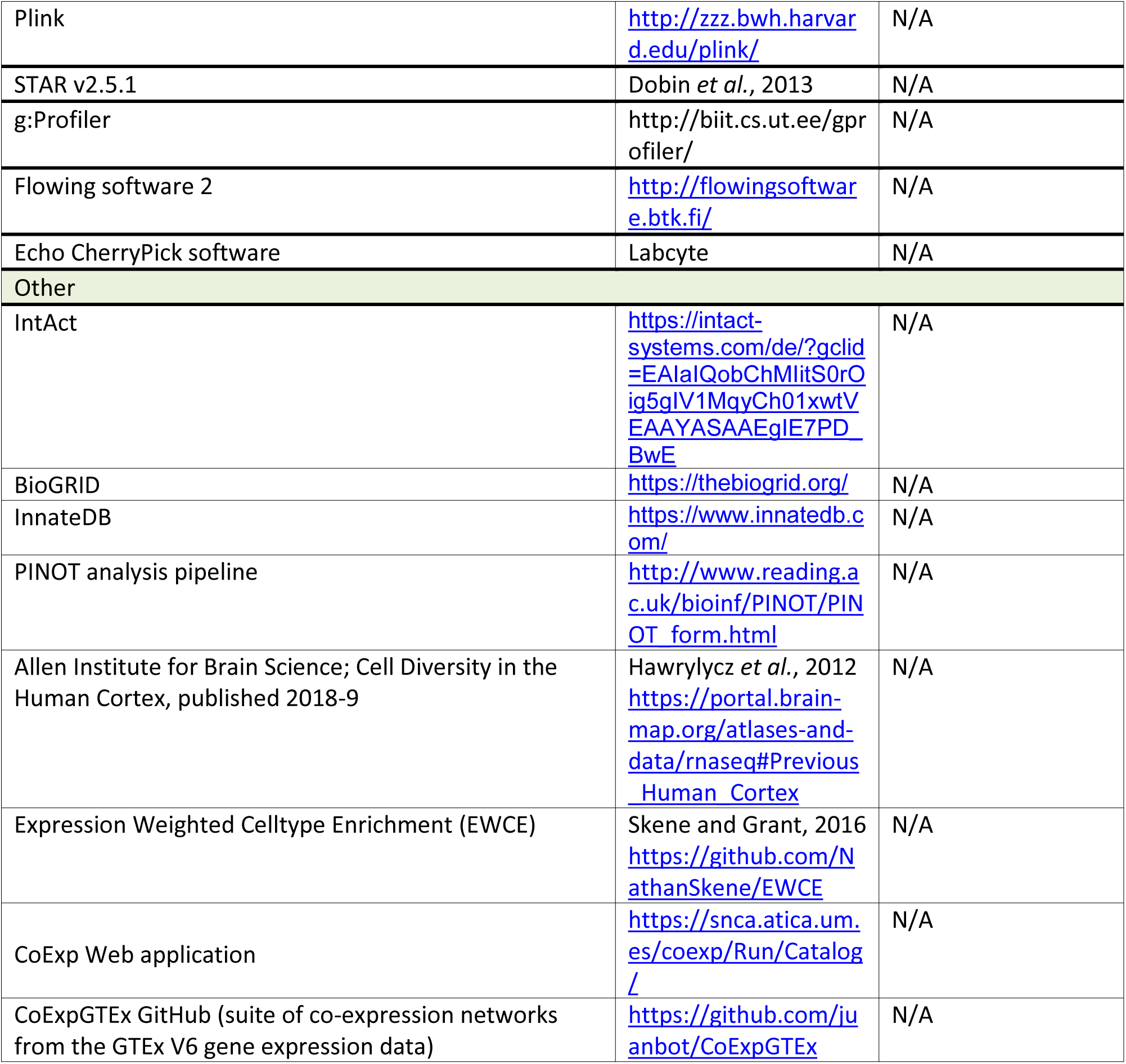

### LEAD CONTACT AND MATERIALS AVAILABILITY

Further information and requests for resources and reagents should be directed to and will be fulfilled by the Lead Contact, Prof. Adriano Aguzzi.

### EXPERIMENTAL MODEL AND SUBJECT DETAILS

#### Tissue culture

A HEK QBI cell line stably overexpressing wild type (WT) a-synuclein (hereafter referred to as HEK-aSyn line) was used in this study (Luk et al., 2009). This cell line was a kind gift from Dr Kelvin Luk (university of Pennsylvania). That cell line was cultured in the following medium: DMEM (#31053-036, ThermoFisher Scientific) + 10%FBS (Hyclone Heat inactivated, #SV30160.03HI, GE Healthcare Bio-Sciences Austria GmbH) + 1%glutamax (#35050-038, Gibco) + 0.2mg/ml geneticin (#10131035, ThermoFisher Scientific).

The following cell lines were also used to assess the cell-to-cell transfer of a-synuclein: U251MG, GIMEN, SHSY5Y, HMC3, HEK 293T. They were cultured in the following media: DMEM (#41965, ThermoFisher scientific) + 10%FBS + 1%glutamax + 1% penicillin-streptomycin (penstrep) (#25300-054, ThermoFisher scientific) (SHSY5Y, HEK 293T), OMEM (31985047, ThermoFisher scientific) + 10%FBS + 1%glutamax + 1%penstrep (GIMEN, HMC3, U251MG).

All cell lines were grown at 37 degrees.

## METHOD DETAILS

### Molecular clonings

#### pAAV-CBA GFP-2a-synuclein

The pAAV-CBA-GFP-2a-tau construct (Wegmann et al., 2019; Wegmann et al., 2015) was used as a vector and was digested with EcoRV and BlpI. The insert was PCR amplified from the pcDNA3.1-aSynuclein construct using the following primers: forward: AAAAAAGCTGAGCAGGATGTATTCATGAAAGGACTTTCAAAGG, reverse: AAAAAAGATATCCTGGGAGCAAAGATATTTCTTAGGC. The vector and insert were ligated, transformed using DH5alpha chemically competent cells (#18258012, ThermoFisher), and the sequence of the miniprep product was confirmed through Sanger sequencing.

#### pcDNA3.1-GFP-2a-aSyn

The pcDNA3.1 vector was digested with HindIII and EcorV enzymes, followed by CIP treatment. The pAAV-CBA-GFP-2a-aSyn construct was digested with the same enzymes to isolate the GFP-2a-aSyn insert. Vector and insert were ligated, transformed using DH5a chemically competent cells, and the sequence of the miniprep product was confirmed through Sanger sequencing.

#### GFP-2a-synuclein-RFP

The pcDNA3.1 backbone was digested with EcoRI, followed by CIP treatment. The GFP-2a-aSyn and RFP fragments were amplified through PCR using the following primers: GFP-2a-aSyn forward: AGTGTGGTGGAATTCTGCAGATGCCGCCACCATGGTGAGCAAGG; GFP-2a-aSyn reverse: GCCATAGGCGCTCCGATAAAGGCTTCAGGTTCGTAGTCTTGATAC; mRFP forward: GCCTTTATCGGAGCGCCTATGGCCTCCTCCGAGGACG; mRFP reverse: CCGCCACTGTGCTGGATTTAGGCGCCGGTGGAGTGG. Those fragments were ligated into the backbone through Gibson assembly, as per manufacturer instructions. Of note, monomeric RFP (mRFP) was used in this and in the two constructs described below (Campbell et al., 2002).

#### GFP-2a-RFP

The pcDNA3.1-GFP-2a-aSyn vector was digested with BlpI and EcorV-HF to remove the aSyn fragment, followed by CIP treatment and gel purification. The mRFP insert was prepared through PCR amplification using pcDNA3.1-GFP-2a-aSyn-RFP as a template and the following primers: forward: AAAAAGCTGAGCTTATGGCCTCCTCCGAGGACG, reverse: TTTTTGATATCTTAGGCGCCGGTGGAGTGG. The digested vector and insert were ligated and transformed with DH5a chemically competent cells. The sequence of the miniprep products was confirmed through Sanger sequencing.

The cloning of the E3-RFP construct has been previously described (Kara et al., 2017; Kara et al., 2018). All maxipreps were done using the Hispeed plasmid Maxi Kit (#12662, Qiagen).

### Flow cytometric assay to quantify the cell-to-cell transfer of a-synuclein

The HEK-aSyn cells were plated in 6 well plates at a density of 870 cells/ul. For the experiment where siRNAs were co-transfected with the constructs, the cells were plated in 24 well plates at a density of 870 cells/ul. 48h later, the cells were transfected with the GFP-2a-aSyn-RFP or the GFP-2a-RFP constructs, plus the siRNAs, as applicable. As single color controls for flow cytometry, the E3-RFP and GFP-2a-aSyn constructs were used. Before transfection, the medium in the 6 well plates was replaced with pure OMEM without phenol (#11058-21, ThermoFisher Scientific) by adding 1ml per well. The medium in the 24 well plates was replaced with the following medium, 440ul per well: pure OMEM without phenol plus 10%FBS plus 1%penstrep. For the transfection in 6 well plate format, 2 tubes were prepared, with the following amounts per well: One tube containing 150ul of pure OMEM without phenol plus 3ug of construct, and one tube containing 150ul of pure OMEM plus 9ul of RNAiMax (#13778500, Invitrogen). After 5min of incubation at room temperature, the contents of the tubes were mixed and 300ul of the mix was added per well. For the transfections in 24 well plate format, the following 2 tubes were prepared per well: One tube containing 30ul of pure OMEM plus 0.4ug of construct plus 0.5ul of stock siRNA solution with a concentration of 5uM, and a second tube containing 30ul of pure OMEM plus 1.8ul of RNAiMax. The final concentration of the siRNA in the tissue culture medium after addition to the cells was 5nM. After 5min of incubation at room temperature, the contents of the tubes were mixed and 60ul of the mix was added per well. 24h after transfection, the medium in the 6 well plates was replaced with the regular growth medium mentioned above. The medium in the 24 well plates was not replaced after transfection. On the 5th day after transfection, where applicable, Hoechst 33342 dye (#H3570, ThermoFisher) was added to PBS -/- (#14190-250, ThermoFisher Scientific) at a dilution of 1:2000. The medium was aspirated from the cells and was replaced with equal volume of the diluted Hoechst staining solution, followed by incubation at 37 degrees in the dark. The staining solution was then aspirated. At the same time, the medium from all wells that were not treated with Hoechst was also aspirated. The cells were washed with PBS -/-, trypsin was added and incubated at 37 degrees for 3min, regular growth medium (containing FBS) was added to inactivate trypsin, the cells were pipetted up and down to detach from the well and transferred to sterile eppendorf tubes, centrifuged at 1500rpm for 7min, and the supernatant was aspirated. At that point, the samples that required staining with far red live dead (#L34974, ThermoFisher scientific) were treated as follows: the stock live dead stain was reconstituted in 50ul of Dimethyl sulfoxide (DMSO) and diluted 1:1000 in PBS-/-. The pelleted cells were resuspended in 1000ul of that solution and incubated at room temperature, in the dark, for 30min. The samples were then centrifuged at 1500rpm for 7min. At that point, 500ul of 2% Paraformaldehyde (PFA) (in PBS-/-) was added to the cell pellet of all samples processed. The samples were stored at 4 degrees in the dark until analysis by flow cytometry.

Cells stained only with Hoechst or siRNA-cy5 or live dead far red were used as single color controls, where applicable. Of note, the far red live dead staining was never combined with the siRNA-Cy5 in the same sample because of the spectral overlap between the 2 dyes.

The siRNAs that were used were the following: a) siRNAs targeting the mRNA for a-synuclein, pooled into a single solution (*SNCA*): s13204, s13205, s13206 (#4427037, Silencer Select predesigned siRNAs, ThermoFisher), b) negative control (scrambled) siRNA: (#4390843, ThermoFisher), c) Cy5 tagged siRNA (siRNA-cy5): MISSION® siRNA Fluorescent Universal Negative Control #2, Cyanine 5 (#SIC006-5X1NMOL, Merck AG).

In the experiments where the effect of certain compounds on the cell-to-cell transfer of a-synuclein was assessed, 6 well plates were first coated with PDL (Poly-D-lysine hydrobromide, average mol wt 30,000-70,000, #P7280-5mg, ThermoFisher) before seeding the cells. 24h after transfection, the medium was changed to normal growing medium and cells were treated with the following compounds and concentrations: Alpha-amanitin (#A2263-1MG, Merck AG): 1, 2.5, 5, 10, 20ug/ml; cdk1 inhibitor (#217695-5MG, Merck AG): 0.5, 1, 2, 3, 5.8uM; N,N,N′,N′-Tetrakis(2-pyridylmethyl)ethylenediamine: 1, 2, 3, 4uM (#P4413-50MG, Merck AG). The appropriate vehicle-only controls were also included: Alpha-Amanitin: water; cdk1 inhibitor: DMSO; N,N,N′,N′-Tetrakis(2-pyridylmethyl)ethylenediamine: ethanol. On the 5^th^ day after transfection, the cells were collected as described above. Hoechst and live dead stainings were also added.

Right before analysis through flow cytometry, the samples were filtered through a 35um cell strainer (#352235, Corning). The samples were analyzed on a BD Fortessa instrument (BD Biosciences). The following lasers and filters were used for the respective fluorophores: GFP: 488nm laser, 530/30nm emission filter; RFP: 561nm laser, 610/20nm emission filter; Hoechst: 405nm laser; 450/50nm emission filter; live dead far red: 640nm laser, 670/14nm emission filter. 100,000 events were recorded per sample. Compensations and voltages were adjusted based on the single color controls.

For the analysis of the flow cytometry data, single cells were gated using the FSC-A/FSC-H plot. The single color controls were used to set the gatings for the various fluorophores. The spreading ratio was calculated as follows: %RFP+GFP-cells/%RFP+GFP+ cells. Where applicable, the percentage of single cells that were positive for siRNA-Cy5 or Hoechst were also calculated. The number of cells that were dead was determined using the live dead histogram.

### Flow cytometry assays to evaluate the mitochondrial membrane potential and the lysosomal-autophagy axis

To quantify mitochondrial membrane potential, we used the MitoTracker Red CMXRos dye (#M7512, ThermoFisher) and followed a similar procedure to a previously published experiment (Xiao et al., 2016). The stock vial contained 50ug of dye and was reconstituted in 94ul DMSO to a concentration of 1mM. The following concentrations were assayed in tissue culture and used for the dose-response experiment: 25, 50, 75, 100, 150, 200nM. 150nM were used for all other experiments.

To assess the lysosomal-autophagy axis, we used the LysoTracker Green DND-2 dye (#L7526, ThermoFisher) (Chikte et al., 2014). This dye is delivered by the company in reconstituted form, with a stock concentration of 1mM. The following concentrations were assayed in tissue culture and used for the dose-response experiment: 10, 20, 30, 50, 75, 150nM. 150nM were used for all other experiments.

Chloroquine diphosphate (#C6628, Sigma) was used as a positive control for the LysoTracker assay. The stock concentration was 25mM, in water. The following concentrations were assayed in tissue culture and used for the dose-response experiment: 25, 50, 100, 200uM. The concentration used in all other experiments was 50uM.

Carbonyl cyanide m-chlorophenyl hydrazine (CCCP) (#ab141229, abcam) was used as a positive control for the MitoTracker assay. The stock concentration was 10mM in DMSO. The following concentrations were assayed in tissue culture and used for the dose-response experiment: 5, 10, 20, 40uM. The concentration used in all other experiments was 20uM. As a vehicle-only control, DMSO was used.

Other compounds used in those 2 assays were the following:

Rapamycin (#R8781-200UL, Sigma): The stock concentration was 2.5mg/ml in DMSO. The following concentrations were assayed in tissue culture and used for the dose-response experiment: 50, 100, 200, 400nM. The concentration used in all other experiments was 200nM. As a vehicle-only control, DMSO was used.

Torin 1 (#Cay10997-50, Cayman chemicals): The stock concentration was 2.5mM in DMSO. The following concentrations were assayed in tissue culture and used for the dose-response experiment: 50, 100, 200, 400nM. The concentration used in all other experiments was 100nM. As a vehicle-only control, DMSO was used.

Bafilomycin A1 (#B1793-10UG, Sigma): The stock concentration was 4uM in DMSO. The following concentrations were assayed in tissue culture and used for the dose-response experiment: 10, 20, 40, 80nM. The concentration used in all other experiments was 40nM. As a vehicle-only control, DMSO was used.

For the assays in 24 well format, the HEK-aSyn cells were plated and 48h later transfected (if applicable), as described above. 3 days after transfection, they were incubated in the concentrations of Mitotracker or LysoTracker indicated above. The dyes were diluted in the following medium: pure OMEM without phenol+10%FBS+1% PenStrep. Right after, the cells were treated with the compounds at the concentrations previously indicated (as applicable). The cells were then incubated at 37 degrees for 3h.

The cells were collected for analysis through flow cytometry, as previously described. The appropriate single color controls were also included.

For the assays in 96 well plate format (only applicable for the LysoTracker experiment), 96 well plates were coated with PDL. Those experiments were completed using reverse transfections. The siRNAs from the ThermoFisher Silencer Select library were printed using the ECHO555 acoustic dispenser (Labcyte). 100nl were printed per well for a final concentration of 5nM (that would occur after cell seeding and addition of transfection mix). The stock concentration of the library was 5mM, in distilled and sterile water. Each siRNA was printed in technical duplicates. 3 plates in total were used per batch of the experiment. Empty wells were left for addition of the positive and negative control treatments after cell seeding. The peripheral wells were used only for single color control samples because they are sensitive to temperature gradients and evaporations which could adversely affect the results of the siRNA experiments. The experiment was repeated independently 3 times. The plates were frozen until the reverse transfections took place.

The 96 well plates were defrosted in the fridge for 1h. A solution containing 19.6ul of pure OMEM and 0.36ul of RNAiMax per well was prepared and incubated for 5min at room temperature. 20ul were added per well. The plates were centrifuged at 2000g for 1min and then incubated at room temperature for 20min. The cells were seeded at 870 cells/ul density and 80ul per well were added. 3 days later, the plates were prepared for analysis through flow cytometry. The dyes and compounds were added as described above. 3h later, the medium was aspirated using the multichannel aspirator. 50ul of PBS-/- were added per well. This was aspirated and 10ul of trypsin were added per well, followed by 3min incubation at 37 degrees in the dark. 40ul of the following medium were added per well to inactivate the trypsin: pure OMEM without phenol+10%FBS+1%penstrep. 125ul of PBS-/-, followed by 25ul of 16%PFA were added per well, for a final volume of 200ul per well. The plates were analyzed using the high throughput accessory to the Fortessa instrument (BD Biosciences). The following parameters were used: Standard mode, 5x mix, 0.5 flow rate, 100ul mix volume, 500ul wash volume, 200 mix speed, 30ul sample volume. Each sample was recorded for 1min or 10,000 events, whichever was achieved first.

The following dyes were used for confirmatory purposes: LysoTracker Blue DND-22 (#L7525, ThermoFisher), MitoTracker Deep Red FM (#M22426, Thermofisher).

### Flow cytometry assay to assess the cell cycle progression

A previously described method was followed for the cell cycle assay (Eddaoudi et al., 2018). Actinomycin D (#11805017, ThermoFisher), a transcription inhibitor, was included as a positive control. The stock concentration was 1mg/ml in DMSO. The following concentrations were assayed in tissue culture: 2.5, 5, 10ug/ml. 10ug/ml was used in all experiments. DMSO was used as a vehicle-only control. After addition of the positive control in predetermined wells, the cells were incubated with Hoechst dye diluted 1:2000 in PBS-/- for 45-75min (stock concentration was 10mg/ml). Pyronin Y (#ab146350, abcam) was then added at 20ug/well and incubated for another 45-75min (stock concentration was 2mg/ml in water).

For experiments completed in 24 well format, the cells were pelleted as previously described, but instead of being fixed in 2%PFA they were resuspended in PBS -/-.

For experiments in 96 well format, the cells were prepared as follows: The medium was aspirated and 50ul of PBS-/-were added per well. The PBS was aspirated and 10ul of trypsin were added per well. After incubation at 37 degrees for 3min, 40ul of the following medium were added per well: pure OMEM without phenol+10%FBS+1% PenStrep. The cells were pipetted to facilitate detachment. 150ul of PBS-/-were added per well. In both versions of this experiment (24 well and 96 well), the cells were not fixed, because fixation affected the fluorescence properties of the dyes. The samples were analyzed using the high throughput accessory to the Fortessa instrument (BD Biosciences). The following parameters were used: Standard mode, 5x mix, 0.5 flow rate, 100ul mix volume, 500ul wash volume, 200 mix speed, 30ul sample volume. Each sample was recorded for 1min or 10,000 events, whichever was achieved first.

### High throughput screen (HTS)

A commercially available library was used (Silencer Select, Thermofisher). This library contains 64,752 siRNAs (0.25nmol per siRNA) targeting 21,584 transcripts (i.e. 3 siRNAs per transcript) that were lyophilized in 384 well ECHO-qualified low dead volume (LDV) plates. The plates were reconstituted in distilled, sterile water to a concentration of 5mM. For the pooled version of the screen, 3ul of each individual siRNA targeting the same transcript were mixed, with a final concentration of 5mM and a final volume of 9ul.

The high throughput screen was completed as previously described (Pease et al., 2019). PDL-Coated CellCarrier-384 ultra 384 well plates were used as destination plates (#6057508, Perkin Elmer). 20ng of GFP-2a-aSyn-RFP construct that was reconstituted in distilled water were printed per well. Afterwards, the library siRNAs were printed using the ECHO555 acoustic dispenser (Labcyte) that was controlled through the Echo Tempo software (Labcyte). 30nl of each siRNA (pooled or singlet, depending on the experiment) were dispensed per well, in technical duplicates spread over multiple plates. The distribution of the siRNAs in the destination plates was determined through a computer algorithm, with each technical duplicate printed on a different plate at a different position. Plates were printed in batches of 8. 44 positive (3 pooled siRNAs targeting *SNCA*) and 44 negative controls (scrambled siRNA) were included per plate, at a final concentration of 5nM. The controls were homogeneously distributed across plates, for quality control purposes. The peripheral wells in each plate (first and last row, first and last column) were excluded from analyses because they are sensitive to evaporation and temperature gradients and therefore the readout for the library siRNAs could have been adversely affected. A total of 166 destination plates were assayed in the primary, pooled screen. The plates were frozen until reverse transfections took place.

The plates were processed and analyzed in batches of 8. The plates were defrosted at 4 degrees for 1h before cell seeding. Reverse transfections were used. First, a mix containing the following reagents and volumes per well was prepared: 0.09ul of RNAiMax plus 4.91ul of pure OMEM without phenol. The mix was incubated at room temperature for 5min before adding 5ul per well. The plates were centrifuged at 2000g for 1min and incubated at room temperature for 20min. The HEK-aSyn cells were trypsinized and seeded at a density of 870 cells/ul. 23ul of cells were added per well. The following medium was used: pure OMEM without phenol plus 10%FBS plus 1%penstrep. The plated cells were then kept in the incubator at 37 degrees for 5 days. To reduce the formation of temperature gradients, the plates were rotated manually in the incubator 1-2 times per day. They were rotated at 180 degrees across their longitudinal axis, along with change in position within the same shelf. The plates were spread out on the same shelf and not stacked.

On the 5^th^ day after seeding, the plates were fixed. The medium was aspirated from the wells, 20ul of 2% PFA were added per well, the plates were incubated for 10-15min at room temperature in the dark, the PFA was aspirated, and 20ul of PBS -/- were added per well. The plates were wrapped in foil and stored at 4 degrees until imaging.

The GE IN Cell Analyzer 2500HS (GE Life Sciences) was used for imaging, with the following parameters: 10x objective (air), 2D deconvolution, 1x1 binning, BGOFR_1 polychroic beam splitter, software autofocus per channel, 10% laser power. The following lasers, emission filters and exposures were used: GFP: excitation 475/28nm, emission filter 511.5/23nm, exposure 0.06sec; RFP: excitation 542/27nm, emission 587.5/47nm, exposure 0.07sec. 2 images were acquired per well. Acquisition time per 384 well plate was less than 30min. A subset of plates (48) were imaged on the Opera Phenix (Perkin Elmer) using equivalent parameters.

During data analysis, all peripheral wells were excluded because they are more sensitive to evaporation and external insults. The metric that was used as a readout for the screen was the cell-to-cell transfer ratio of a-synuclein (number of RFP+GFP-units/number of GFP+ units).

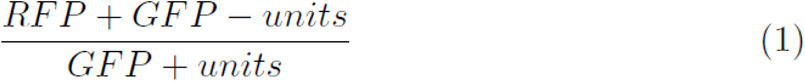

All analyses were done in MATLAB. First, the vignetting artefact, inherent to any microscopy experiment, was eliminated by global background subtraction where each image was divided pixel-by-pixel to the corresponding background. The background image was obtained using an operator size of 15 pixels.

Further pre-processing included the use of a 2D median filter, automatic contrast adjustment and binarization. The binarization was obtained by thresholding the overall image for a fixed value (th=0.1) for all images. This value has been optimized empirically. Once the images for both channels were computed, the difference between the red and green channel was calculated. The cell-to-cell transfer ratio was computed using that difference. A heatmap for each plate was also generated. The code used for data analysis and for the generation of the picklists is available on Github: https://github.com/alecrimi/aSynuclein_siRNA_screen.

Two statistical metrics were used as quality control, the z’-prime factor and the strictly standardized mean difference (SSMD) (Zhang, 2008). Plates with a z’-factor below 0 were repeated, unless indicated otherwise in the results section of this manuscript.

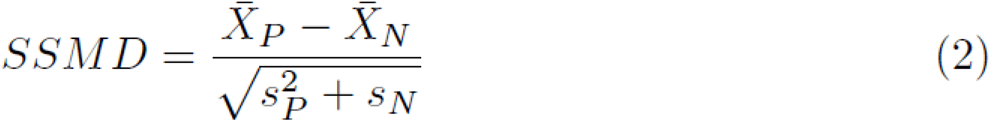

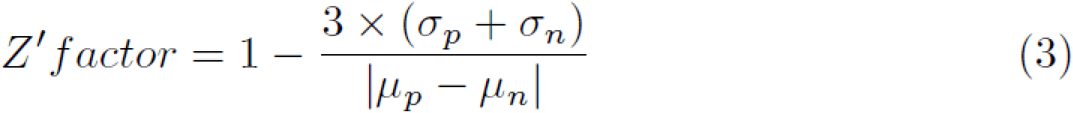

For each gene, the t-test with the corresponding p-value was computed and was used for ranking the hits, after Bonferroni correction for multiple testings. Other metrics that were calculated per gene and used mainly for visual representation of the data were the SSMD and log fold change. All those metrics were calculated relatively to the scrambled controls per batch of 8 384well plates.

### Fluorescence-activated cell sorting (FACS)

Cells were plated in 35mm dishes at a seeding density of 870 cells/ul. 48h later, the cells were transfected with the GFP-2a-aSyn-RFP, E3-RFP or GFP-2a-aSyn constructs. 5 days later, the cells were washed with PBS-/- and trypsinized. Once the cells were detached, wells were pooled and subjected to centrifugation at 1500rpm for 7 min. Pellets were resuspended in FACS buffer (PBS, 2% FCS, 1 mM EDTA) and the samples proceeded to flow cytometry and sorting. Acquisition and sorting of cells was performed using a BD FACS Aria III 5L harnessing a 70 µm nozzle. The drop delay was performed using BD FACS Accudrop beads (BD Biosciences, #345249) according to the manufacturers’ guidelines. Optical configurations were set as follows: a 488 nm Blue and a 561 nm YellowGreen laser were used for optimal excitation of GFP and RFP, respectively. The emission of GFP was recorded using a LP502 mirror in combination with a BP530/30 filter, whereas the RFP was recorded using a LP600 mirror in combination with a BP610/20 filter. Cells were gated for singlets on the FSC-A vs FSC-H plot and for debris exclusion on the FSC-A vs SSC-A plot. RFP+GFP- and RFP+GFP+ populations were determined on the plot for RFP vs GFP. Approximately 1,000,000 cells were collected per tube for the RFP+GFP+ population and 200,000 cells for the RFP+GFP-population.

### Western blot

Cells were plated in 35mm dishes. Where applicable, they were transfected 48h after seeding with the GFP-2a-aSyn-RFP construct. 5 days later they were washed with PBS, trypsinized and pelleted with centrifugation at 1500rpm for 7min. The pellet was frozen until further usage. A pellet from a 35mm dish was lysated in 100ul of lysis buffer consisting of RIPA 1x (#9806S, Cell signaling) and cOmplete protease inhibitors 1x (#11697498001, Merck). For the FACS sorted cells, approximately 200,000 and 1,000,000 cells were lysated in 20ul of lysis buffer for the RFP+GFP- and the RFP+GFP+ populations, respectively. The protein concentration in each sample was measured with the BCA method (#23225, ThermoFisher scientific). A total of 20ug of protein were blotted per sample, after being diluted in 1x NuPage LDS sample buffer (#NP0008, ThermoFisher scientific) and 1x NuPAGE Sample Reducing Agent (#NP0009, ThermoFisher scientific) (final protein concentration 1ug/ul). The samples were run on 4-12% Bis-Tris gels (#NP0321BOX, ThermoFisher scientific) at 120Volt at room temperature with 1x MES running buffer (#NP0002, ThermoFisher scientific). The gels were transferred onto PVDF membranes using the iBlot 2 dry blotting system. After blocking for 2h at room temperature in 5% SureBlock (#SB232010-50G, Lubioscience) in PBS, the membranes were incubated for 2h at room temperature in mouse anti-a-Synuclein antibody, 1:1000 (#610786, BD Bioscience) in 1% SureBlock. Subsequently, the membranes were washed with PBS + 0.1%Tween and incubated with the secondary HRP conjugated goat anti-mouse antibody 1:12000 (#115-035-003, Jackson laboratories) for 1h at room temperature.

After washing, the membranes were treated with Crescendo HRP substrate (#WBLUR0100, Merck) and imaged with the Fusion Solo 7S (witec ag). The membranes were also stained with a mouse anti-actin antibody for 1h at room temperature (#MAB1501R, Merck).

### Reverse transcription quantitative PCR (RT-qPCR)

Cells were seeded in 35mm dishes and transfected as previously described. RNA was extracted from cells from a single 35mm dish using the miRNeasy Mini Kit (#217004, Qiagen) and eluted in 30ul of ddH2O. DNAase I treatment (#79254, Qiagen) was also included during RNA extraction. For FACS-isolated cells, RNA was extracted from 200,000 and from 1,000,000 cells for the RFP+GFP- and for the RFP+GFP+ populations, respectively. cDNA was generated using the High-Capacity cDNA Reverse Transcription Kit (ABI, # 4368814).

qPCR was performed following the relative standard curve method, as previously described (de Calignon et al., 2012). Briefly, five 10-fold serial dilutions of cDNA template from HEK-aSyn cells transiently transfected with the GFP-2a-aSyn-RFP construct were prepared for generation of the standard curve. 50ng of RNA reverse transcribed to cDNA were assayed per sample of interest. Each sample and each concentration of the standard curve were assayed in technical triplicates. No template controls were also included for each primer pair. A primer pair targeting the aSyn-RFP transcript (5’-TGC CTT CTG AGG AAG GGT AT-3’ within a-synuclein, 5’-AAG CGC ATG AAC TCC TTG AT-3’ within RFP) and the *GAPDH* housekeeping gene (5’-AAG GTG AAG GTC GGA GTC AA-3’, 5’-AAT GAA GGG GTC ATT GAT GG-3’) were used. For each sample, 5ul of cDNA were included in a reaction with 10ul of 2x Fast SYBR green master mix (#4385612, Applied Biosystems), 1ul of each of forward and reverse primers (3.3uM each) and 3ul of ddH2O. The samples were analyzed in 96 well plates on a 7500 Fast Real-Time PCR System (Applied Biosystems). Ct values were calculated using the 7500 v2.04 software (Applied Biosystems). Standard curves depicting the Ct values vs the log of the RNA input amount were generated for each of the genes. The RNA amount of each of the 2 transcripts was interpolated based on the standard curve for each of the samples assayed. *aSynRFP* expression was normalized to *GAPDH* expression for each sample, and the fold-difference in expression of *aSynRFP* in the RFP+GFP-sample was calculated relatively to the RFP+GFP+ (calibrator) sample.

### Weighted gene co-expression network analysis (WGCNA)

In order to obtain co-expression models from brain tissue, we used GTEx gene expression V6. For all the 13 brain tissues available, we ran the same pipeline. For each tissue sample dataset, we selected only Ensembl genes expressed above 0.1 Reads per kilobase of transcript, per million mapped reads (RPKM) values at least in 80% of the corresponding samples. Then we corrected for batch effects with ComBat by using CENTER variable. Those residuals were normalised at sample and gene level and then the expression was further corrected for a number of SVA axes while controlling for age, sex and PMI covariates. The resulting gene expression values were regressed for PMI, age, sex and the surrogate variables detected by SVA. These residuals, along with the networks and annotations are accessible, for each tissue at the CoExpGTEx GitHub repository https://github.com/juanbot/CoExpGTEx. The co-expression networks are obtained with the WGCNA R package (Langfelder and Horvath 2008) and an additional refinement step of the clusters, based on the k-means algorith implemented in the CoExpNets R package (Botia et al., 2017). We constructed a set of clusters from gene expression values based on correlation between genes across samples through building a gene expression adjacency matrix (with scale free topology). This was converted into a distance (Euclidean) based matrix that we used to create a dendrogram with the hclust package with default values for the corresponding method. Then we used k-means to refine the clusters obtained from the dendrogram. Network annotations are based on the gProfileR package (Reimand et al. 2007). All these models can be downloaded and used locally but they can also be accessed from a Web interface, the CoExp Web app https://snca.atica.um.es/coexp/Run/Catalog/.

### Expression weighted cell type enrichment (EWCE)

EWCE (https://github.com/NathanSkene/EWCE) (Skene and Grant, 2016) was used to determine whether the 38 hits implicated in regulation of a-synuclein transfer have higher expression within particular brain-related cell types than would be expected by chance. As our input we used 1) the list of 38 hits (which excluded *SNCA*) and 2) specificity values calculated for level 1 cell types from two independent human single-nuclear RNA-sequencing (snRNA-seq) datasets. These datasets included 1) snRNA-seq data from the middle temporal gyrus (Allen Institute for Brain Science; Cell Diversity in the Human Cortex, published 2018-9; https://portal.brain-map.org/atlases-and-data/rnaseq#Previous_Human_Cortex) (Hawrylycz et al., 2012) and 2) massively parallel snRNA-seq with droplet technology (DroNc-seq) datasets from the prefrontal cortex and hippocampus (Habib et al., 2017). For the Allen Institute dataset, the cell-type specificity of each gene (i.e. proportion of total expression of a gene in one cell type compared to all others) was estimated using exonic read count values together with the ‘generate.celltype.data’ function of the EWCE package. Specificity values for the human DroNc-seq data had been previously published by Skene et al (Skene and Grant, 2016). EWCE with the target list was then run with 100,000 bootstrap replicates, each of which was selected such that it had comparable transcript lengths and GC-content to the target list, thus controlling for these biases. Data are displayed as standard deviations from the mean, and any values < 0, which reflect a depletion of expression, are displayed as 0. P-values were corrected for multiple testing using the Benjamini-Hochberg method over all level 1 cell types tested in both studies.

### GWAS data and burden analyses

To investigate the role of the 38 identified genes and the effects of their genetic variation on the risk of PD, we utilized summary statistics from the latest International Parkinson’s Disease Genomics Consortium (IPDGC) genome-wide association study (GWAS) consisting of 37.7K cases, 18.6K UK Biobank proxy-cases, and 1.4M controls, and including 7.8M SNPs (Nalls et al., 2019).

For gene-based burden analyses, IPDGC individual level data containing 21,478 cases and 24,388 controls was used. The genotyping data underwent standard quality control and was imputed using the Haplotype Reference Consortium r1.1 2016 (http://www.haplotype-reference-consortium.org), under default settings with phasing using the EAGLE option, as previously described (Nalls et al., 2019).

Imputed variants with more than 10% missing genotypes were excluded and filtered for imputation quality (RSQ) > 0.8. Analyses were adjusted by sex, 10 principal components to account for population stratification, and dataset to account for possible chip bias. Since the variable age was missing for many patients, it was not included as a covariate in our analyses.

Gene-based burden analyses SKAT, SKAT-O, Madson Browing, Fp, Zeggini and CMC were performed by using RVTESTS (Zhan et al., 2016) to assess the cumulative effect of multiple variants at different minor allele frequency thresholds (≤ 0.05, ≤ 0.03, ≤ 0.01) on the risk for PD according to default parameters.

### Weighted protein-protein network interaction analysis (WPPNIA)

WWPNIA was completed as previously described (Ferrari et al., 2018; Ferrari et al., 2017). Briefly, the direct interactors (first layer nodes) of proteins of interest (seeds), as reported in the literature, were downloaded (June 2019) from the following repositories BioGrid, bhf-ucl, InnateDB, IntAct, MINT, UniProt, and MBInfo, as catalogued in the PSICQUIC platform. The pipeline for the download of interactors is freely available as an online tool (http://www.reading.ac.uk/bioinf/PINOT/PINOT_form.html). All interactors were filtered based on the number of publications in which they were reported and the number of different methods used, keeping interactions with a final score >2. UBC was excluded from downstream analyses as it can bind to a large array of proteins targeted for degradation. Analyses were performed in R and networks were visualized with the Cytoscape 2.8.2 software.

Gene Ontology (GO) terms enrichment analyses (June 2019) was performed in g:Profiler (http://biit.cs.ut.ee/gprofiler/).

### Confocal imaging

Confocal imaging was performed on the Zeiss LSM 880 or 980 with Airyscan instruments. Cells were plated and transfected in 8 well glass bottom chamber slides (#055087, LabTek) and fixed in 2% PFA 5 days later for this experiment. Images were processed in Image J.

### RNA sequencing

Cells of passage comparable to the one used for the HTS were analyzed through RNA sequencing to determine whether the genes detected as hits were indeed expressed.

The libraries were prepared following Illumina TruSeq stranded mRNA protocol. The quality of the initial RNA and the final libraries was determined using an Agilent Fragment Analyzer. The libraries were pooled equimolecularly and sequenced in an Illumina NovaSeq sequencer with a depth of around 20 Mio reads per sample.

FastQC was used for quality control of the reads. Sequencing adaptors were removed with Trimmomatic (Bolger et al., 2014) and reads were trimmed by 5 bases on the 3’ end. The TruSeq universal adaptor sequence was as follows: 5’ AATGATACGGCGACCACCGAGATCTACACTCTTTCCCTACACGACGCTCTTCCGATCT. Reads that were at least 20bp long and had an average phred quality score over 10 were aligned to the human reference transcriptome using STAR v2.5.1 (Dobin et al., 2013) under default parameters for single end reads. The distribution of the reads across the isoforms transcribed was quantified with the R package GenomicRanges (Bioconductor Version 3.0) (Lawrence et al., 2013). Differentially expressed genes were identified with the R package EdgeR (Bioconductor Version 3.0) (Robinson et al., 2010). Genes with at least 10 raw counts in at least half of the samples were retained. The TMM normalized expression metric from EdgeR was used to determine if a gene was expressed or not.

## QUANTIFICATION AND STATISTICAL ANALYSIS

All statistical analyses were performed in Graphpad Prism version 8.

For flow cytometry experiments, experiments were repeated 5-6 times independently, and each independent experiment contained 2 technical replicates per condition. The technical replicates were averaged and the independent experiments were collated for collective assessment of the data, after normalizing the data within each experiment to a common sample, usually the negative control or the first data point of a timecourse or dose response experiment, which was arbitrarily designated as 1 (Kara et al., 2017; Kara et al., 2018). In some cases where the absolute values were important (for example to assess cell viability), the data was not normalized prior to collation of independent experiments. P values below 0.05 were considered significant. In experiments where more than one conditions were compared, the appropriate Bonferroni correction for multiple testing was used. The number of biological replicates conducted for each experiment, statistical test used, normalization process (if applicable), along with method for correction for multiple testing is indicated in the respective figure legend. In all scatter plots included in the paper, mean +/- standard deviation is shown, and each dot represents the mean result from one independent experiment.

The analysis and quantification for the HTS, GWAS, WPPNIA, WGCNA, EWCE are included under the previous section.

## DATA AND CODE AVAILABILITY

The code generated during this study is available on Github https://github.com/alecrimi/aSynuclein_siRNA_screen

## Results

### Development of an assay to study the cell-to-cell transfer of a-synuclein that is amenable to high throughput

To study the cell-to-cell transfer of a-synuclein, we used a genetically encoded reporter. Inspired by a system previously developed to study the spread of tau (Wegmann et al., 2015), we cloned a construct encoding GFP-2a-aSynuclein-RFP under the CMV promoter within the mammalian expression vector pcDNA3.1. As a model system, we used a previously reported HEK QBI cell line stably overexpressing human wild type (WT) a-synuclein (HEK-aSyn line) (Luk et al., 2009). We chose that cell line because we reasoned that endogenous expression of a-synuclein is necessary for the correct assessment of the “permissive templating” hypothesis (Hardy, 2005). Upon transfection of the cell line with the construct, the translated protein is cleaved at the 2a position to produce two daughter proteins, GFP and a-synuclein-RFP. A-synuclein-RFP can then spread to neighboring cells, without concomitant transfer of GFP. The 2 cellular populations (donor and recipient cells of a-synuclein transfer) can, therefore, be identified by their colors: the cells expressing a-synuclein generated through expression from the plasmid are positive for GFP and RFP fluorescence, whereas the cells that received a-synuclein through transfer are positive only for RFP fluorescence (figure 1a). We observed that a-synuclein that underwent transfer did not enter the nucleus of the recipient cell, but instead formed punctate structures that localized in the cytoplasm, primarily within the processes (figure 1b). We then isolated through FACS the cellular population that expressed a-synuclein-RFP through transfection (RFP+GFP+), and the population that had taken up a-synuclein-RFP through cell-to-cell transfer (RFP+GFP-) 5 days after transfection (supplementary figure 1a), and analyzed them through western blot, along with transfected but unsorted cells. Both populations contained the aSyn-RFP fusion protein (expected molecular weight 41kDa), confirming the anticipated cleavage pattern of the fusion protein at the 2a position and indicating that aSyn-RFP is the species transferred from cell-to-cell. To a lesser extent, the transfected cells expressed the uncleaved GFP-2a-aSyn-RFP protein (expected molecular weight 68kDa), consistent with previous reports (Wegmann et al., 2015). This species was absent from the recipient cells (RFP+GFP-), as expected. All populations also contained untagged a-synuclein protein (expected molecular weight 14kDa), consistent with expectations, as the HEK-aSyn line was used for transient transfections. The HEK 293T line that was used as a negative control contained no detectable levels of a-synuclein (figure 1c). The absence of aSyn-RFP transcript in the recipient cells was confirmed through qPCR analysis, providing further evidence that the fusion protein was received through cell-to-cell transfer rather than from aberrant expression of the transfected construct (supplementary figure 1b).

**Figure 1:**
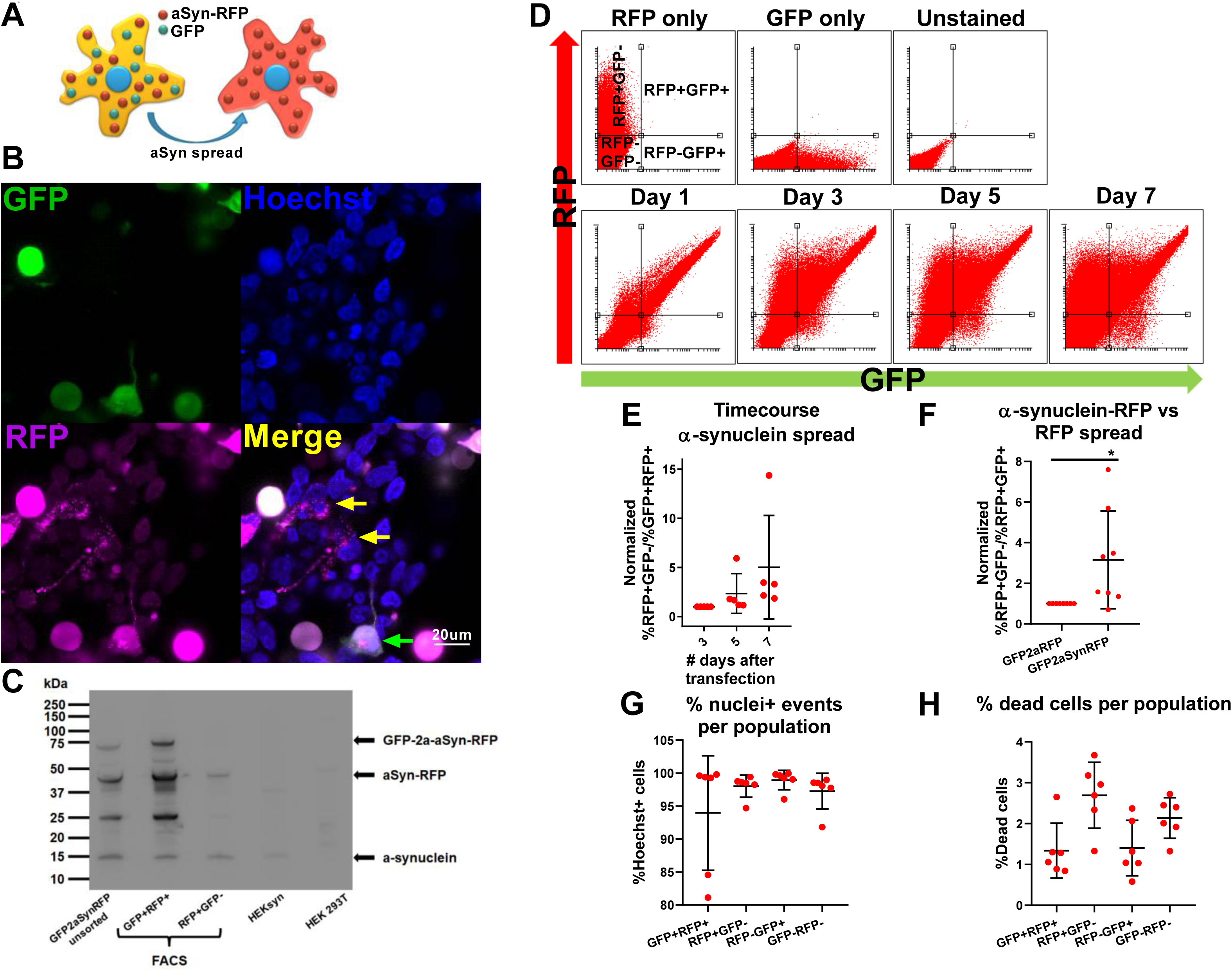
a) Cartoon depicting the rationale behind the a-synuclein cell-to-cell transfer assay used. Cells were transiently transfected with the GFP-2a-aSyn-RFP construct. The cells that express aSyn-RFP through transfection are double positive for GFP and RFP, whereas the cells that have taken up the fusion protein through transfer are positive only for RFP. b) Confocal images showing 1 HEK-aSyn cell that was transfected with the GFP-2a-aSyn-RFP construct and is double positive for GFP and RFP (green arrow), and 2 cells that have received a-synuclein-RFP through transfer and are positive for RFP but negative for GFP (yellow arrows). Scale bar=20um. c) Western blot experiment on bulk HEK-aSyn cells transfected with the GFP-2a-aSyn-RFP construct. FACS-sorted RFP+GFP+ and RFP+GFP-cells were also included. The membrane was probed with an anti-a-synuclein antibody. The expected cleavage pattern at the 2a position of the translated protein was identified, with the aSyn-RFP fusion protein being the major species. The aSyn-RFP fusion protein was also present within the recipient cells (RFP+GFP-), therefore confirming that the RFP fluorescence in those cells originates from the cell-to-cell transfer of the fusion protein. d) Flow cytometry plots corresponding to a representative experiment from panel e. Single color controls and unstained samples are shown to demonstrate how the gates were set. The 4 gates represent the following populations of cells: RFP+GFP+: Transfected cells, RFP+GFP-: Cells receiving a-synuclein through transfer, RFP-GFP-: Untransfected cells, GFP+RFP-: GFP-only cells. e) Timecourse experiment where the HEK-aSyn cells were transfected with the GFP-2a-aSyn-RFP construct, followed by collection at days 3,5 and 7 and analysis through flow cytometry. This experiment was repeated independently 5 times. Data was normalized to the samples collected 3 days after transfection. One way anova with test for trend, a=0.05. f) The transfer propensity of aSyn-RFP was compared to that of the negative control, RFP, using the constructs GFP-2a-aSyn-RFP and GFP-2a-RFP, respectively. The aSyn-RFP fusion protein transferred from cell-to-cell significantly more than RFP alone. One sample t test was used for statistical analysis. *=0.05<p-value<0.01. The experiment was repeated independently 8 times and normalized to the negative control (GFP-2a-RFP) prior to metaanalysis. g) Plot showing the percentage of the events within each of the 4 populations that were positive for Hoechst, indicating that they had a nucleus and were, therefore, intact cells. The majority of the cells contained a nucleus, and the proportions were comparable between the 4 populations. The experiment was repeated independently 5 times and the data were plotted without prior normalization. h) Plot showing the percentage of cells that were dead, as assessed through staining with a far read live dead dye. In all populations the percentage of dead cells was below 4% and was comparable between populations. The experiment was repeated independently 5 times and the data were plotted without prior normalization.

We first validated and characterized the assay using transient transfections in tissue culture, followed by analysis through flow cytometry. In a time course experiment where we collected and analyzed the cells at 3,5 and 7 days after transfection, we found a progressive increase over time of the number of RFP+GFP-cells that received a-synuclein-RFP through transfer (figure 1d,e).

As a negative control, we used a construct encoding GFP-2a-RFP, to check whether the cell-to-cell transfer of the fusion protein a-synuclein-RFP is driven by a-synuclein or by RFP. We collected the cells 5 days after transient transfection with either GFP-2a-aSyn-RFP or GFP-2a-RFP and analyzed them through flow cytometry. We found that, while RFP was capable of transferring between cells, the fusion protein aSynuclein-RFP transferred at a significantly higher ratio (figure 1f). No difference between the transfer ratio of RFP and aSyn-RFP was observed in other cell lines tested (supplementary figure 1c,d,e,f,g), none of which expressed endogenously a-synuclein (supplementary figure 1h). Those results suggest that endogenous expression of a-synuclein could promote the cell-to-cell transfer of the aSyn-RFP fusion protein to levels significantly higher than those of the intercellular transfer of RFP alone.

As a-synuclein in the recipient cells did not diffusely spread within the cytoplasm and nucleus but instead formed punctate structures that apparently localized to the processes, we wanted to confirm that indeed the recipient RFP+GFP-events detected on flow cytometry were intact cells and not debris. To this end, before collecting and fixing the cells for flow cytometry, we stained them with the nuclear dye Hoechst. Analysis through flow cytometry showed that, on average, 98.04% of the RFP+GFP-cells contained a nucleus, a percentage comparable to that seen in the other populations analyzed (figure 1g). Additionally, we incubated the cells with a far red live dead dye before collection and found that all 4 populations of cells had a death ratio of less than 4% (figure 1h). Collectively, those data confirm that the RFP+GFP-events identified through flow cytometry are intact, alive cells.

### Genome-wide siRNA screen for the identification of genetic modifiers of a-synuclein spread

To establish a high throughput screen, we first determined the optimal concentration and incubation time of the siRNA. As a positive control, we used three pooled siRNAs against a-synuclein, each targeting a different region of the mRNA, with a final concentration of 5nM, and co-transfected them with the GFP-2a-aSyn-RFP construct. In a separate experiment, we co-transfected the construct with an siRNA tagged with a far red fluorophore (siRNA-Cy5). Analysis through flow cytometry that was done 5 days later showed that the transfection efficiency of the siRNAs was 92.76%+/-16.12 (figure 2a), with an 80.73%+/-12.02 reduction in aSyn-RFP mean fluorescence intensity (MFI) (figure 2b).

**Figure 2:**
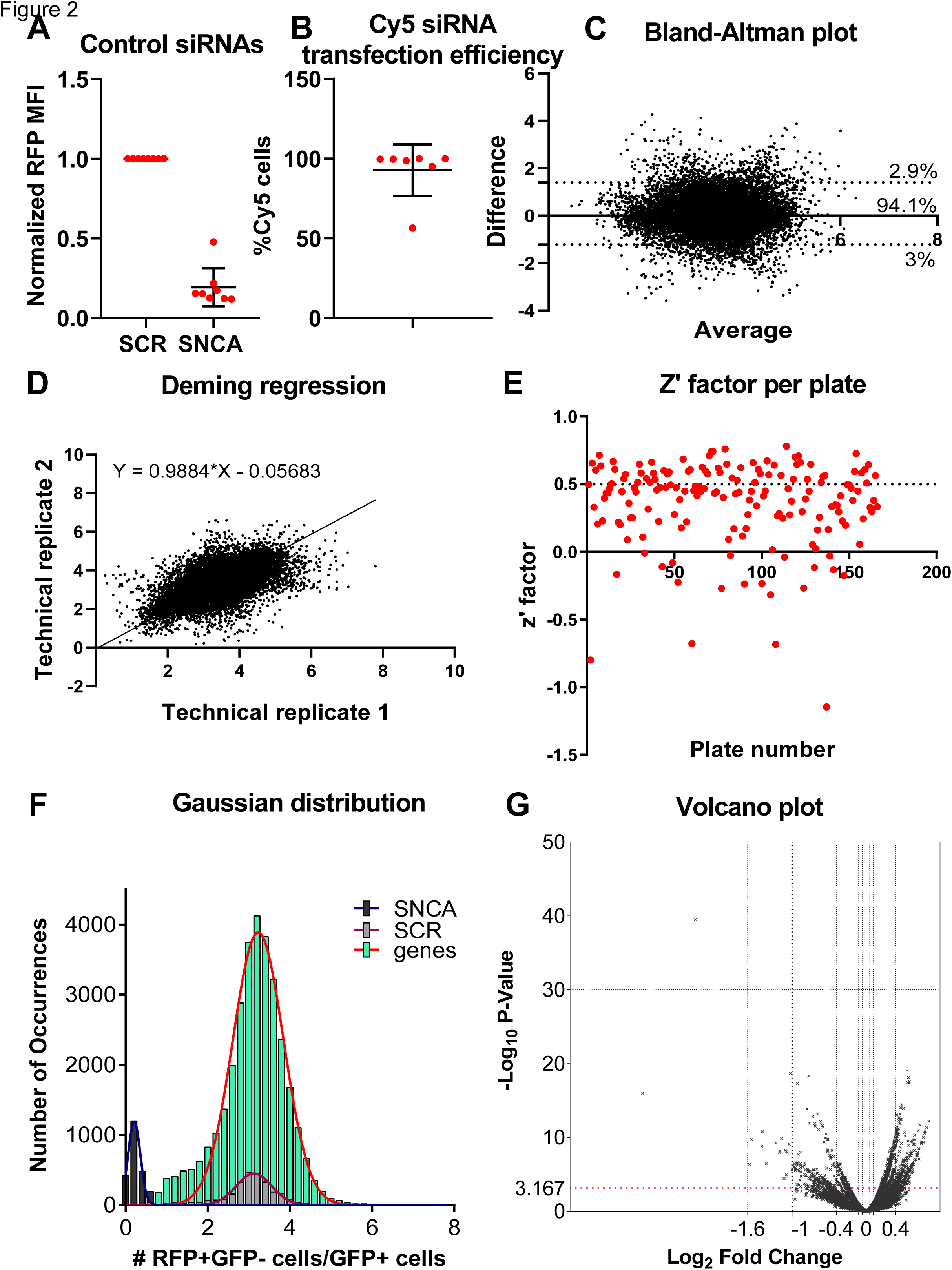
a) The mean fluorescence intensity (MFI) of aSyn-RFP was quantified through flow cytometry after treatment with the positive control siRNA against *SNCA* in comparison to the negative control of scrambled siRNA. There was a 80.73%+/-12.02 (mean+/-SD) reduction of the MFI after siRNA treatment. The experiment was repeated independently 5 times. The data was normalized to the negative control before collation. b) The transfection efficiency of the siRNAs was quantified in an experiment where the GFP-2a-aSyn-RFP construct was co-transfected with an siRNA that was tagged with a Cy5 fluorophore. The transfection efficiency of the siRNA was 92.76%+/-16.12. The experiment was repeated independently 5 times. c) Plot generated by the Bland-Altman test depicting the difference versus the average between the two technical replicates. 94.1% of the genes were within the 95% confidence intervals of the difference between the 2 technical replicates. The plot was symmetrical along both axes indicating that the replicability between technical replicates was not affected when the value of the cell-to-cell transfer ratio increased. The bias metric calculated with this test was 0.096, also indicating that the technical replicates were not diverging. d) Deming regression plot. The slope was significantly different from zero. The slope was almost 1, i.e. a 45 degree angle, indicating a good concordance between the technical replicates. e) Plot depicting the z’-factors per plate for the 166 plates included in the primary screen. Each dot represents one plate. The average z’-factor was 0.36, within the acceptable range. The cutoffs are as follows: <0 unacceptable, 0-0.5 acceptable, >0.5 excellent. f) Histogram of the positive (black), negative (grey) controls and library siRNAs (green) analyzed in the entire primary screen. The separation between the positive and negative controls was good throughout the entire screen, and the library siRNAs overlapped nicely with the scrambled siRNAs. g) Volcano plot depicting the log2(fold change) versus –log10(p-value) for each siRNA.

We next miniaturized the assay into a 384 well plate format that is suitable for an imaging-based readout. Briefly, 5nM of pooled siRNA (3 siRNAs targeting different regions of the same mRNA mixed into one stock solution) were printed per well, in technical duplicates. 44 positive and 44 negative controls were also printed in each plate in a predetermined pattern (supplementary figure 2a). HEK-aSyn cells were seeded and transfected using reverse transfections. The plates were fixed and imaged 5 days after plating.

As part of the quality control procedure, we assessed the plate heatmaps by visual inspection and calculated the z’-factor and SSMD (Zhang, 2008). Some plates in our screen showed plate gradients of variable degrees of severity (supplementary figure 2b). To assess whether this adversely affected the ability of our screen to detect genes modifying the rate of a-synuclein spread by affecting the replicability between technical duplicates, we used two statistical tests, the Bland-Altman test and the Deming regression. The bias metric calculated through the Bland-Altman test was 0.096, and 94.1% of the genes were included within the 95% confidence intervals of the difference between the 2 technical replicates, indicating a good concordance between technical replicates. The Bland-Altman plot was symmetrical and the difference did not skew as the ratio of the two replicates increased (figure 2c). The slope calculated through the Deming regression was significantly different than zero and approached 1, corresponding to a 45 degree angle, also supporting a good concordance between technical replicates (figure 2d). Therefore, the plate gradients do not affect the robustness of the screen and we decided not to repeat the plates with gradients.

The z’-factor and SSMD were used to assess the separation between the populations of positive and negative controls and determine the capability of the screen to detect genes whose knockout either decreases or increases the spread of a-synuclein. The average z’ factor from the entire screen across all 166 plates was 0.36, which is within the acceptable range (figure 2e). 20 plates had a negative z’-factor; however, we decided not to repeat those plates because the second technical replicate of the genes included on them was located on a different plate with positive z’-factor. The SSMD metric was also acceptable for the majority of the plates (supplementary figure 2c). Finally, we plotted the Gaussian distribution/histogram of the positive (*SNCA* siRNAs) and negative (scrambled siRNAs) controls, along with all the 21,584 siRNAs assayed from the library. This graph again confirmed a good separation between the positive and negative controls for the entire screen. It also showed a good overlap between the scrambled controls and the library siRNAs, with the latter population extending on either end of the distribution of the scrambled controls; those extensions correspond to the hits either increasing or decreasing cell-to-cell transfer of a-synuclein (figure 2f).

We quantified the effect of each of 21,584 pools of 3 siRNAs targeting the same gene on the spread of a-synuclein by calculating the t-test p value, SSMD, and logFc. Those metrics were plotted for a visual overview of the results (figure 2g, supplementary figure 2d,e,f). The genes were then ranked by p-value.

*SNCA* was included within the siRNA library and was identified as the top hit of the screen in an unbiased way. Its profound effect on a-synuclein spread was also readily identifiable by visual inspection of the plate heatmaps (supplementary figure 2g). The top 1000 genes as ranked by p-value were selected for a confirmatory secondary screen.

### Secondary and tertiary screens confirm the role of 38 genes in the regulation of cell-to-cell transfer of a-synuclein

A stepwise approach was used to filter the top 1,000 genes (figure 3a). The top 1,000 genes identified through the primary screen were subjected to a series of follow up screens to filter out false positive results. We first completed a screen on the 1,000 genes using single siRNAs per well to assess potential off target effects (secondary screen). Each siRNA was assessed in technical triplicates; therefore, 9 wells were included per gene. Pooled positive and negative controls were also included within the plates, in the same distribution as in the primary screen.

**Figure 3:**
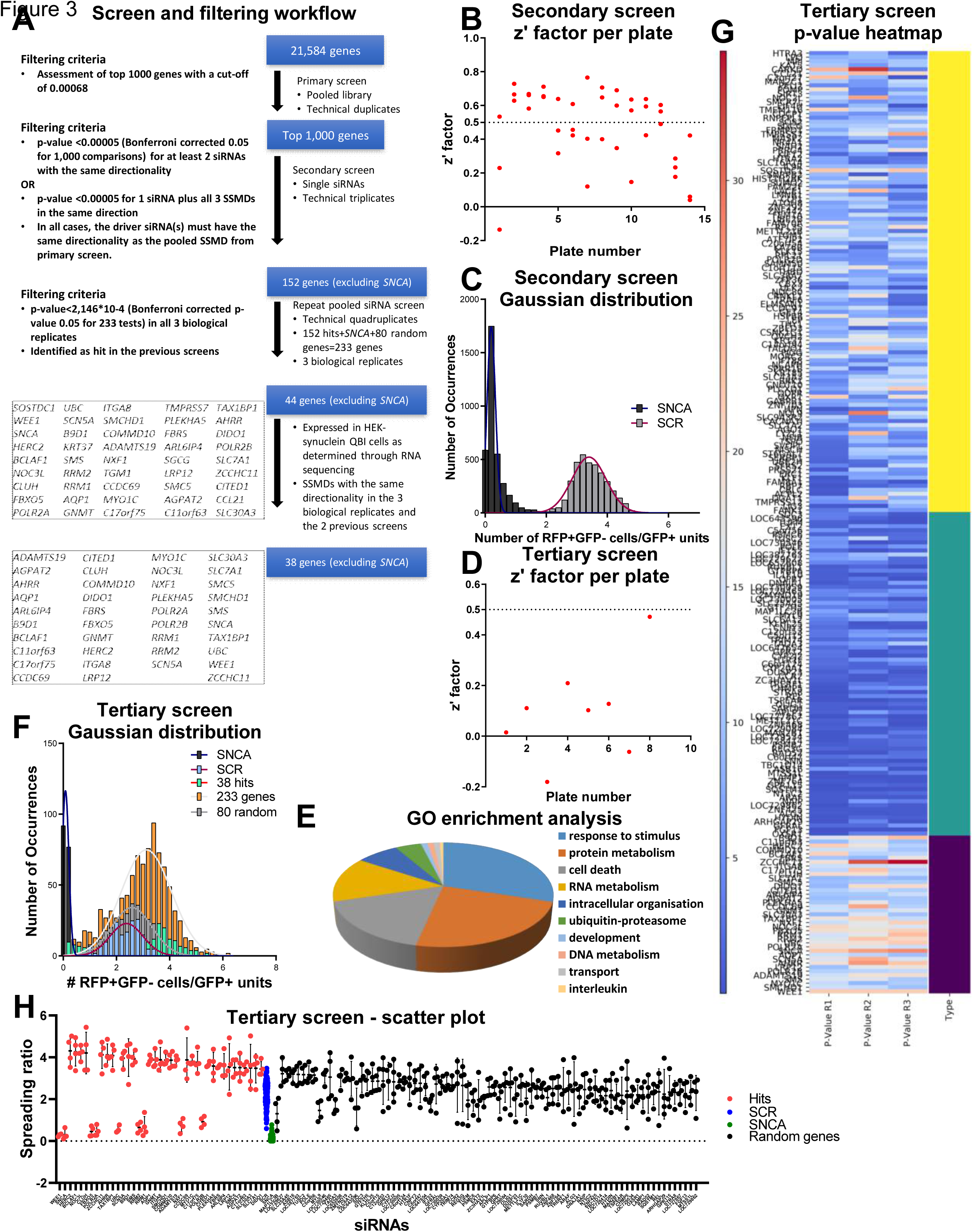
a) Stepwise process used during the high throughput screens and follow up analyses for gene filtering. b) Plot depicting the z’-factors of all plates included in the secondary screen. Only one plate had a negative z’-factor, and the average was 0.48. d) Histogram depicting all the positive and negative controls included in the secondary screen, showing a good separation between them. f) Plot showing the z’-factor per plate for one of the 3 tertiary screens. h) gProfiler analysis for the 38 screens confirmed after the tertiary screens. i) Histogram from one of the 3 tertiary screens showing the distribution of the positive and negative controls, 39 hits, 80 random genes, and all 233 genes included. There is a good separation between the positive and negative controls. The 39 hits cluster on either side of the bars depicting the random genes. j) Heatmaps showing the p-values for each of the 233 genes included in the tertiary screens. The 39 hits form a separate cluster that is identifiable through visual inspection. The 80 random genes are indistinguishable from the 114 hits from the primary screen that did not pass the cutoffs. Color code: purple=39 hits, green= 114/152 filtered out genes, yellow= random genes.

As a quality control assessment, we plotted the z’-factors and SSMDs per plate. The average z’-factor was 0.48 and only one plate had a negative value (figure 3b). The SSMDs were also largely within the good or excellent range (supplementary figure 3a). The histogram for the positive and negative controls for the entire screen showed a good separation between the 2 populations (figure 3c).

The criteria that needed to be fulfilled for a gene to be carried on to the next screening stage were the following:

a. at least 2 of the 3 different siRNAs targeting the same gene should have a p-value smaller than 5*10-5 (Bonferroni corrected p-value of 0.05 for 1000 tests) and the same directionality of effect as determined by the SSMDs, or
b. one of the 3 siRNAs should have a p-value smaller than 5*10-5, plus all 3 different siRNAs targeting the same gene should have the same directionality of effect, as determined by the SSMD values. In any case, the driver siRNA(s) must have the same directionality as the pooled siRNAs from the primary screen.

A total of 152 genes fulfilled the aforementioned criteria and were carried on to the next stage (supplementary table 1) (supplementary figure 3b).

Those 152 genes were then included in a tertiary screen, along with 80 random genes and *SNCA* (supplementary table 2). The pooled version of the library was used (3 siRNAs targeting different regions of the same mRNA, pooled within the same well). Each gene was assessed in technical quadruplicates.

The tertiary screen was repeated 3 times independently. For a gene to be included in the final list of hits, it had to have a p-value smaller than 2.146*10-4 (Bonferroni corrected p-value of 0.05 for 233 tests) in all 3 screens.

For quality control, we again used the z’-factor and SSMD that were calculated per plate (figure 3d) (supplementary figure 3c). 44 genes were confirmed through the 3 tertiary screens, one of which was *SNCA* (figure 3a). Of note, none of the 80 random genes passed the p-value cutoffs specified above. Those results were intersected with the results from RNA sequencing performed on the HEK-aSyn cell line. 39 genes (including *SNCA*) were expressed, and this was the final list of hits (figure 3a). The GO enrichment results for the 38 genes (excluding *SNCA*) are shown in figure 3e. The most common processes seen are response to stimulus (29.8%), protein metabolism (23.6%) and cell death (16.9%).

The separation between the 39 hits and the random genes, and between the 39 hits and the 114 genes from the original list of 152 that did not make it into the final list, is apparent through visual inspection of the respective histograms (figure 3f). In addition, the 39 hits formed a separate cluster as assessed through the heatmaps for p-values and SSMDs (figure 3g) (supplementary figure 3d). Finally, the separation between the hits and the random genes was also apparent in the scatter plot (figure 3h).

The cells were plated at high confluency to increase efficiency of cell-to-cell transfer of a-synuclein through cell-to-cell contact and through increased concentration of secreted a-synuclein in the surrounding microenvironment of the candidate recipient cells. The cells were then incubated for 5 days for adequate cell-to-cell transfer to occur. Therefore, the cells were highly confluent at the time of imaging, which did not permit individual cell segmentation during analysis. For this reason, we quantified the aSyn-RFP “units” transferred from cell-to-cell (RFP+GFP-/GFP+) instead of the number of cells receiving the aSyn-RFP fusion protein through transfer; we then confirmed those quantifications through flow cytometric single cell analysis of the 39 hits plus the positive and negative controls. Those samples were prepared by mechanical dissociation of the fixed cells from the 384 well plates from one of the tertiary screen experiments, followed by pooling of the quadruplicates in one sample, in order to provide enough cell numbers and sample volume. Of note, only directionality of effect, and not effect size, could be confirmed through this technique because it had not undergone rigorous optimization and miniaturization as the imaging-based HTS had. 82% of hits had consistent directionality with the imaging-based screen. 7 hits were inconsistent (supplementary figure 3e).

As the screen was done using a construct encoding for a fusion protein aSyn-RFP, we counter-screened the hits against RFP alone. Not surprisingly, since 1) we know that a small amount of RFP alone can “transfer” between cells (figure 1f) and 2) we are using a nonbiased screen that detects genes that impact a process that involves both uptake and subsequent stability and cytoplasmic localization of extracellular proteins, some of the 39 hits also impact the transfer of RFP (supplementary figure 3f, supplementary table 3). We interpret this to mean that the biology underlying the observed aSyn-RFP uptake effect for these hits is based on an impact on cell processes important for uptake and cytoplasmic localization of extracellular proteins.

### Pharmacological validation supports the role of *SLC30A3*, *ADAMTS19*, *POLR2A*, *POLR2B* and *WEE1* as regulators of the spread of a-synuclein

We sought to confirm a subset of hits through an alternative method. For this purpose, we selected compounds that have been previously reported to target either the product of some of our hits directly, or one of the pathways mediated by those genes. We transfected HEK-aSyn cells with the GFP-2a-aSyn-RFP construct and the following day treated them with various concentrations of the aforementioned compounds, including a vehicle-only control, to generate the dose-response curves. After 5 days, the cells were collected and stained with a far red live dead staining, followed by analysis through flow cytometry.

The zinc chelator N,N,N′,N′-Tetrakis(2-pyridylmethyl)ethylenediamine, treatment which is likely analogous to silencing *ADAMTS19* (a zinc-dependent metalloproteinase) and *SLC30A3* (a protein that mediates accumulation of zinc in synaptic vesicles), showed a significant dose-dependent effect, with the highest concentrations resulting in an increase in cell-to-cell transfer of a-synuclein (figure 4a) and toxicity levels indistinguishable from those of the vehicle-only control (figure 4b). A similar dose-response was not seen in the vehicle-only control experiment (supplementary figure 4a). The directionality of this effect is consistent with that seen during the high throughput imaging-based screens, where silencing of those genes is also associated with an increased cell-to-cell transfer of a-synuclein.

**Figure 4:**
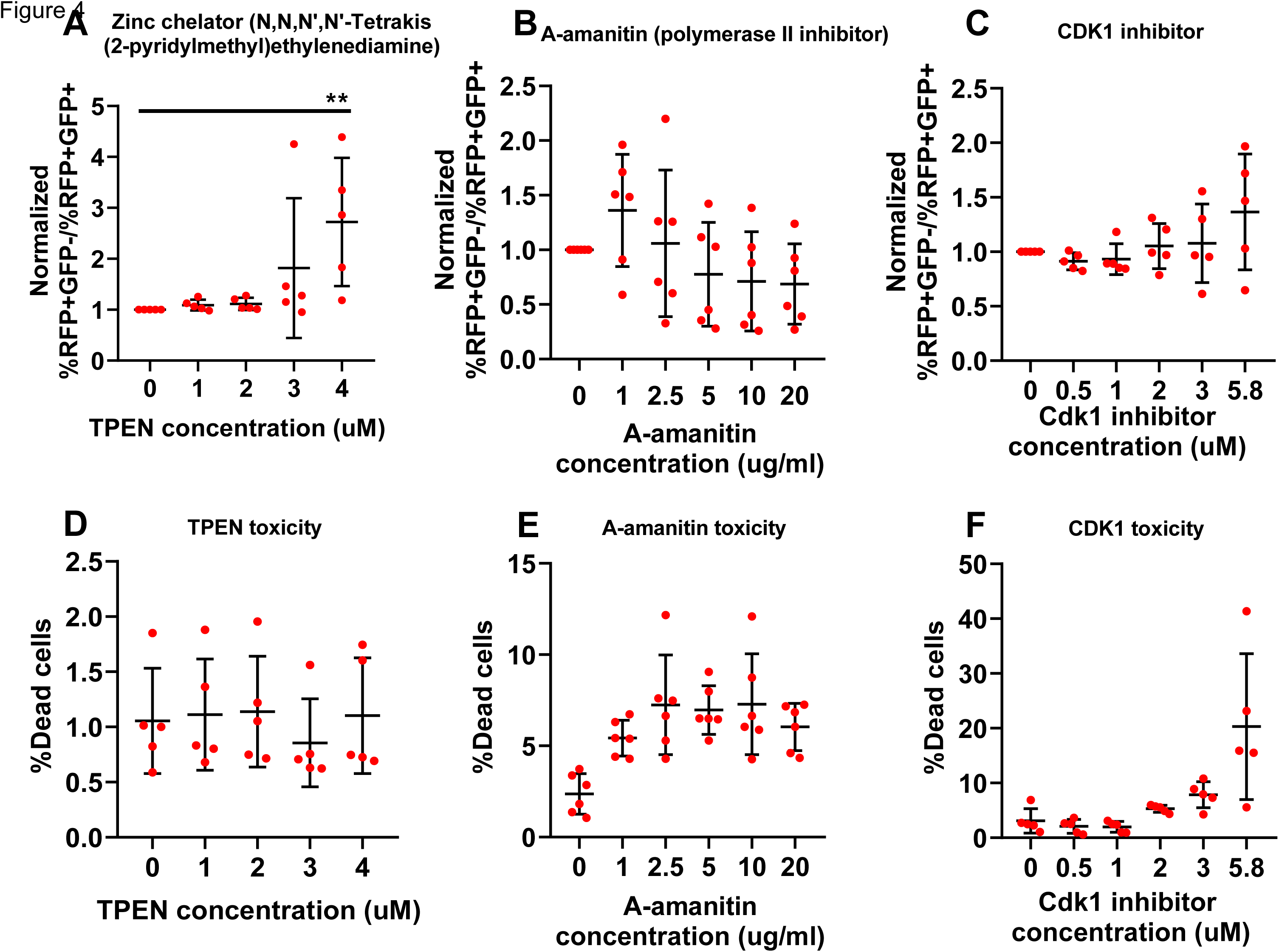
a) Plot showing the effect of the zinc chelator N,N,N′,N′-Tetrakis(2-pyridylmethyl)ethylenediamine (TPEN) on the cell-to-cell transfer of a-synuclein, as assessed through the GFP-2a-aSyn-RFP construct. Treatment with TPEN showed a good dose response effect, with higher concentrations resulting in a greater increase in cell-to-cell transfer. 5 independent experiments were performed and were normalized to the negative control prior to metaanalysis. Statistical analysis was done with one way anova with test for trend. *=0.05<p-value<0.01, **=0.01<p-value<0.001, ***=p-value<0.001. b) Toxicity associated with the compound treatment, as assessed through the far red live dead staining. Toxicity was similar for all concentrations of TPEN used and was not higher than that seen for the vehicle-only control (on average less than 2%). The experiment was repeated independently 5 times. Unnormalized values are shown. c) Dose-response experiment for a-amanitin. The results were not statistically significant, but there was a trend for dose response, with higher concentrations of the compound being associated with a reduction in cell-to-cell transfer of a-synuclein. 6 independent experiments were performed and were normalized to the negative control prior to metaanalysis. Statistical analysis was done with one way anova with test for trend. d) Toxicity measurement in the a-amanitin experiments using the far red live dead stain. Treatment with the compound was associated with a higher toxicity in comparison to the vehicle-only control, which, however, was on average below 10%. The experiment was repeated independently 5 times. Unnormalized values are shown. e) Dose-response experiment for the Cdk1 inhibitor. There was a trend for the spread of a-synuclein to increase as the concentration of the compound increases, though the results were not statistically significant. 5 independent experiments were performed and were normalized to the negative control prior to metaanalysis. Statistical analysis was done with one way anova with test for trend. f) Toxicity measurement in the cdk1 inhibitor experiments using the far red live dead stain. For the 4 lowest concentrations, the toxicity was similar to that of the vehicle-only control and was on average less than 10%. However, this increased to, on average, 20% for the highest concentration of the compound. The experiment was repeated independently 5 times. Unnormalized values are shown.

A-amanitin, which is a selective inhibitor of polymerase II and III, the protein produced by the *POLR2A* and *POLR2B* genes, showed a dose-dependent effect, with higher doses reducing the cell-to-cell transfer of a-synuclein, which, however, was not statistically significant (figure 4c,d). Nevertheless, the directionality of this effect was consistent with what was seen in the imaging-based screen, where silencing of *POLR2A* and *POLR2B* also reduced a-synuclein’s cell-to-cell transfer. Of note, A-amanitin was reconstituted in water, therefore that dose-response effect cannot be attributed to the increasing concentration of the vehicle. *POLR2A* and *POLR2B* were two of the genes whose directionality was inconsistent between the imaging-based screen and the confirmatory analysis through flow cytometry. Therefore, this data supports the robustness of the imaging-based data, in cases of conflict with the confirmatory flow cytometric analysis.

An indolylmethylene-2-indolinone derivative that acts as a Cdk1/cyclin B inhibitor by binding to the ATP pocket on the Cdk1 active site was also assessed regarding its effect on the cell-to-cell transfer of a-synuclein. We expected that treatment with this compound would have the opposite effect of *WEE1* silencing, given that WEE1 is an inhibitor of Cdk1 (Den Haese et al., 1995). Indeed, treatment with the Cdk1 inhibitor showed a dose-dependent effect, with higher concentrations having a trend towards increasing the cell-to-cell transfer of a-synuclein (figure 4e,f), whereas silencing of *WEE1* had an opposite effect. Of note, the dose-response curve for the vehicle (DMSO), showed no trends, with the rates of the cell-to-cell transfer of a-synuclein being similar between all concentrations assayed (supplementary figure 4a).

As a control experiment, we assessed the dose-response effect of various concentrations of DMSO and ethanol (vehicle-only controls) on the cell-to-cell transfer of a-synuclein. The cell-to-cell transfer was similar between all concentrations assessed and no trends were observed (supplementary figure 4a). The toxicity rate was also similar between the various concentrations (supplementary figure 4b).

### Assessment of the lysosomal-autophagy axis, mitochondrial membrane potential and cell cycle progression suggest links of the 39 hits with key functions related to the homeostasis of a-synuclein

We looked for additional evidence supporting the role of our 39 hits in the regulation of a-synuclein transfer, while simultaneously further dissecting their function. To this end, we optimized three assays to assess the mitochondrial membrane potential, the lysosomal-autophagy axis and the cell cycle progression along with RNA quantity per cell cycle phase. We selected those functions because they have been shown to have a crucial role in the homeostasis of a-synuclein (Kara et al., 2013; Ludtmann et al., 2016; Ludtmann and Abramov, 2018; Ludtmann et al., 2018; Wang et al., 2019).

LysoTracker is a dye that has been found to localize to the lysosomes and can therefore be used to indirectly measure lysosomal mass. However, this dye does not exclusively localize to the lysosomes but is also taken up by organelles with an acidic pH. Therefore, the fluorescence intensity of LysoTracker per cell is higher when its lysosomes are larger and/or when the pH within the cell and intracellular organelles is more acidic. MitoTracker is used to assess the mitochondrial membrane potential; cells with depolarized mitochondria fluoresce less intensely when stained with MitoTracker. For the cell cycle assay, the cells that were used as positive controls were first treated with actinomycin D, a global inhibitor of transcription. Thereafter, cells were stained with Hoechst, a dye capable of binding to double stranded nucleic acids. After 45-75min incubation, pyronin Y, a dye that can stain both single and double stranded nucleic acids, was added to the cells and incubated for another 45-75min. Addition of the dyes in that sequence ensured that pyronin Y bound specifically to RNA molecules and, therefore, its intensity would be correlated to the RNA quantity per cell (Eddaoudi et al., 2018).

We first generated dose-response curves for each of the dyes used in those experiments (LysoTracker green, MitoTracker red and pyronin) to confirm that the selected concentration was within the linear range. Those experiments were completed in 24 well plate format. For each concentration of the dye, we included the appropriate positive and negative controls for the respective assay. For the LysoTracker assay, chloroquine was the positive control and untreated the negative control (because chloroquine is reconstituted in water) (figure 5a). For the MitoTracker assay, CCCP was the positive control and DMSO the negative control (figure 5b). For the cell cycle assay, actinomycin D was the positive control and DMSO the negative control (figure 5c). The subcellular distribution of the LysoTracker and MitoTracker dyes was assessed through confocal microscopy (figure 5d).

**Figure 5:**
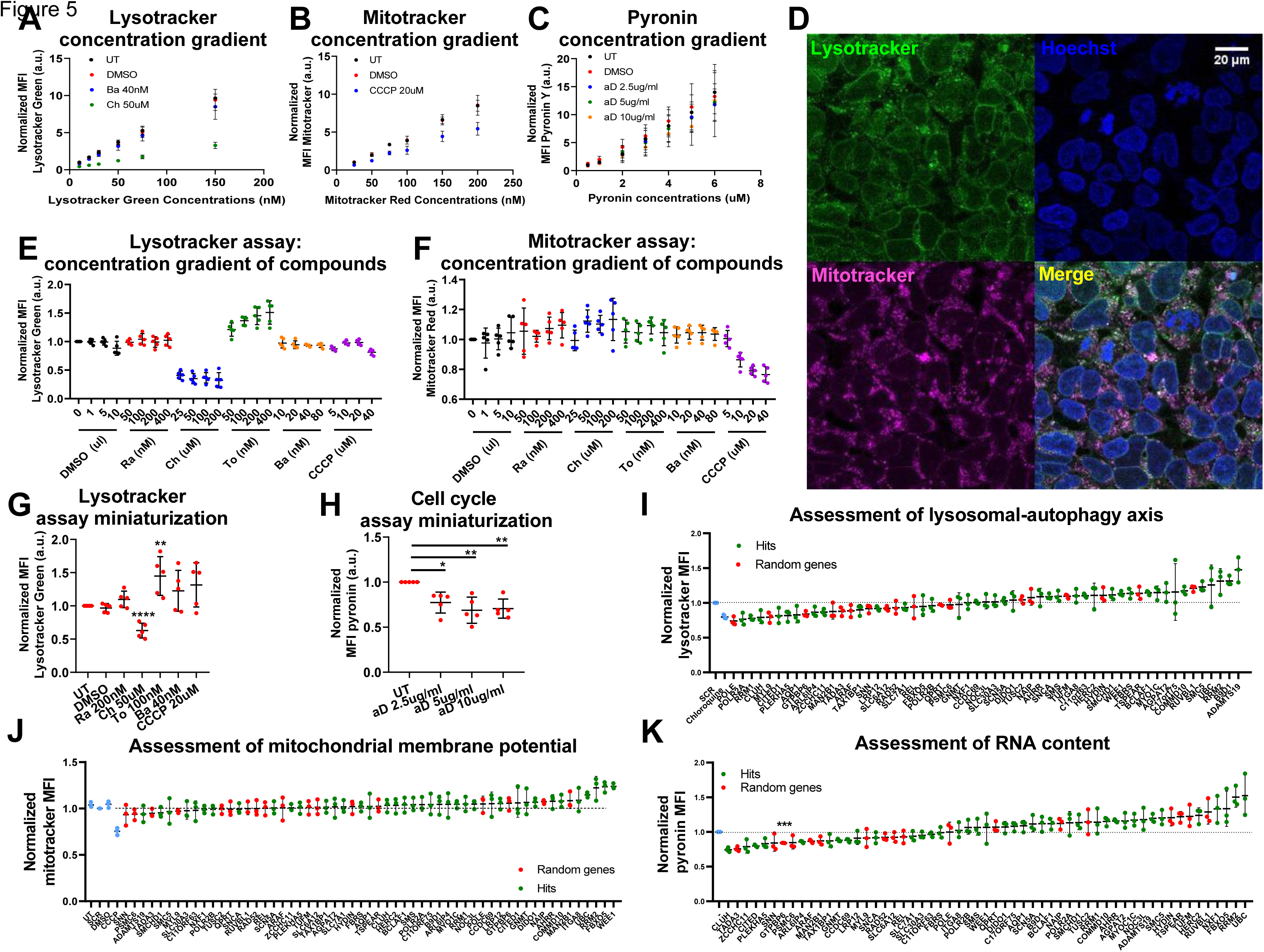
a) Dose-response experiment for the concentration of the LysoTracker dye in the LysoTracker experiment. A linear increase in LysoTracker MFI was observed when the concentration of the dye increased. A good separation between the positive (chloroquine) and the negative (untreated) controls was also seen. The experiment was repeated 5 times independently and the data was internally normalized to the lowest concentration of the dye prior to metaanalysis. b) Dose-response experiment for the concentration of the MitoTracker dye in the MitoTracker experiment. A linear increase of the MitoTracker MFI was seen when the concentration of the dye increased. CCCP caused a reduction in the MitoTracker MFI in relation to the DMSO negative control. The experiment was repeated 5 times independently and the data was internally normalized to the lowest concentration of the dye prior to metaanalysis. c) Dose-response experiment for the concentration of Pyronin Y in the cell cycle experiment. The MFI of pyronin Y increased in a linear way when the concentration of the dye increased. The positive control, actinomycin D clearly separated from the negative control (DMSO-only) and the separation was biggest at the highest concentration. The experiment was repeated 5 times independently and the data was internally normalized to the lowest concentration of the dye prior to metaanalysis. d) Confocal imaging of HEK-aSyn cells stained with the LysoTracker and MitoTracker dyes, depicting the subcellular distribution of those dyes. The concentrations used for LysoTracker and MitoTracker was 150nM. Scale bar=20um. e) Dose-response experiment for the concentration of various compounds in the LysoTracker assay. A good dose response was seen for chloroquine and torin 1, whereas bafilomycin and CCCP had no effect. The experiment was repeated 5 times independently and the data was internally normalized to the lowest concentration of the negative control (DMSO-only) prior to metaanalysis. f) Dose-response experiment for the concentration of various compounds in the MitoTracker assay. A good dose response was seen for CCCP but not for the other compounds. The experiment was repeated 5 times independently and the data was internally normalized to the lowest concentration of the negative control (DMSO-only) prior to metaanalysis. g) Miniaturized version of the LysoTracker assay in 96 well plate format. A significant effect was seen for treatment with chloroquine and torin 1, with the former decreasing and the later increasing the MFI of LysoTracker, similarly to what was seen in the 24 well format of the assay. The experiment was repeated 5 times independently and the data was internally normalized to the untreated sample prior to metaanalysis. The statistical analysis was done with one way anova with Tukey’s correction for multiple testing. *=0.05<p-value<0.01, **=0.01<p-value<0.001, ***=0.001<p-value<0.0001, ****=p-value<0.0001. h) Miniaturized version of the cell cycle assay in 96 well plate format. A significant effect was seen for all concentrations of the positive control, actinomycin D. The experiment was repeated 5 times independently and the data was internally normalized to the untreated sample prior to metaanalysis. The statistical analysis was done with one way anova with Tukey’s correction for multiple testing. *=0.05<p-value<0.01, **=0.01<p-value<0.001, ***=0.001<p-value<0.0001, ****=p-value<0.0001. i) 57 different genes were assayed for LysoTracker MFI (39 hits plus 18 random genes). None of the genes assayed had a statistically significant effect on LysoTracker MFI, but the top 5 upregulators were all hits. 3 independent experiments were performed and normalized to the negative control prior to metaanalysis. Statistical analysis was performed using one sample t tests followed by Bonferroni correction for multiple testings. j) 57 different genes were assayed for MitoTracker MFI (39 hits plus 18 random genes). None of the genes assayed had a statistically significant effect on MitoTracker MFI, but the top 5 genes causing a hyperpolarization in the mitochondria were all hits. 3 independent experiments were performed and normalized to the negative control prior to metaanalysis. Statistical analysis was performed using one sample t tests followed by Bonferroni correction for multiple testings. k) 57 different genes were assayed for Pyronin Y MFI (39 hits plus 18 random genes). None of the hits assayed had a statistically significant effect on the MFI of pyronin Y, but a random gene, *GTPBP6*, caused a significant reduction in the MFI. 3 independent experiments were performed and normalized to the negative control prior to metaanalysis. Statistical analysis was performed using one sample t tests followed by Bonferroni correction for multiple testings. *=0.05<p-value<0.01, **=0.01<p-value<0.001, ***=0.001<p-value<0.0001, ****=p-value<0.0001. Abbreviations: Ch = chloroquine, To = torin, Ra = rapamycin, Ba = bafilomycin, CCCP = Carbonyl cyanide m-chlorophenyl hydrazine, aD = actinomycin D

We also did dose response experiments for the compounds used as positive controls, plus for additional compounds including rapamycin, torin and bafilomycin for the LysoTracker and MitoTracker assays. We saw a good dose response for chloroquine and torin in the LysoTracker assay (figure 5e), and for CCCP in the MitoTracker assay (figure 5f). The rest of the compounds did not have an effect. The ability of the assay to detect the effect of the positive control was confirmed through using different colors of the same dyes, namely LysoTracker blue and MitoTracker far red (supplementary figure 5a,b,c).

The LysoTracker and cell cycle assays were successfully miniaturized into a 96 well format to enable rapid and automatic flow cytometric analysis through the high throughput accessory of the fortessa instrument. The predetermined concentrations of the dyes and the compounds were used in that experiments and gave consistent results with those seen in the 24 well format (figure 5g,h).

The assays were then applied after transfection of HEK-aSyn cells siRNAs against the 38 hits, *SNCA* or 18 random genes. After 3 days of incubation, the dyes were added to the cells and the samples were analyzed through flow cytometry. The appropriate positive and negative controls were also included.

While none of the samples reached statistical significance, the top 5 genes increasing LysoTracker and MitoTracker fluorescence upon silencing were hits from the HTS (figure 5i,j).

For the cell cycle assay, the percentage of the cells in each phase of the cell cycle was quantified (G0G1, S, G2M). The MFI of Pyronin Y per phase of the cell cycle was also determined. Regarding the transition between cell cycle phases, none of the results was statistically significant. However, several of the hits had a tendency to increase cell proliferation, such as *WEE1*, *RRM1*, *RRM2*, *FBXO5*, *AQP1*, *B9D1* and *POLR2A*, as indicated by the fact that a decreased proportion of the cells were in the G0G1 phase and an increased proportion in phases S and G2M (supplementary figure 5d,e,f). This is consistent with what is known about the function of *WEE1* as a regulator of cell cycle progression through modulation of cdk1 (Den Haese et al., 1995). None of the hits significantly modified the RNA quantity within the cell, though there were hits whose silencing was associated with a sub-significant modification of RNA levels (figure 5k).

As the readouts were measured 3 days after transfection, the effect of the siRNAs at that time point was also quantified (supplementary figure g,h).

### Weighted gene co-expression network analysis (WGCNA) shows that most of the hits are included in the same genetic networks as known PD Mendelian and risk genes within the brain

Thanks to genetic studies on familial forms of PD through exome sequencing, linkage analyses and homozygosity mapping, and on sporadic forms of PD through GWAS, we now know numerous Mendelian genes and genetic risk factors for this condition (Singleton and Hardy, 2019). We wanted to assess whether our hits are related to those genes by participating in the same genetic networks. In all bioinformatics analyses, *SNCA* was excluded from the list of hits to avoid introducing a PD-favorable bias; therefore, a total of 38 genes were included in those analyses. For this purpose, we used WGCNA. This analysis is based on the hypothesis that genes that are co-expressed are functionally related.

Through WGCNA, genes are grouped in modules (clusters) within each tissue studied, based on their expression patterns. P-values do not matter regarding the relative importance of each module; rather, the clustering of genes within modules is a qualitative instead of quantitative feature. Those modules can then be queried for the presence of particular genes, followed by the generation of a GO enrichment score and p-value for the most enriched annotated functions. Enrichment for cell type-specific markers can also be assessed (Cahoy et al., 2008; Lein et al., 2007; Miller et al., 2010; Oldham et al., 2008; Winden et al., 2009).

WGCNA was performed using the GTEx version 6 dataset. The modules within each tissue assayed were generated using the recently developed method with an additional k-means clustering step (Botia et al., 2017). We first assessed the ability of WGCNA to correctly identify functional relationships between genes and to correctly annotate those relationships and their enrichment for particular cell type markers. As models for those analyses, we used known Mendelian genes (as catalogued in the Genomics England Panel App) and risk genes for PD (Nalls et al., 2019) (supplementary table 4), and known risk genes for Alzheimer’s disease (AD) (Jansen et al., 2019) and focused exclusively on modules within the brain. The analyses for PD showed that one module passed the p-value threshold after correction for multiple testings. This module was within the substantia nigra, contained 13 PD Mendelian and risk genes, was significantly enriched for GO processes including catechol-containing biosynthetic processes, cell-to-cell signaling, ion transport, and regulation of neurotransmitter levels, and was also enriched for dopaminergic neuron cell markers (supplementary table 5). Similar analyses for the AD genes showed several significant modules within the hypothalamus, pituitary, hippocampus, nucleus accumbens, cortex, cerebellum and caudate nucleus, all of which were enriched for GO functions related to the immune system and for microglial cell markers (supplementary table 6). Therefore, WGCNA is capable of correctly identifying functional relationships between genes and classifying them into modules enriched for functions and cell types experimentally determined to be important in particular disease processes.

We performed a similar analysis for our 38 hits. Assessment of the modules within all brain regions, after application of a cutoff nominal p-value of 0.05, indicated several modules in which the hits cluster. Those modules were enriched for GO functions such as protein modification, regulation of gene expression, cytoskeletal organization, synaptic transmission and organelle organization. Several of the modules were enriched for neuronal cell markers (table 1). Finally, analysis of the GTEX v6 dataset showed that 37/38 (97.37%) of the hits were expressed in human brain, with only *RRM2* not expressed. Altogether, those results support a relevance of the 38 hits to neurons.

**Table 1:** WGCNA for 38 hits identified from the screen. Overlap: number of hit genes relatively to the total size of the module; GO report: GO enrichment analysis results; cell type: enrichment for particular cell type-specific markers in each module; void: not enriched for particular cell markers.

| Network | Module | Overlap | Hits | Fisher | FDR | GO Report | Cell Type |
| --- | --- | --- | --- | --- | --- | --- | --- |
| Cortex | Sky blue | 5/693 | *RRM1*,<br>*C17orf75*,<br>*POLR2B*,<br>*PLEKHA5*,<br>*C11orf63* | 0.0067 | 0.5565 | protein modification by small protein conjugation or removal GO:0070647 (p-value 1.34e-11), mitotic cell cycle GO:0000278 (p-value 1.09e-10), mitotic cell cycle process GO:1903047 (p-value 2.44e-10), protein modification by small protein conjugation GO:0032446 (p-value 5.05e-10), cell cycle GO:0007049 (p-value 1.45e-08) | Neurons |
|  | orange | 4/619 | *NOC3L*,<br>*ZCCHC11*,<br>*SMCHD1*,<br>*SMC5* | 0.0222 | 0.5565 | regulation of RNA metabolic process GO:0051252 (p-value 4.15e-27), regulation of gene expression GO:0010468 (p-value 7.51e-27), regulation of nucleobase-containing compound metabolic process GO:0019219 (p-value 2.61e-26), nucleobase-containing compound biosynthetic process GO:0034654 (p-value 3.52e-26), RNA biosynthetic process GO:0032774 (p-value 2.12e-25) | void |
|  | white | 3/400 | *ITGA8*,<br>*CCDC69*,<br>*MYO1C* | 0.0323 | 0.5565 | vasculature development GO:0072001 (p-value 1.08e-24), cardiovascular system development GO:0031214 (p-value 2.4e-24), anatomical structure morphogenesis GO:0014706 (p-value 8e-24), blood vessel development GO:0003206 (p-value 8.15e-24), circulatory system development GO:0001655 (p-value 2.1e-23) | Endothelial,<br>Microglia |
| Frontal Cortex | brown | 8/2305 | *TAX1BP1*,<br>*SMS*,<br>*COMMD10*<br>*C17orf75*,<br>*POLR2B*,<br>*PLEKHA5*,<br>*LRP12*,<br>*C11orf63* | 0.0407 | 0.5917 | organelle organization GO:0061024 (p-value 3.12e-10), protein ubiquitination GO:0032446 (p-value 1.36e-08), protein ubiquitination involved in ubiquitin-dependent protein catabolic process GO:0043161 (p-value 2.25e-08), modification-dependent macromolecule catabolic process GO:0016236 (p-value 2.47e-07), protein modification by small protein conjugation GO:0070647 (p-value 2.84e-07) | Neuron,<br>Neuron (Pyamidal),<br>Neuron<br>(Interneuron),<br>Neuron<br>(Dopaminergic),<br>In2 |
| Nucleus<br>Accumbens | ivory | 5/768 | *FBXO5*,<br>*TAX1BP1*,<br>*SMS*,<br>*RRM1*,<br>*COMMD10* | 0.0089 | 0.5565 | RNA splicing, via transesterification reactions GO:0000375 (p-value 3.06e-12), RNA splicing GO:0008380 (p-value 7.79e-12), RNA splicing, via transesterification reactions with bulged adenosine as nucleophile GO:0000377 (p-value 9.69e-12), mRNA splicing, via spliceosome GO:0000398 (p-value 9.69e-12), mRNA processing GO:0006397 (p-value 1.23e-08) | Neurons |
|  | sienna3 | 5/789 | *ZCCHC11*,<br>*SMCHD1*,<br>*SMC5*,<br>*SLC7A1*,<br>*DIDO1* | 0.0099 | 0.5565 | regulation of gene expression GO:0010468 (p-value 9.02e-11), regulation of RNA metabolic process GO:0051252 (p-value 6.55e-10), RNA biosynthetic process GO:0032774 (p-value 7.16e-10), nucleic acid-templated transcription GO:0097659 (p-value 2.34e-09), regulation of nucleobase-containing compound metabolic process GO:0019219 (p-value 2.8e-09) | void |
|  | salmon | 3/314 | *WEE1*,<br>*ITGA8*,<br>*CCDC69* | 0.0158 | 0.5565 | axoneme assembly GO:0035082 (p-value 1.8e-10), cilium movement GO:0003341 (p-value 2.31e-10), microtubule bundle formation GO:0001578 (p-value 8.31e-10), cilium assembly GO:0060271 (p-value 4.93e-09), cilium organization GO:0044782 (p-value 9.35e-09) | Ependymal |
| Cerebral Hemisphere | Light cyan1 | 3/338 | *NXF1, FBRS, DIDO1* | 0.0156 | 0.5565 | cytoskeleton organization GO:0007010 (p-value 0.0368) | void |
|  | Floral white | 3/409 | *ZCCHC11, GNMT, PLEKHA5* | 0.0257 | 0.5565 | regulation of cellular macromolecule biosynthetic process GO:2000112 (p-value 2.68e-06), regulation of macromolecule biosynthetic process GO:0010556 (p-value 9.89e-06), regulation of gene expression GO:0010468 (p-value 1.8e-05), regulation of biosynthetic process GO:0009889 (p-value 4.67e-05), regulation of RNA metabolic process GO:0051252 (p-value 5.34e-05) | void |
|  | maroon | 2/171 | *AHRR, CCDC69* | 0.0293 | 0.5565 | chemical synaptic transmission GO:0099536 (p-value 1.62e-05), modulation of synaptic transmission GO:0099537 (p-value 2.33e-05), regulation of system process GO:0032501 (p-value 0.00406), system process GO:0060996 (p-value 0.00596), cell morphogenesis involved in neuron differentiation GO:0031175 (p-value 0.00613) | Neurons, In6 |
| Caudate | blue | 7/1767 | *BCLAF1, FBXO5, TAX1BP1, RRM1,* | 0.0252 | 0.5565 | protein modification by small protein conjugation or removal GO:0070647 (p-value 2.01e-22), protein modification by small protein conjugation GO:0032446 (p-value 4.27e-19), modification-dependent macromolecule catabolic process GO:0043632 (p-value 8.94e-18), ubiquitin-dependent protein catabolic process GO:0006511 (p-value 2.57e-17), modification-dependent protein catabolic process GO:0019941 (p-value 2.66e-17) | Neuron, Neuron (Pyamidal) |
| Hypothalamus | green | 5/1022 | *WEE1, B9D1, CCDC69, PLEKHA5, C11orf63* | 0.0265 | 0.5565 | cilium assembly GO:0070925 (p-value 6.61e-38), cilium organization GO:0030031 (p-value 5.89e-29), axoneme assembly GO:0044458 (p-value 1.61e-27), microtubule bundle formation GO:0000226 (p-value 2.34e-24), cilium movement GO:0007018 (p-value 5.36e-21) | Ependymal |
|  | red | 5/1196 | *ZCCHC11, GNMT, NXF1, FBRS, DIDO1* | 0.0474 | 0.5962 | nucleic acid-templated transcription GO:0097659 (p-value 7.25e-05), transcription, DNA-templated GO:0006351 (p-value 8.58e-05), regulation of gene expression GO:0010468 (p-value 0.000164), regulation of cellular macromolecule biosynthetic process GO:2000112 (p-value 0.000183), regulation of RNA metabolic process GO:0051252 (p-value 0.000205) | void |
| Hippocampus | orangered4 | 3/377 | *CLUH, GNMT, CITED1* | 0.0278 | 0.5565 | void | Neuron (Pyamidal) |
| Amygdala | white | 3/397 | *ITGA8, MYO1C, SLC7A1* | 0.031 | 0.5565 | vasculature development GO:0001944 (p-value 3.71e-24), cardiovascular system development GO:0072358 (p-value 8.01e-24), blood vessel development GO:0001568 (p-value 3.07e-23), angiogenesis GO:0001525 (p-value 5.68e-23), blood vessel morphogenesis GO:0048514 (p-value 3.55e-22) | Endothelial, Microglia |
| Pituitary | salmon4 | 2/182 | *AQP1, GNMT* | 0.0322 | 0.5565 | void | void |
|  | honeydew1 | 2/210 | *TAX1BP1, RRM1* | 0.0417 | 0.5917 | ncRNA metabolic process GO:0034660 (p-value 0.00886), acetyl-CoA biosynthetic process GO:0006085 (p-value 0.0281) | void |
| Substantia Nigra | Dark grey | 3/478 | *AHRR, B9D1, LRP12* | 0.0398 | 0.5917 | chemical synaptic transmission GO:0099536 (p-value 2.04e-16), cell-cell signaling GO:0007267 (p-value 1.71e-12), regulation of synaptic plasticity GO:0060998 (p-value 3.46e-12), behavior GO:0007610 (p-value 5.53e-12), learning or memory GO:0007611 (p-value 7.15e-12) | Neurons, Neuron (Pyamidal), In1 |

We then wanted to assess if our genes are functionally related to known PD Mendelian and risk genes (supplementary table 4). We co-clustered the 38 hits with the PD genes in all brain tissues available in the GTEx v6 dataset in modules with nominal p values <0.05. Using only hits (37 genes) and PD genes that are present in the network, we found that 78.38% of the hits were present within modules that included at least one known PD gene. We then modelled the null distribution of the percentage of genes overlapping with PD genes by generating 100,000 random simulations, each of which containing 37 genes. Based on the fraction of times that we saw a greater % of overlap of the random gene lists with the PD genes than the % overlap observed for the 37 hits, an empirical p-value was generated. We found that the co-clustering of hits with PD genes occurred more frequently than expected by random chance, with an empirical p-value of 0.014 (supplementary figure 6). The hits that are located in the same modules as PD Mendelian and risk genes are listed in supplementary table 7.

**Figure 6:**
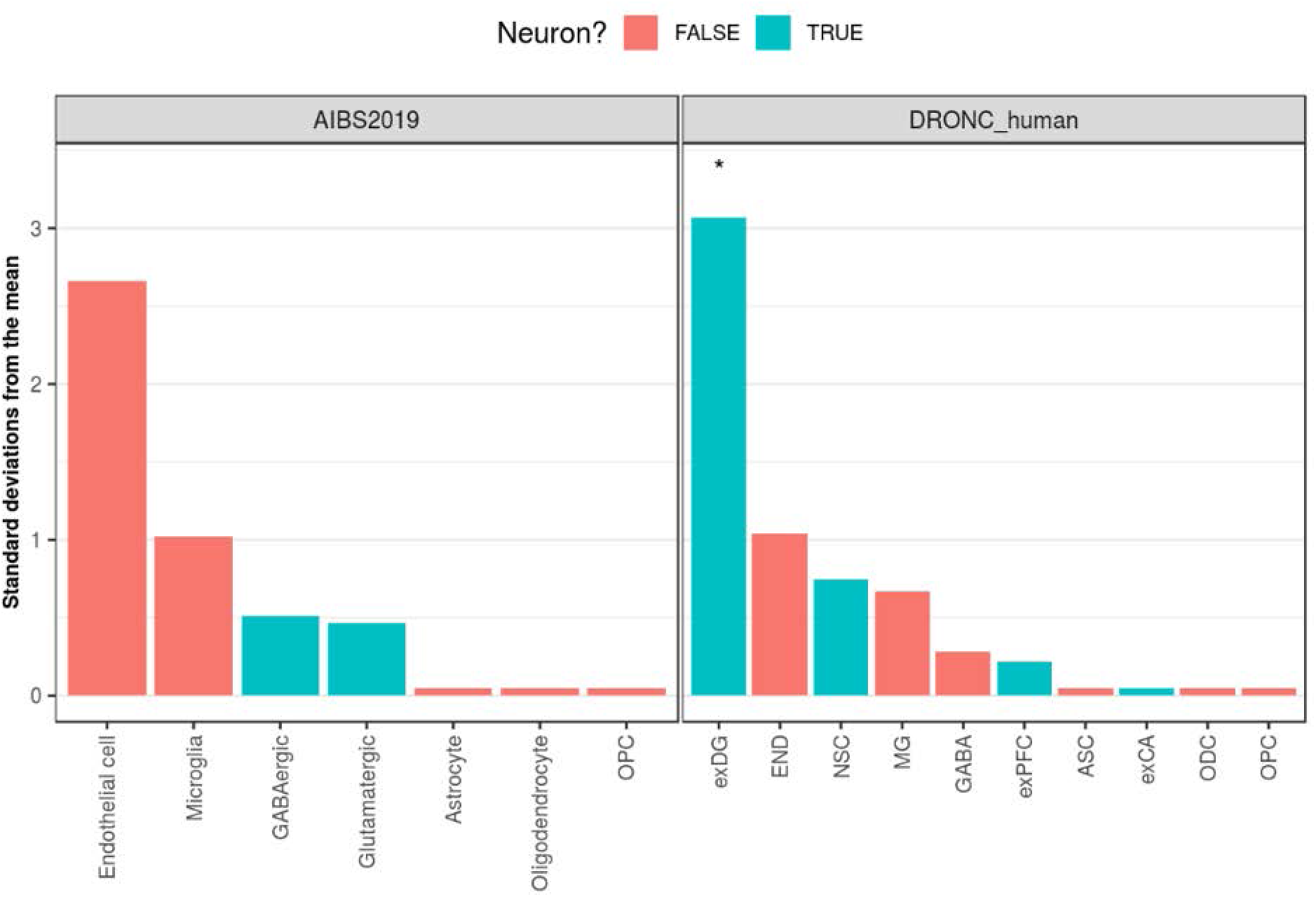
a) Bootstrapping results from EWCE, using data from the AIBS2019 study of the middle temporal gyrus or the DRONC human study of the prefrontal cortex and hippocampus. Standard deviations from the mean denotes the distance (in standard deviations) of the target list from the mean of the bootstrapped samples. Asterisks used to denote results passing FDR < 0.05. Cell-types on the x-axis are colored by whether they are neuronally-related. exPFC=glutamatergic neurons from the PFC, exCA1/3=pyramidal neurons from the Hip CA region, GABA=GABAergic interneurons, exDG=granule neurons from the Hip dentate gyrus region, ASC=astrocytes, NSC=neuronal stem cells, MG=microglia, ODC=oligodendrocytes, OPC=oligodendrocyte precursor cells, NSC=neuronal stem cells, SMC=smooth muscle cells, END= endothelial cells.

Finally, we assessed whether the 38 hits had any cell type-specific characteristics. For this purpose, we performed an expression-weighted cell type analysis (EWCE) (Skene and Grant, 2016). This analysis tests whether the cell type specificity distribution of a certain gene list is greater in one cell type than would be expected if a random gene list of the same length that contained genes with a similar length and GC content was selected. We found the 38 genes to be more highly expressed than expected by chance in granule neurons of the dentate gyrus (FDR-adjusted p-value = 0.045). Given the lack of replicability in other neuronal cell types and the limited availability of single-nuclear RNA-sequencing covering granule neurons of the dentate gyrus (which meant we could not validate the result), we interpret this result with some caution.

### Weighted Protein-protein Interaction Network Analysis (WPPINA) confirms the functional relation between the 38 hits and known PD genes

As an additional way of confirming the functional connection between the 38 hits from the screen and known PD genes, we performed WPPINA. This analysis method was used to mine the literature for previously published protein-protein interactions and generate the networks that are formed by the proteins included in the input list. The networks were generated separately for the 38 hits, the PD Mendelian genes and the PD GWAS risk genes. The network for the 38 hits (hits-network) included 34 seeds (4 hits were excluded as they did not show any interactor surviving our filtering step at the time of the analysis) plus the direct interactors of the seeds, for a total of 615 nodes. The PD Mendelian network included 18 seeds plus their direct interactors, for a total of 545 nodes. Finally, the PD risk gene network included 65 seeds and their direct interactors, total of 902 nodes. The overlap between a) the network of the hits and the PD Mendelian network, and b) the network of the hits and the PD risk gene network was then assessed by determining the communal nodes, which could be either seeds or direct interactors of the seeds. As a negative control, we generated 100,000 random gene lists, each comprising of 615 genes (to simulate the number of nodes in the real hits-network). We then calculated the overlap of these random lists with the real PD Mendelian and risk networks. We drew the distribution curve of random overlaps between the random lists and the (real) PD Mendelian network (real node matches = 79 vs random nodes matches = 16) and between the random lists and the (real) risk network (real node matches = 132 vs random nodes matches = 28) indicating that the real networks intersected more frequently than expected by random chance (supplementary figure 7a and b). A visual representation of the overlap between those networks is depicted in figure 7a,b. The intersection of the networks formed by the hits, the PD Mendelian and the PD risk genes contained 50 genes (figure 7c). GO enrichment analysis of those genes showed that the top 3 most enriched processes were cell signaling (23.9%), cytoskeleton-mediated transport (22.7%) and cell death (18.2%) (figure 7d).

**Figure 7:**
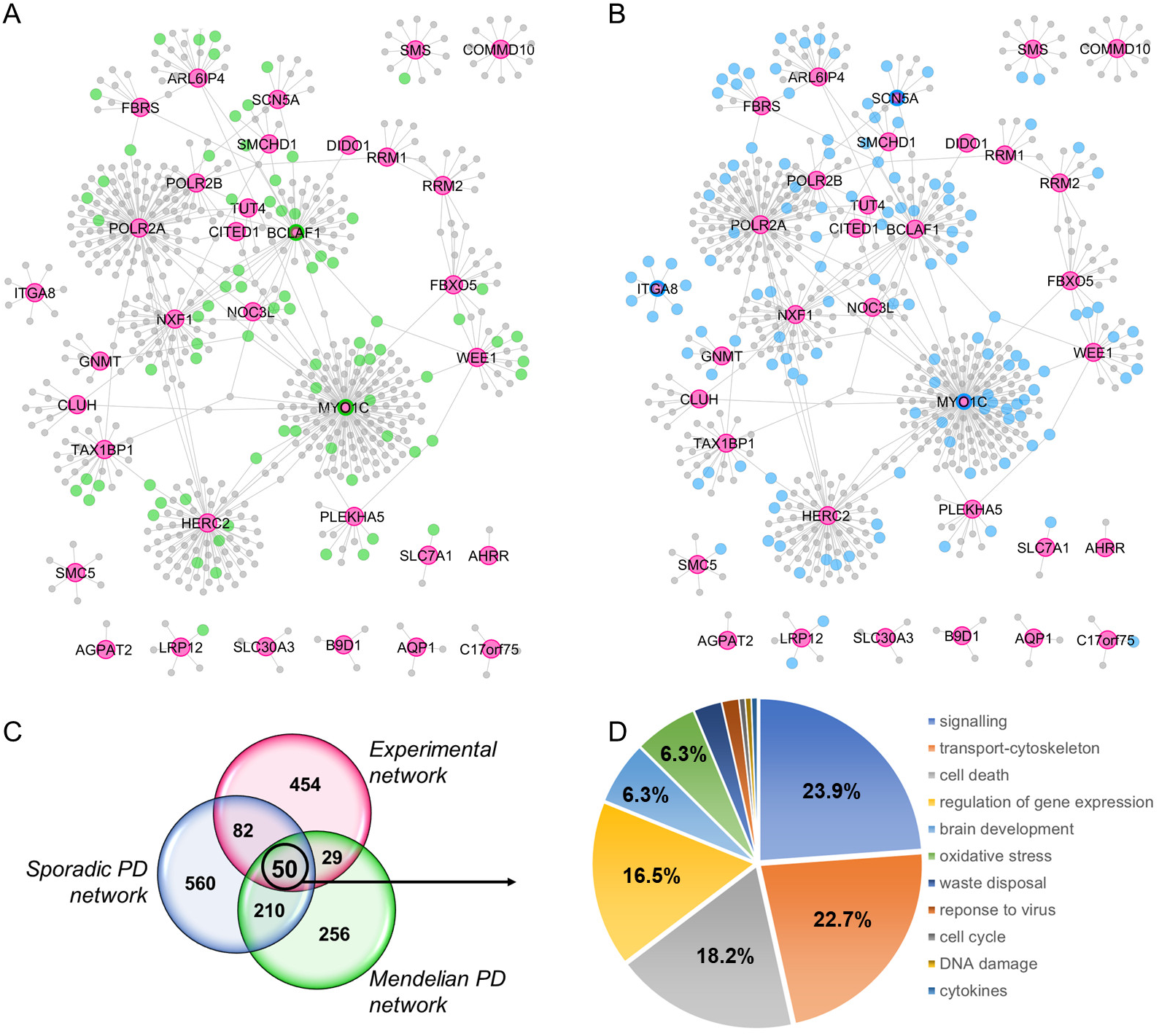
a) 34 seeds from the “experiment” (Supplementary File 8) have been used as seeds for building the “Experimental protein-protein interactions network” or hits network. The network was built after downloading the (first layer) protein interactors of the seeds and by filtering for reproducibility of interactions (see materials and methods). Pink nodes are the seeds of the experimental network while green nodes are those that overlap across the experimental network and the protein interaction network built around the PD Mendelian genes (Supplementary file 9 and materials and methods). b) Pink nodes are the seeds of the experimental network while blue nodes are those that overlap across the experimental network and the protein interaction network built around the risk (sporadic) PD genes (Supplementary file 9 and materials and methods). c) Venn diagram detailing the overlaps across the nodes composing the experimental - Mendelian PD - sporadic PD networks. d) Functional enrichment (for Gene Ontology biological processes) performed with the 50 nodes that are communal across the 3 networks. Gene ontology terms are filtered and grouped into semantic classes by semantic similarity. General semantic classes have been excluded from analysis (the entire enrichment is presented in Supplementary file 10).

### PD patients do not carry an increased burden of rare variants within the 38 hits in comparison to healthy controls

We found that the networks formed by the 38 hits significantly overlap with the networks formed by the PD Mendelian and risk genes. Subsequently, we assessed whether the 38 hits contained risk variants that directly contribute to the causation of PD. To this end, we used the most recent IPDGC GWAS summary statistics including 37.7k PD patients, 18.6k UK Biobank proxy-cases, 1.4M controls, and 7.8M SNPs that identified 90 risk loci contributing to the risk of disease (Nalls et al., 2019). We first compared our 38 hits with the top candidates per locus, as determined by previously undertaken eQTL analysis (Nalls et al., 2019). One gene was in common, *ITGA8*.

We then checked whether the hits overlapped with PD GWAS loci using a 500kb window up and downstream the region, to account for synthetic associations. Out of the tested hits, four overlapped, including *ARL6IP4*, *FBRS*, *POLR2A* and *ITGA8*. We also generated a list containing all the 1,335 genes spanning those regions. Statistical analysis using a Fisher’s exact test showed that the clustering of the 4 hits within GWAS loci did not occur more frequently than expected by random chance.

Finally, we performed burden analyses on imputed genotyping data to check whether the 38 hits contained an increased frequency of rare variants in 21,478 PD patients and 24,388 healthy controls (IPDGC dataset, excluding 23andMe cohort). The burden was computed in 2 different ways: a) within each gene individually, b) across WGCNA modules in which the hits clustered. None of the WGCNA modules showed enrichment after Bonferroni correction for multiple testing. When we looked at the burdens within each gene individually, no genes were significant after Bonferroni multiple testing correction (supplementary table 11).

## Discussion

We performed an imaging-based siRNA screen to identify genes that regulate the cell-to-cell transfer of a-synuclein. We identified 38 novel genes that modulate this process, 9 of which decrease and 29 of which increase the cell-to-cell transfer of a-synuclein upon silencing. Interestingly, silencing the majority of those genes resulted in the increase of a-synuclein spread, suggesting that the cell has mechanisms that act as “brakes” preventing the cell-to-cell transfer of this and possibly also other proteins. Those genes were enriched in functions implicated in the homeostasis and intercellular spread of a-synuclein, such as cell-to-cell signaling, cytoskeletal-based transport and waste disposal. Several of the identified genes have been previously linked to the pathogenesis of parkinsonisms and other neurodegenerative diseases. *NFX1* has been identified through a CRISPR-Cas9 screen as a gene that reduces the levels of dipeptide repeat proteins (DPRs) by lowering the nuclear export of GGGGCCC repeat-containing transcripts upon silencing (Cheng et al., 2019; Hautbergue et al., 2017). That gene was also identified through our screen as a reducer of a-synuclein spread upon silencing, presumably through lowering intracellular levels of a-synuclein, thereby having a consistent directionality of effect as reported in that c9orf72 screen.

Our screen also identified *FBXO5* as a hit that reduces a-synuclein transfer upon silencing. That gene belongs to the same family as *FBXO7*, a gene mutated in complex parkinsonisms (Di Fonzo et al., 2009; Paisan-Ruiz et al., 2010; Zhao et al., 2013) that is involved in mitochondrial homeostasis and in the ubiquitin ligase complex called SCFs (SKP1-cullin-F-box). FBXO5 is also part of the SCF complex (Randle and Laman, 2016), and our MitoTracker assay showed that FBXO5 induces mitochondrial depolarization upon silencing, suggesting a role of that gene in mitochondrial homeostasis as FBXO7 (Burchell et al., 2013).

We found that *SLC30A3* and *ADAMTS19* increase the transfer of a-synuclein when silenced. Both those genes are implicated in the homeostasis of zinc. SLC30A3 has been implicated in the accumulation of zinc within the synaptic vesicles in mice (Cole et al., 1999), whereas ADAMTS19 is a member of the ADAMTS family of zinc-dependent proteases. *ADAMTS4*, a gene belonging to the same family as *ADAMTS19*, has been identified as a candidate risk gene for AD through GWAS (Jansen et al., 2019). We have further shown that zinc chelation results in an increase of a-synuclein transfer, a directionality that is consistent with the effect seen when those 2 genes are silenced. There are several lines of evidence supporting a role of zinc in the pathogenesis of PD. It has been found that the parkinsonism gene *ATP13A2* encodes a zinc pump that facilitates transport of zinc into vesicles (Kong et al., 2014), a similar function as *SLC30A3*. While *ATP13A2* was not identified as a hit through our screen, silencing of that gene resulted in a trend towards increased a-synuclein cell-to-cell transfer. Reduced zinc levels in the serum and plasma have also been found in advanced forms of PD (Du et al., 2017). The exact mechanism with which zinc modulates the cell-to-cell transfer of a-synuclein remains to be elucidated.

*LRP12* and *ITGA8* were also identified as hits from our screen. *LRP12* belongs to the same family as *LRP10*, a gene encoding a low-density lipoprotein receptor-related protein that has been found to carry risk variants in patients with PD (Quadri et al., 2018). Previously published genetic analyses have already implicated *ITGA8* in the pathogenesis of PD. *ITGA8* is a risk gene for PD that was identified in the most recent PD GWAS meta-analysis (Nalls et al., 2019). In addition, Mendelian Randomization-QTL analyses showed a significant association between decreased expression of *ITGA8* and PD risk in brain caudate basal ganglia data (Consortium, 2013) (Nalls et al., 2019); concurrently, our data indicated that reduction in the expression levels of *ITGA8* results in an increase in the cell-to-cell transfer of a-synuclein. Therefore, the directionality of the effect associated to reduction of mRNA levels in both studies is the same. Finally, *ITGA8* downregulation has been found through an unbiased screen to increase the susceptibility of cells to infection with proteinase K-resistant PrP (PrP^sc^) (Imberdis and Harris, 2014). *ITGA8* encodes alpha-8 integrin, a heterodimeric transmembrane receptor that is involved in functions such as cell adhesion, cell signaling and cytoskeletal organization (Humbert et al., 2014), functions that could all conceivably be related to the cell-to-cell transfer of a-synuclein.

Even though one hit, *ITGA8*, is the top candidate for a PD GWAS locus, the burden analyses did not identify an increased frequency of rare variants within the hits in PD patients in comparison to healthy controls. However, it is still possible that our hits are causal but the current analysis methods and technologies were not sufficiently powered to detect such variants. GWAS have been designed to detect common variation related to complex diseases, and rare variants are difficult to impute with accuracy from that data. In addition, structural variation is not captured in GWAS data.

Our screen had some technical limitations. First, we decided to add an RFP tag to a-synuclein to enable its detection through imaging, in order to eliminate extra processing steps for antibody staining. It has been found that addition of a tag to a-synuclein modifies its ability to form aggregates (McLean et al., 2001), therefore possibly also modifying its ability to spread (Gustafsson et al., 2018). However, we have controlled for this limitation by including a negative control, GFP-2a-RFP. We found that, while RFP has the ability to spread, it does so significantly less than the fusion protein a-synuclein-RFP. Second, our 384 well plates sometimes exhibited temperature-related gradients, which could cause poor replicability between technical replicates. However, statistical analyses using the Bland-Altman test and the Deming regression, indicated that replicability between technical replicates is good and, therefore, the plate gradients do not adversely affect our ability to draw robust conclusions from our screen. Third, we had to resort to the quantification of “units” transferred from cell-to-cell instead of cells as a readout of the imaging-based screen. This was because we were unable to segment individual cells due to their high confluency. Nevertheless, the directionality of the effect was confirmed through single cell flow cytometric analysis for 82% of the genes. From the 7 inconsistent genes, we looked with more detail into *POLR2A* and *POLR2B*. The imaging-based screen showed a reduction in spread upon their silencing, while flow cytometry analysis an increase. Pharmacological approximation of their silencing using a-amanitin, a compound that selectively inhibits polymerases II and III, showed a reduction in the spread of a-synuclein, which is consistent with the results of the imaging-based screen.

Conversely, our system also has a significant advantage in that it studied the spread of a-synuclein produced endogenously from the donor, mammalian cells. Our study also did not make any assumptions about the aggregation status of the transferred species of a-synuclein, given that the nature of this species is still unclear (Cremades et al., 2012; Kurnik et al., 2018; Volles and Lansbury, 2003). This presents a considerable advantage over the commonly used system assessing the uptake of a-synuclein preformed fibrils (PFFs) from recombinant protein, and over systems based on primary neurons or mouse models. Systems using PFFs presuppose that the spreading species of a-synuclein is the aggregated, fibrillar form. While data from intracerebral injection of PFFs in mice supports this hypothesis, there is a growing body of data contradicting this notion and supporting that the pathogenic species is the oligomeric form (Cremades et al., 2012; Kurnik et al., 2018; Volles and Lansbury, 2003). In addition, the PFFs are highly artificialized units and it is unclear what, if any, similarities they have to physiologically produced aggregates (Strohaker et al., 2019). Finally, systems using PFFs do not take into account the existence of donor cells as potentially crucial partners in the spread of a-synuclein from cell-to-cell.

A recent study was performed using PFF uptake in primary neuronal and induced pluripotent stem cell (iPSc) cultures to study the effects of known PD Mendelian genes on the uptake of a-synuclein in the form of PFFs (Bieri et al., 2019). Among known PD genes, *LRRK2* was found to significantly reduce a-synuclein uptake upon silencing. However, *LRRK2* was not identified as a hit in our screen, even though the directionality of the effect was the same. There are several possible explanations for the discrepancies between the two studies. First, the model systems used were different, both in terms of the cell type under study and the source and state of the transferred a-synuclein. Second, the total number of genes studied was substantially different; the study by Bieri et al included only a small number of genes, therefore employing less strict p-value cutoffs compared to the ones used in our study. It is possible that if we had only included the same genes and employed the same p-value cut-offs, we would have also identified *LRRK2* as a modifier of a-synuclein transfer.

The 38 genes that regulate the cell-to-cell transfer of a-synuclein participate in genetic and protein networks formed by Mendelian and risk PD genes. The fact that there is a genetic overlap between the phenomenon of a-synuclein spread and the causation of PD indicates that the role of a-synuclein transfer is a crucial part of the pathogenesis of the disease. Those observations support the validity of the “prionoid” hypothesis (Aguzzi et al., 2007; Aguzzi and Rajendran, 2009), in lieu of (or in combination with) alternative explanations for the progression of a-synuclein pathology such as the selective vulnerability hypothesis (Alegre-Abarrategui et al., 2019; Fu et al., 2018; Hardy, 2016).

The data generated by our screen could be consistent with the “omnigenic” model of disease, a recently formulated, intriguing hypothesis regarding the genetic causes of complex diseases such as PD and AD (Boyle et al., 2017; Wray et al., 2018). A few protein-coding genes and their direct regulators (termed ”core genes”) carry highly penetrant mutations inherited in a Mendelian fashion that cause a particular disease. However, GWAS have found that the majority of the known genetic causes of complex disease are common, low effect size variants found primarily within non-coding and regulatory regions. The omnigenic model proposes that any genes expressed within tissues affected by the particular disease process are likely to be functionally linked to the “core” disease genes; any variant within those genes can have a regulatory effect on the “core” genes, and therefore even minimally affect disease pathogenesis. Consequently, genetic variants could be classified into three categories: a) highly penetrant mutations in core genes causing Mendelian forms of the disease, b) lower penetrant risk variants usually within intergenic non-coding, regulatory regions, with sufficient effect size to enable detection through GWAS, and c) variants of variable frequency but very low effect size that are located within housekeeping genes or regulatory regions. While the first 2 types can be identified using conventional genetic approaches, the third category is notoriously difficult to study because of the low effect size of the variants, low penetrance, and requirement of very large case-control cohorts that are exceedingly difficult to assemble and costly to analyze through GWAS and exome sequencing. The data generated through our screen support this model because the identified hits encode genes with housekeeping functions, or with a regulatory role in gene expression, and are located within the same genetic networks as genes included in categories 1 and 2. Given the critical functions in cellular homeostasis in which those genes are implicated, it is conceivable that fully penetrant mutations within those genes could be too deleterious to be compatible with life. Therefore, only variants of a low effect size, with a weak regulatory, disease-modifying role, could be tolerated within those genes. Such genes could be identified through functional screens in which genome-wide, gene-by-gene silencing is followed by a functional readout for a particular trait related to a disease. Our work provides a framework for such a gene discovery strategy that could be applied to complex diseases when conventional approaches are nearing limits of what is achievable.

In conclusion, this study has identified 38 genes that integrate within PD genetic networks and have a regulatory role on the cell-to-cell transfer or a-synuclein. Future perspectives include a more detailed dissection of the functions of those genes and of the ways in which they interact with PD Mendelian and risk genes.

## Acknowledgments

Imaging was performed with support of the Center for Microscopy and Image Analysis, University of Zurich. Flow cytometry was performed with equipment of the flow cytometry facility, University of Zurich. RNA sequencing experiments and basic data analyses were performed at the functional genomics center Zurich (FGCZ). EK is the recipient of an HFSP long term fellowship (LT001044/2017), a Dr. Wilhelm Hurka Foundation project grant and an EMBO long term fellowship (ATLF-815-2014, which is co-funded by the Marie Curie Actions of the European Commission (LTFCOFUND2013, GA-2013-609409)). AA is the recipient of an Advanced Grant of the European Research Council (ERC 250356) and is supported by grants from the Swiss National Foundation (SNF, including a Sinergia grant), the Swiss Initiative in Systems Biology, SystemsX.ch(PrionX, SynucleiX), the Klinische Forschungsschwerpunkte (KFSPs) “small RNAs” and “Human Hemato-Lymphatic Diseases,” and a Distinguished Investigator Award of the NOMIS Stiftung. This work was supported by the UK Dementia Research Institute, which receives its funding from DRI Ltd, funded by the UK Medical Research Council, Alzheimer’s Society and Alzheimer’s Research UK. JH is supported by the Medical Research Council (award number MR/N026004/1), the Wellcome Trust (award number 202903/Z/16/Z), the Dolby Family Fund, the National Institute for Health Research University College London Hospitals Biomedical Research Centre and the BRCNIHR Biomedical Research Centre at University College London Hospitals NHS Foundation Trust and University College London. This work was supported in part by the Intramural Research Programs of the National Institute of Neurological Disorders and Stroke (NINDS), the National Institute on Aging (NIA), and the National Institute of Environmental Health Sciences both part of the National Institutes of Health, Department of Health and Human Services; project numbers 1ZIA-NS003154, Z01-AG000949-02 and Z01-ES101986. The funders played no role in study design, data collection and analysis, decision to publish, or preparation of the manuscript. We would like to thank Gilles Kratzer (UZH Institute of mathematics) for expert advice on statistical analyses, Dr. Kelvin Luk (University of Pennsylvania) for providing us with vials of their HEK QBI wt synuclein cell line, Dr. Berend Snijder for allowing us access to his Opera Phenix for imaging a subset of plates, Dr. Giancarlo Russo for assistance with RNA sequencing analysis and data management, and Rita Moos and Jacqueline Wiedler for assistance with ordering consumables and grant administration.

## Statement of contribution

Acquired funding: AA, EK, BTH, JH, MR, PAL. Supervised study: AA, BTH, JH, MR, PAL, JB, EK, AT. Performed clonings: EK, ZF, JDM, AW. Performed and optimized HTS imaging: EK, CA. Optimized and performed wet lab experiments for HTS: EK. Performed and optimized confocal imaging: EK. Printed siRNAs: EK, MC, ME, MA, DH, ACarrella, DS, LM, DP, AW. Analyzed GWAS data: SBC, CB. Analyzed gene expression data: JB, EK, KD, RR, MR, SGR. Analyzed protein protein network interaction data: CM, PAL. Performed flow cytometry: EK, AW. Performed FACS: ML, VL, EK, AW. Performed western blot: EK. Performed qPCR: EK. Performed tissue culture: EK, AW, SCH, JDM. Optimized and maintained robotics and provided critical advice on robotics usage: ME. Analyzed wet lab data: EK, ACrimi, AW, AChincisan. Wrote code for analysis of HTS data: ACrimi, AChincisan. RNA sequencing: AM, DH, MA, EK. Wrote the manuscript: EK, AA. Edited manuscript: all authors.

## Declaration of interests

The authors declare no competing interests.

## SUPPLEMENTARY TABLES LEGENDS

**Supplementary table 1:**
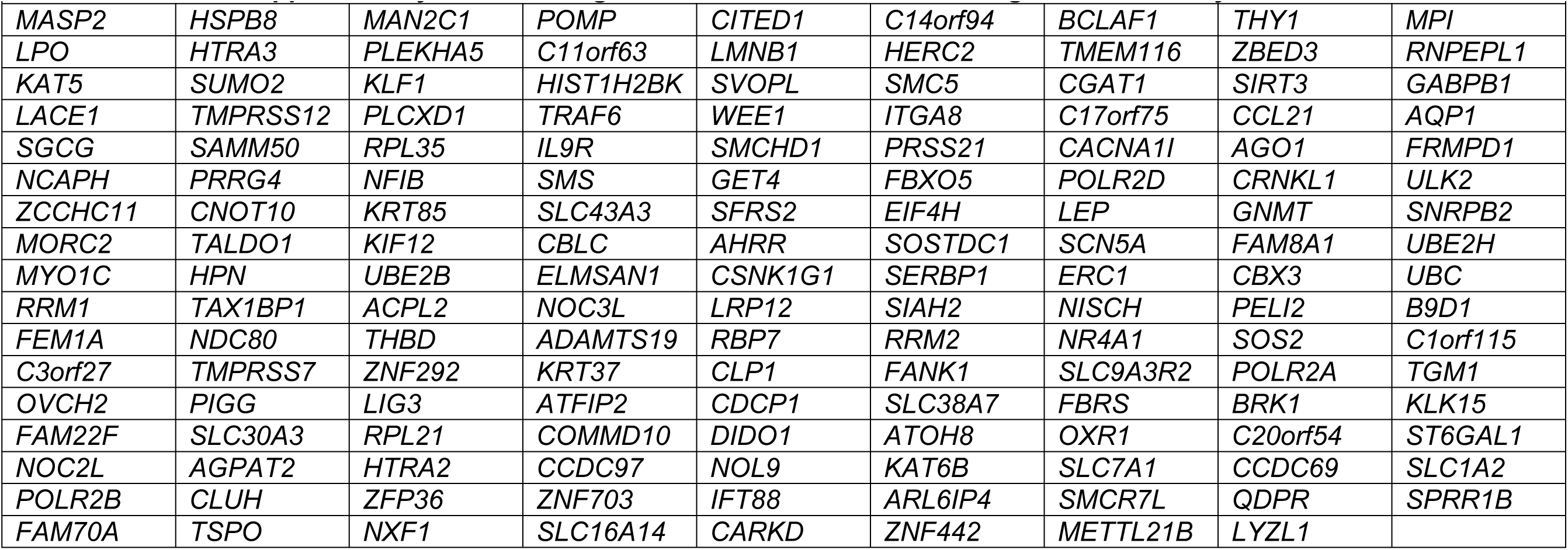
152 genes that were confirmed through the secondary screen.

**Supplementary table 2:**
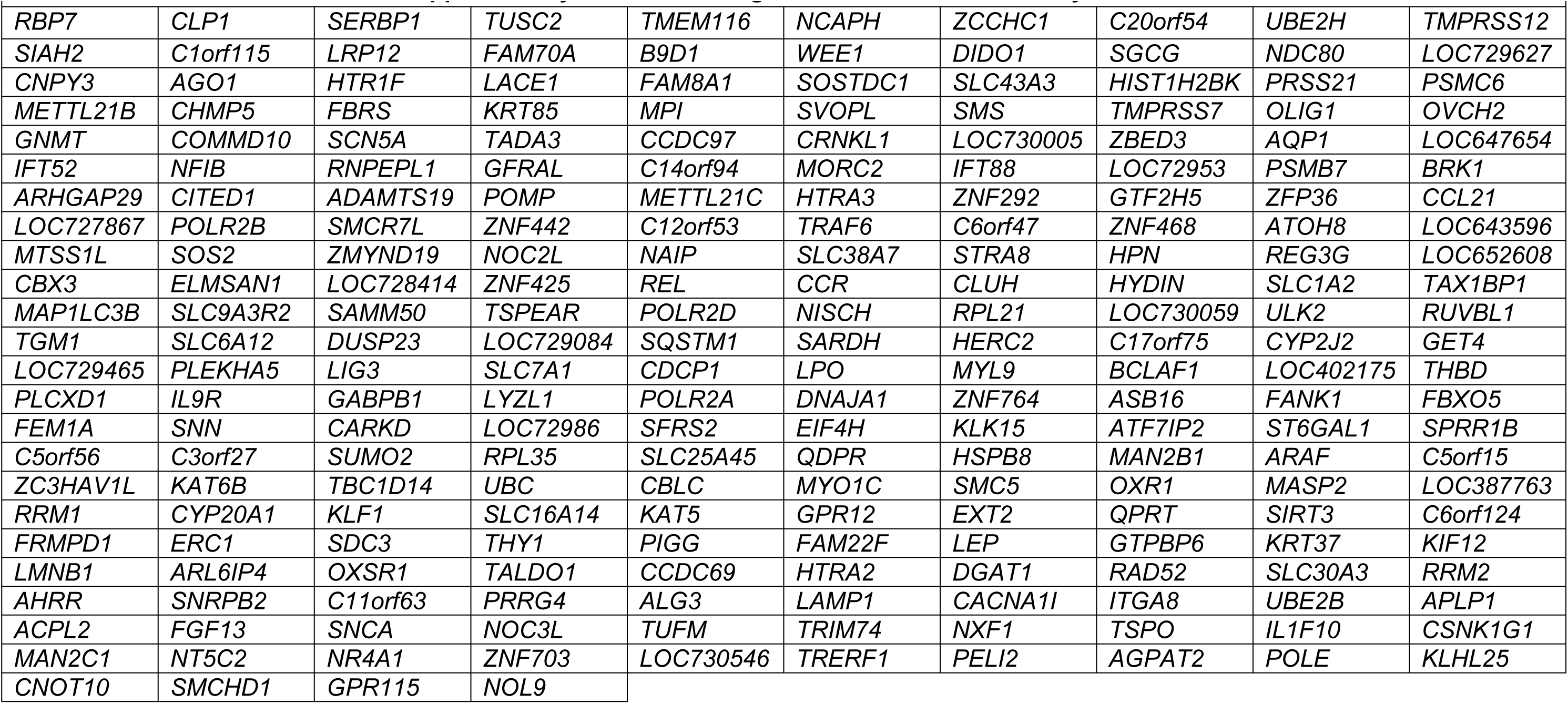
List of genes included in the tertiary screen. Those genes were the 152 genes that were confirmed through the secondary screen, 80 random genes and *SNCA*.

**Supplementary table 3:** List of genes that modify the cell-to-cell transfer of RFP, as well as aSyn-RFP.

|  |
| --- |
| **Supplementary table 3:**<br>**Genes that regulate the**<br>**cell-to-cell transfer of**<br>**both aSyn-RFP and RFP.** |
| *POLRA* |
| *FBXO5* |
| *UBC* |
| *RRM1* |
| *CITED1* |
| *MYOC1* |
| *GNMT* |
| *SCN5A* |
| *ZCCHC11* |
| *SLC7A1* |
| *DIDO1* |
| *AHRR* |
| *LRP12* |
| *SMS* |

**Supplementary table 4:**
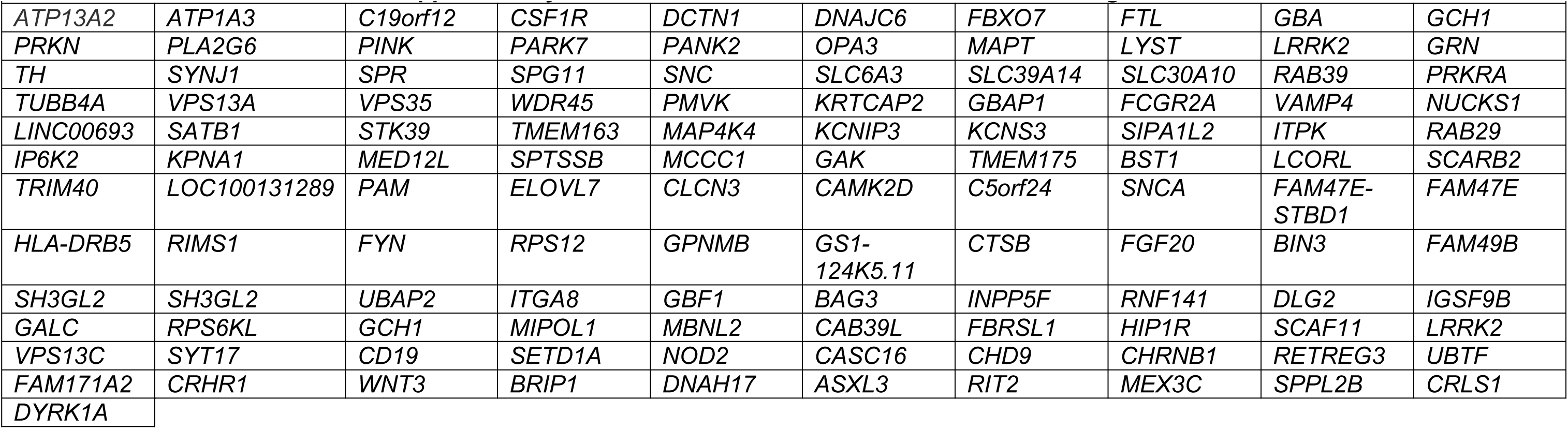
List of known PD Mendelian and GWAS genes included in the WGCNA and WPPNIA analyses.

**Supplementary table 5:**
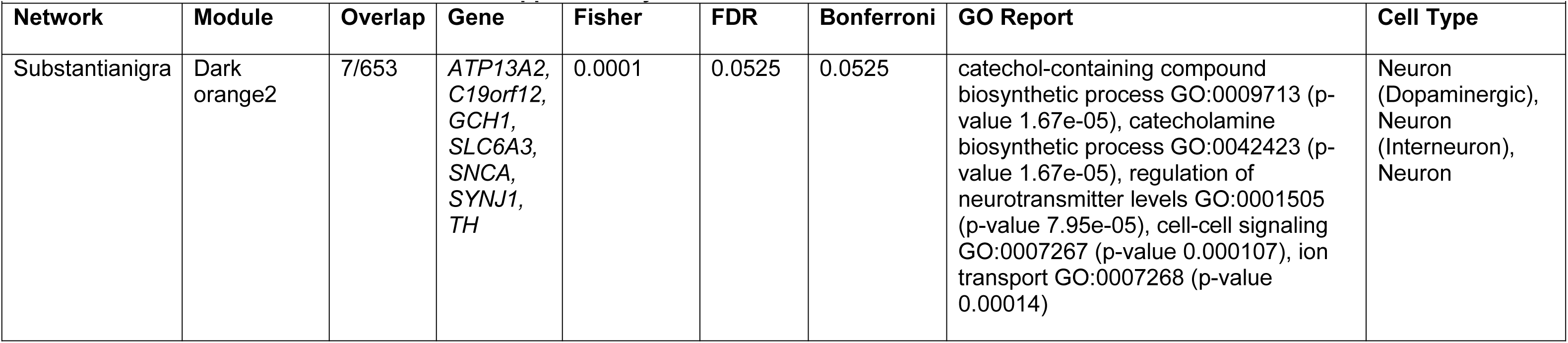
WGCNA for known PD genes. Overlap: number of hit genes relatively to the total size of the module; GO report: GO enrichment analysis results; cell type: enrichment for particular cell type-specific markers in each module; void: not enriched for particular cell markers.

**Supplementary table 6:**
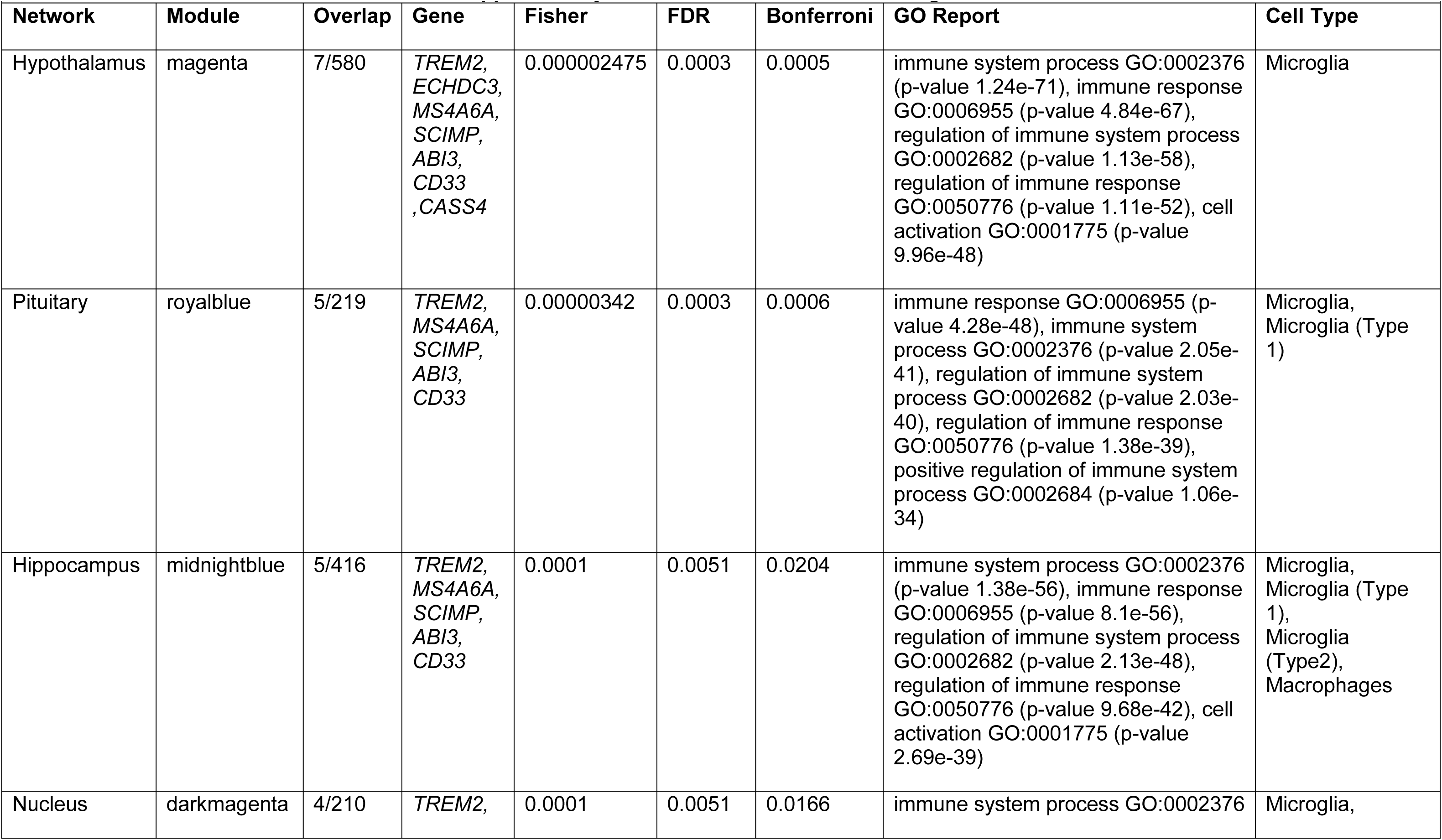
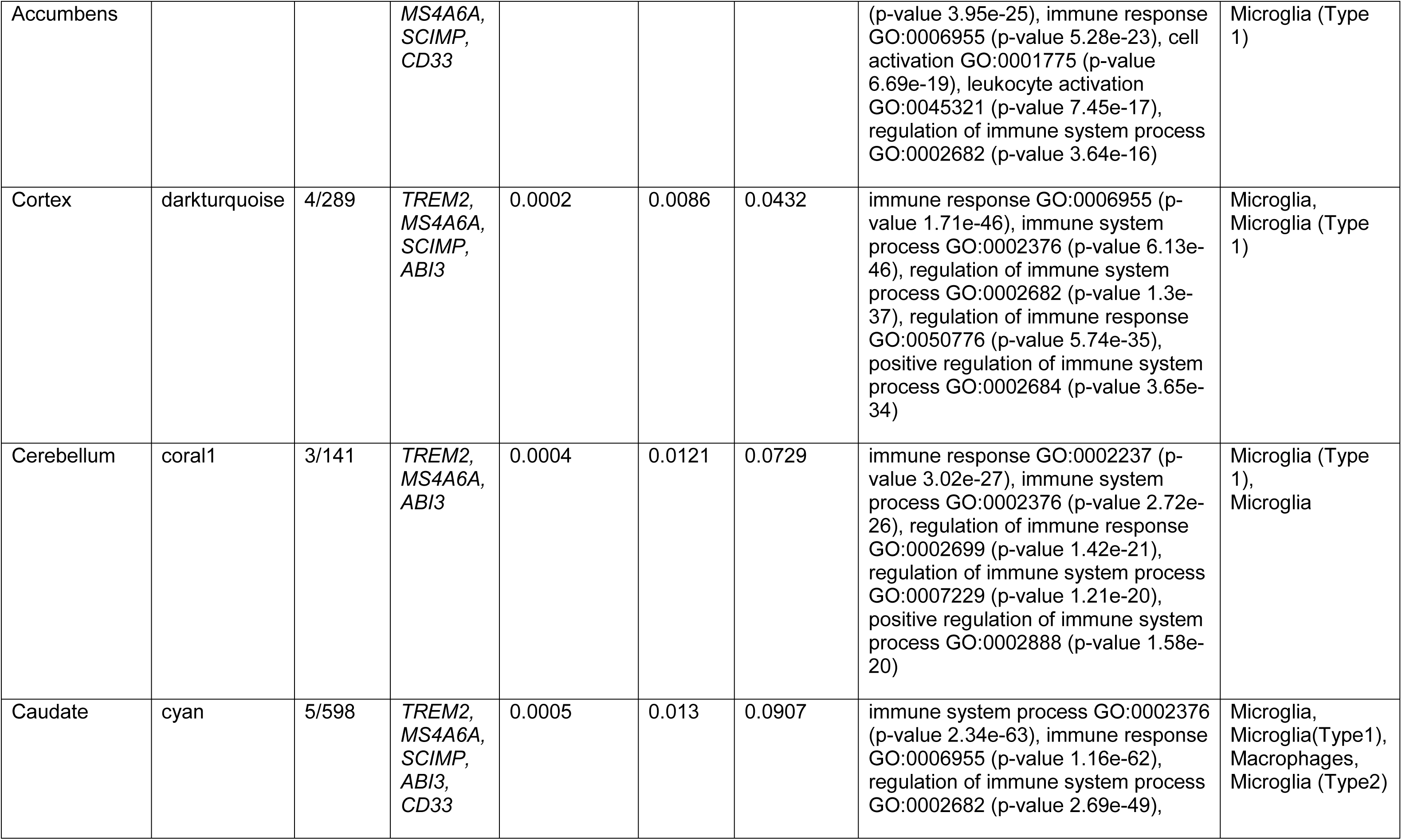
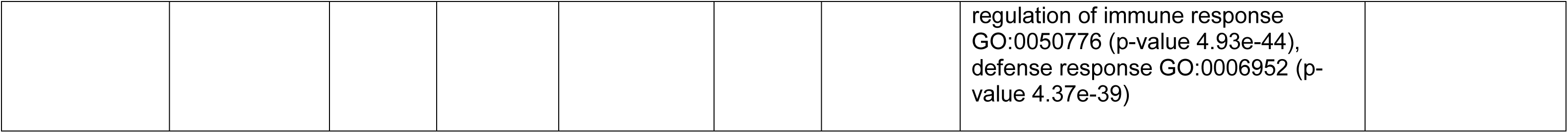
WGCNA for known AD genes. Overlap: number of hit genes relatively to the total size of the module; GO report: GO enrichment analysis results; cell type: enrichment for particular cell type-specific markers in each module; void: not enriched for particular cell markers.

**Supplementary table 7:** Hits present in the same modules as PD Mendelian and GWAS genes.

| **Gene ID** | **Gene Symbol** |
| --- | --- |
| ENSG00000109944 | *C11orf63* |
| ENSG00000134744 | *ZCCHC11* |
| ENSG00000173145 | *NOC3L* |
| ENSG00000063438 | *AHRR* |
| ENSG00000145808 | *ADAMTS19* |
| ENSG00000139514 | *SLC7A1* |
| ENSG00000147650 | *LRP12* |
| ENSG00000132361 | *CLUH* |
| ENSG00000106052 | *TAX1BP1* |
| ENSG00000128731 | *HERC2* |
| ENSG00000108666 | *C17orf75* |
| ENSG00000167325 | *RRM1* |
| ENSG00000047315 | *POLR2B* |
| ENSG00000029363 | *BCLAF1* |
| ENSG00000145781 | *COMMD10* |
| ENSG00000112029 | *FBXO5* |
| ENSG00000162231 | *NXF1* |
| ENSG00000240583 | *AQP1* |
| ENSG00000181222 | *POLR2A* |
| ENSG00000198624 | *CCDC69* |
| ENSG00000198887 | *SMC5* |
| ENSG00000156860 | *FBRS* |
| ENSG00000124713 | *GNMT* |
| ENSG00000101596 | *SMCHD1* |
| ENSG00000150991 | *UBC* |
| ENSG00000169692 | *AGPAT2* |
| ENSG00000183873 | *SCN5A* |
| ENSG00000102172 | *SMS* |
| ENSG00000052126 | *PLEKHA5* |

**Supplementary table 8:** WPPNIA networks for hits, PD Mendelian and PD GWAS genes. The tabs in the table are as follows: Seeds: 34 hit genes, 18 PD Mendelian genes and 65 PD GWAS genes forming the input for the WWPNIA analysis; Experimental_Network_filtered: First layer interactors of 34 hits; PD_network_filtered: first layer interactors of PD Mendelian genes; GWAS_network_filtered: first layer interactors of PD GWAS genes. Terms used: NameA: seed genes; NameB: first layer interactors of seeds; SwissA, EntrezA, SwissB, EntrezB: identifiers of genes listed in columns NameA and NameB; Method.Score: scoring of the strength of published evidence based on the assessment of the methodology used; Publication.Score: scoring of the strength of the evidence based on the number of publications found; Final.Score: composite score of the 2 scoring methods.

**Supplementary table 9:** WPPNIA overlaps between the network of the hits with the network for PD Mendelian genes (tab: Attributes_EXP-PD), and between the network of hits with the network for PD GWAS genes (Attributes_EXP-GWAS). Definition of terms used: Nodes: all genes belonging to the network formed by the 34 hits (34 seeds and first layer interactors, for a total of 615 nodes); Experimental network: role of each gene in the network formed by the hits (seed: one of 34 hits, direct_interactor_of_hit: first layer interactor of hit gene); PD_Network: role of the genes in the network formed by PD Mendelian genes (interactor_PD: that gene is a first layer interactor of the PD Mendelian genes used as seeds); GWAS_Network: role of the genes in the network formed by PD GWAS risk genes (interactor_GWAS: that gene is a first layer interactor of the PD GWAS genes used as seeds); NA: not applicable; communal: common genes between the network of hits and either the network formed by PD Mendelian genes or PD GWAS genes; SEED_EXP: one of 34 hit genes; experimental: one of the first layer interactors of the hits; seed_GWAS: one of the 65 PD GWAS genes used as input for the WPPNIA.

**Supplementary table 10:** GO enrichment analyses for the 50 genes that are in common between the 3 networks (hits, PD Mendelian, PD GWAS genes), as depicted in figure 7c. GO terms have been grouped in semantic classes by semantic similarities and general semantic classes (in light grey) have been removed from the analysis. Intersections: genes belonging to each GO term; term size: number of genes belonging to each GO term category; intersection size: number of genes from the 50 submitted genes that belonged to each GO term category.

**Supplementary table 11:** Burden per hit gene, as assessed by analyzing pre-existing GWAS data. Three different minor allele frequencies (MAFs) were assessed per gene. MAF, minor allele frequency; Q, Q statistic; Perm Pvalue, permuted P value based on 1000 permutations; the following nominated genes did not have enough variants to perform gene-based analyses *ADAMTS19, AQP1, CITED1, CLUH, COMMD10, DIDO1, FBXO5, RRM2, SMCHD1, SMC5, SMS, TAX1BP1*. Please note that supplementary tables 8-10 can be found on Github under the following links: https://github.com/alecrimi/aSynuclein_siRNA_screen/blob/master/supplementary%20table%208%20191219.xlsx https://github.com/alecrimi/aSynuclein_siRNA_screen/blob/master/supplementary%20table%209%20191219.xlsx https://github.com/alecrimi/aSynuclein_siRNA_screen/blob/master/supplementary%20table%2010%20191219.xlsx

|  | MAF ≤ 0.05 |  |  |  |  | MAF ≤ 0.03 |  |  |  |  | MAF ≤ 0.01 |  |  |  |  |
| --- | --- | --- | --- | --- | --- | --- | --- | --- | --- | --- | --- | --- | --- | --- | --- |
| Gene | NumVar | Q | Pvalue | Stat | PermPvalue | NumVar | Q | Pvalue | Stat | PermPvalue | NumVar | Q | Pvalue | Stat | PermPvalue |
| AGPAT2 | 110 | 314175 | 0.742231 | 314175 | 0.775795 | 110 | 314175 | 0.742231 | 314175 | 0.775795 | 109 | 310291 | 0.585821 | 310291 | 0.670691 |
| AHRR | 5 | 780220 | 0.0531187 | 780220 | 0.0532 | 5 | 780000 | 0.0531187 | 780220 | 0.0532 | 4 | 779553 | 0.0293771 | 779553 | 0.0282 |
| ARL6IP4 | 4 | 139764 | 0.166394 | 139764 | 0.154273 | 4 | 140000 | 0.166394 | 139764 | 0.154273 | 4 | 139764 | 0.166394 | 139764 | 0.154273 |
| B9D1 | 102 | 1150000 | 0.498234 | 1150000 | 0.551876 | 100 | 1150000 | 0.450802 | 1150000 | 0.470588 | 98 | 282857 | 0.850827 | 282857 | 0.881057 |
| BCLAF1 | 197 | 362224 | 0.993425 | 362224 | 0.98912 | 197 | 362224 | 0.993425 | 362224 | 0.98912 | 195 | 331630 | 0.969203 | 331630 | 0.970874 |
| C11orf63 | 595 | 2.00E+07 | 0.30269 | 2.00E+07 | 0.369549 | 588 | 16700000 | 0.344692 | 16700000 | 0.405186 | 505 | 2550000 | 0.883885 | 2550000 | 0.938086 |
| C17orf75 | 43 | 139639 | 0.487042 | 139639 | 0.546747 | 43 | 140000 | 0.487042 | 139639 | 0.546747 | 42 | 62455.9 | 0.690898 | 62455.9 | 0.757002 |
| CCDC69 | 1 | 593087 | 0.00558966 | 593087 | 0.0053 | - | - | - | - | - | - | - | - | - | - |
| FBRS | 3 | 1686.35 | 1 | 1686.35 | 0.943396 | 3 | 1686.35 | 1 | 1686.35 | 0.943396 | 3 | 1686.35 | 1 | 1686.35 | 0.943396 |
| GNMT | 1 | 90813.2 | 0.259477 | 90813.2 | 0.27894 | 1 | 90800 | 0.259477 | 90813.2 | 0.27894 | - | - | - | - | - |
| HERC2 | 688 | 18300000 | 0.235104 | 18300000 | 0.253807 | 669 | 5900000 | 0.254667 | 5900000 | 0.229674 | 659 | 5160000 | 0.195998 | 5160000 | 0.17059 |
| ITGA8 | 27 | 3140000 | 0.196073 | 3140000 | 0.191131 | 27 | 3140000 | 0.196073 | 3140000 | 0.191131 | 5 | 10186.4 | 1 | 10186.4 | 0.949668 |
| LRP12 | 17 | 3310000 | 0.0999204 | 3310000 | 0.10009 | - | - | - | - | - | - | - | - | - | - |
| MYO1C | 264 | 11100000 | 0.176478 | 11100000 | 0.311818 | 249 | 8730000 | 0.0191883 | 8730000 | 0.0556 | 227 | 2070000 | 0.131282 | 2070000 | 0.209161 |
| NOC3L | 26 | 670955 | 0.661473 | 670955 | 0.676133 | 26 | 670955 | 0.661473 | 670955 | 0.676133 | 15 | 495276 | 0.284418 | 495276 | 0.268384 |
| NXF1 | 4 | 162807 | 0.346761 | 162807 | 0.325945 | 4 | 162807 | 0.346761 | 162807 | 0.325945 | 3 | 159022 | 0.103421 | 159022 | 0.0895 |
| PLEKHA5 | 14 | 409829 | 0.317075 | 409829 | 0.316256 | 14 | 409829 | 0.317075 | 409829 | 0.316256 | 14 | 409829 | 0.317075 | 409829 | 0.316256 |
| POLR2A | 25 | 635251 | 0.672459 | 635251 | 0.687758 | 7 | 400684 | 0.306258 | 400684 | 0.290529 | 4 | 73410.9 | 0.506109 | 73410.9 | 0.530223 |
| POLR2B | 1 | 18561.8 | 0.49331 | 18561.8 | 0.474608 | 1 | 18561.8 | 0.49331 | 18561.8 | 0.474608 | 1 | 18561.8 | 0.49331 | 18561.8 | 0.474608 |
| RRM1 | 313 | 6220000 | 0.790566 | 6220000 | 0.817661 | 280 | 1770000 | 0.925395 | 1770000 | 0.957854 | 265 | 1180000 | 0.789328 | 1180000 | 0.841043 |
| SCN5A | 13 | 242494 | 0.739297 | 242494 | 0.734754 | 10 | 45051.1 | 0.95417 | 45051.1 | 0.95511 | 10 | 45051.1 | 0.95417 | 45051.1 | 0.95511 |
| SLC30A3 | 9 | 599339 | 0.336942 | 599339 | 0.339674 | 8 | 599306 | 0.31302 | 599306 | 0.306091 | 1 | 2194.18 | 0.691126 | 2194.18 | 0.687758 |
| SLC7A1 | 774 | 18200000 | 0.622987 | 18200000 | 0.690131 | 717 | 7840000 | 0.268829 | 7840000 | 0.356252 | 703 | 4920000 | 0.2953 | 4920000 | 0.383436 |
| UBC | 44 | 116129 | 0.978811 | 116129 | 0.978474 | 42 | 147605 | 0.875424 | 147605 | 0.878735 | 41 | 112852 | 0.77982 | 112852 | 0.783699 |
| WEE1 | 97 | 1830000 | 0.0051296 | 1830000 | 0.008 | 94 | 1440000 | 0.00902511 | 1440000 | 0.0155 | 90 | 669541 | 0.0284746 | 669541 | 0.0404 |
| ZCCHC11 | 4 | 34259.2 | 0.760374 | 34259.2 | 0.745712 | 4 | 34259.2 | 0.760374 | 34259.2 | 0.745712 | - | - | - | - | - |

## SUPPLEMENTARY FIGURE LEGENDS

**Supplementary figure 1:**
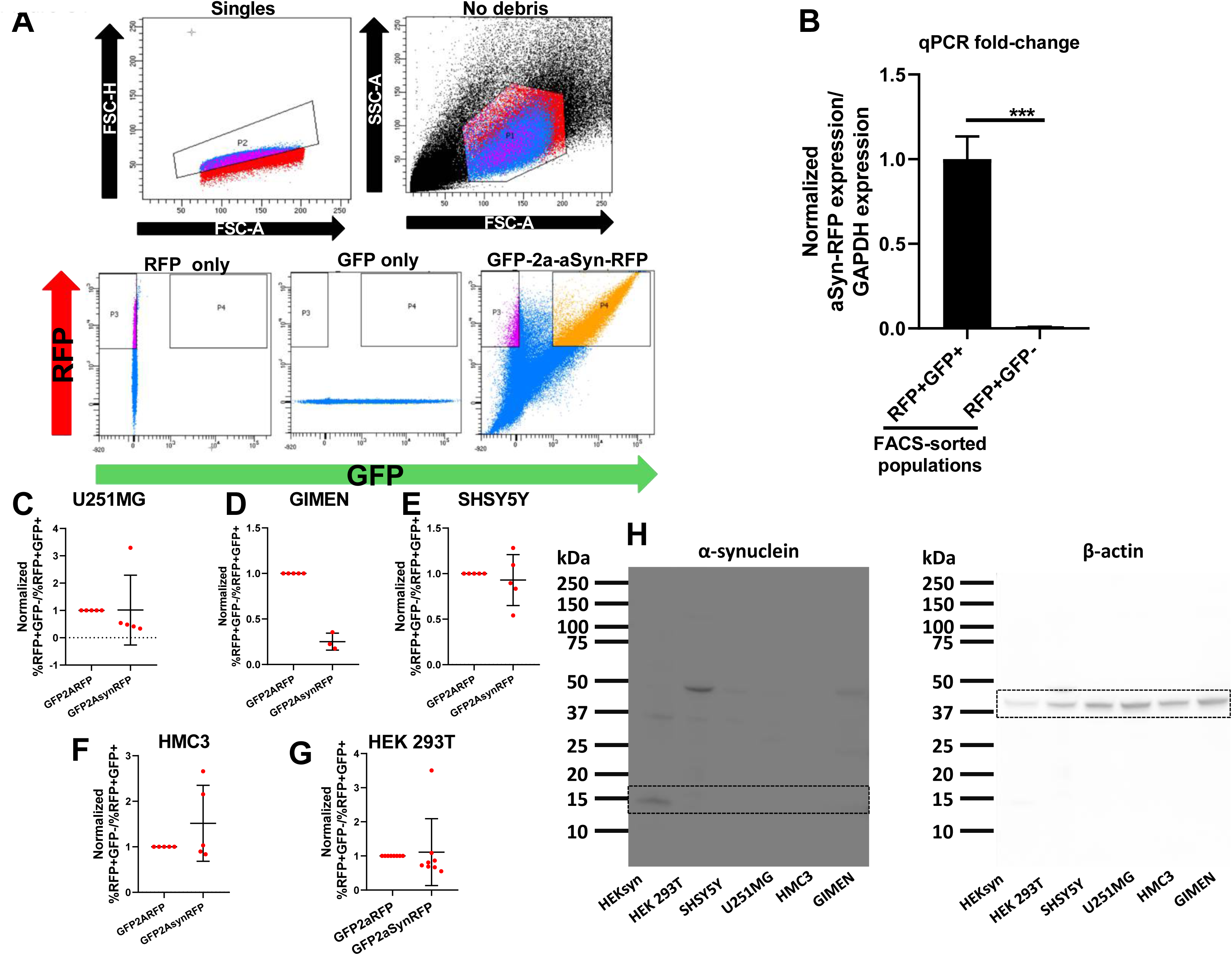
a) FACS plots showing the gating strategy used for sorting the GFP+RFP+ and RFP+GFP-populations. b) qPCR results showing that the aSyn-RFP mRNA levels within the RFP+GFP-cells were negligible in comparison to those within RFP+GFP+ cells. Those results confirm that the aSyn-RFP fusion protein got into the RFP+GFP-cells through cell-to-cell transfer rather than through aberrant expression of the genetic reporter. One sample t-test. *=0.05<p-value<0.01, **=0.01<p-value<0.001, ***=0.001<p-value<0.0001, ****=p-value<0.0001. c,d,e,f,g) Cell-to-cell transfer ratio of RFP versus aSyn-RFP in various cell lines (U251MG, GIMEN, SHSY5Y, HMC3, HEK 293T). In none of the cell lines tested did the fusion protein aSyn-RFP transfer significantly more than the RFP protein alone. h) Western blot indicating a-synuclein protein levels in cell lysates from the various cell lines tested in panels c,d,e,f and g. None of the cell lines showed detectable levels of a-synuclein protein. The HEK-aSyn line was used as a positive control. The membranes were probed using an anti-aSynuclein antibody.

**Supplementary figure 2:**
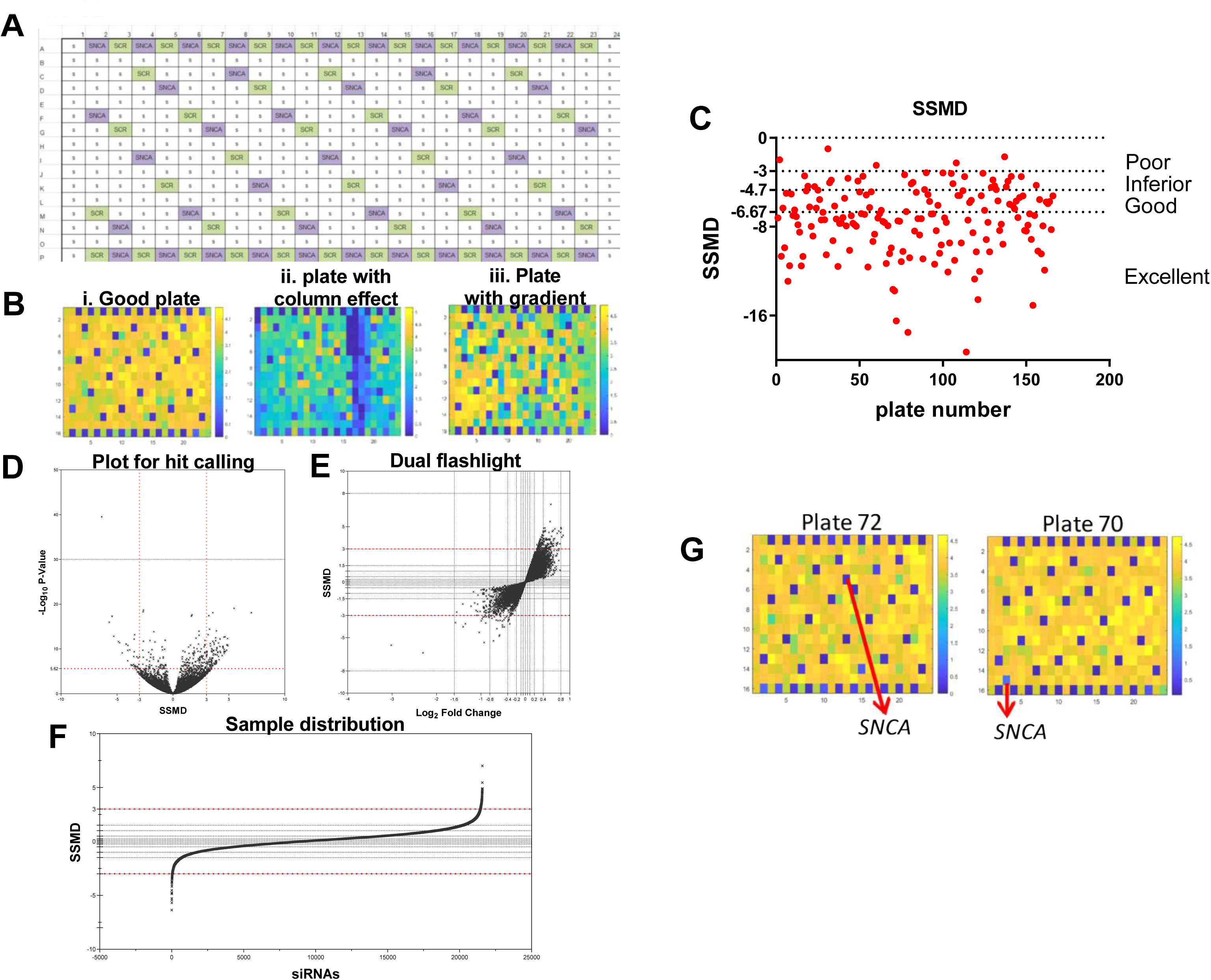
a) Plate layout of the destination plates used in the high throughput screen. The positive (*SNCA*) and negative (scrambled) siRNAs were distributed across the plate to enable quality control. 44 positive and 44 negative controls were used per plate. b) Representative heatmaps. i) A good quality plate, ii) a plate with a column effect, possibly indicating a printing problem of the plasmid or a problem during cell seeding, iii) a plate with a temperature gradient. c) Plot depicting the SSMD values per plate for all 166 plates included in the primary screen. Each dot represents one plate. The cutoffs were set assuming a very strong positive control, as previously described (Zhang, 2008) (0 to -3: poor; -3 to -4.7: inferior; -4.7 to -6.67: good; >-6.67: excellent). d) Plot depicting the SSMDs versus –log10(p-value) for each of the siRNAs. e) Dual flashlight plot depicting the log2(fold change) versus the SSMD for each siRNA. f) Sample distribution plot. The siRNAs were ranked by SSMD and plotted in the ranked format. g) Heatmaps of the 2 plates including the *SNCA* gene. The wells including the library siRNAs against *SNCA* are readily identifiable by visual inspection.

**Supplementary figure 3:**
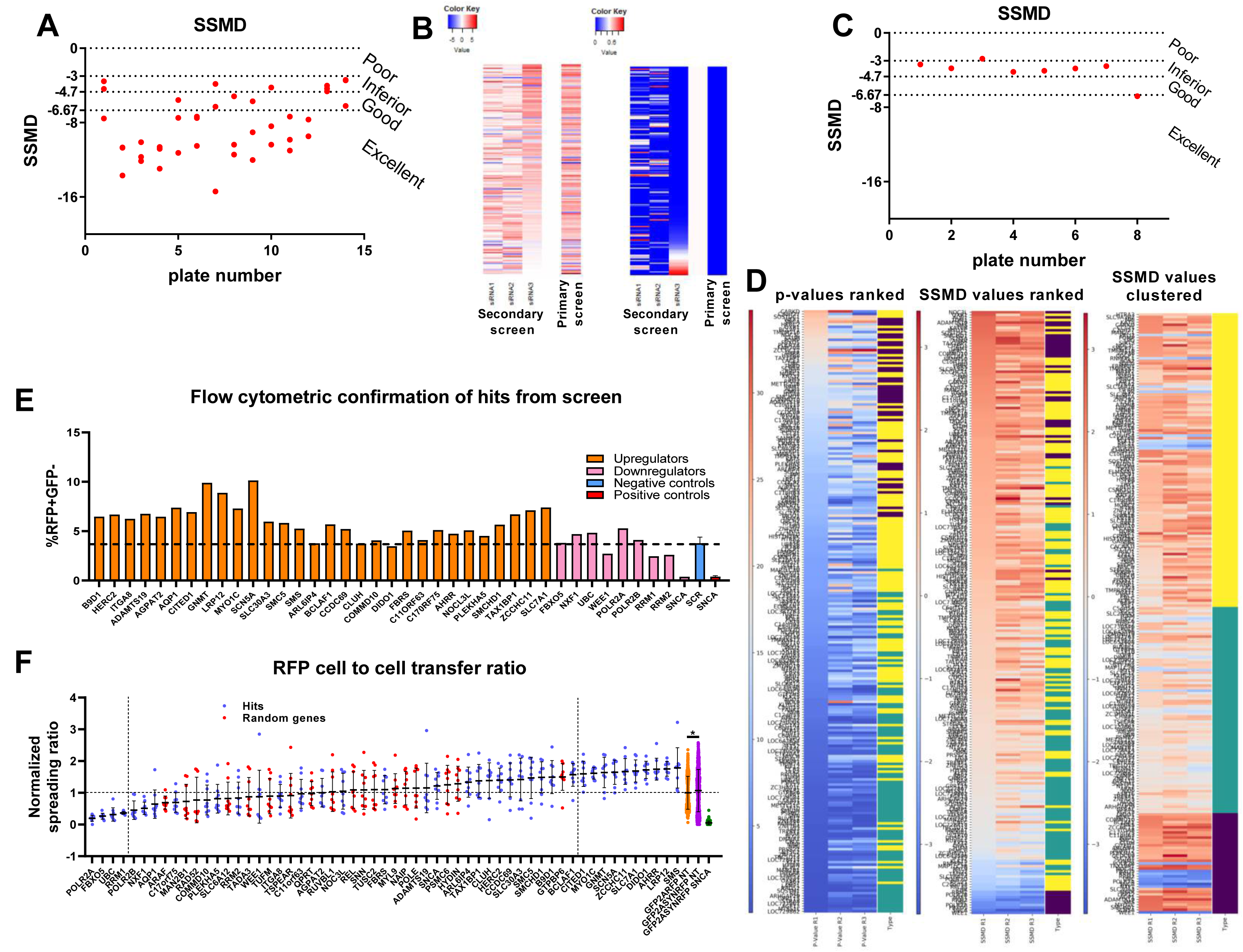
a) Plot depicting the SSMDs of all plates included in the secondary screen (assessment of SSMD values: 0 to -3: poor; -3 to -4.7: inferior; -4.7 to -6.67: good; >-6.67: excellent). b) Heatmaps for the SSMD (left) and p-values (right) per gene for the 152 hits confirmed through the secondary screen. The same metrics for the same genes from the primary screen are shown for reference. Each of the 3 siRNAs targeting the same transcript is shown separately. c) Plot showing the SSMD per plate for one of the 3 tertiary screens (assessment of SSMD values: 0 to -3: poor; -3 to -4.7: inferior; -4.7 to -6.67: good; >-6.67: excellent). d) Heatmaps showing the p-values and SSMDs for each of the 233 genes included in the tertiary screens. Color code: purple=39 hits, green= 114/152 filtered out genes, yellow= random genes. On the first 2 heatmaps, the genes have been ranked by p-value and SSMD, respectively. On the heatmap on the right, the genes have been clustered based on their status (i.e. belonging to one of the following 3 groups: 39 hits, 80 random genes, 114 genes identified through the secondary screen that did not pass the cutoffs of the tertiary screens). e) Validation of 39 hits generated after the tertiary screens through flow cytometry. 82% of the genes had a directionality of effect that was consistent with that seen in the imaging-based screen. f) Plot showing the rate of cell-to-cell transfer of RFP. Significance level cutoffs (Bonferroni corrected p-values with a=0.05) are depicted on the plot in the pink boxes. This screen was completed on the HEK-aSyn cell line.

**Supplementary figure 4:**
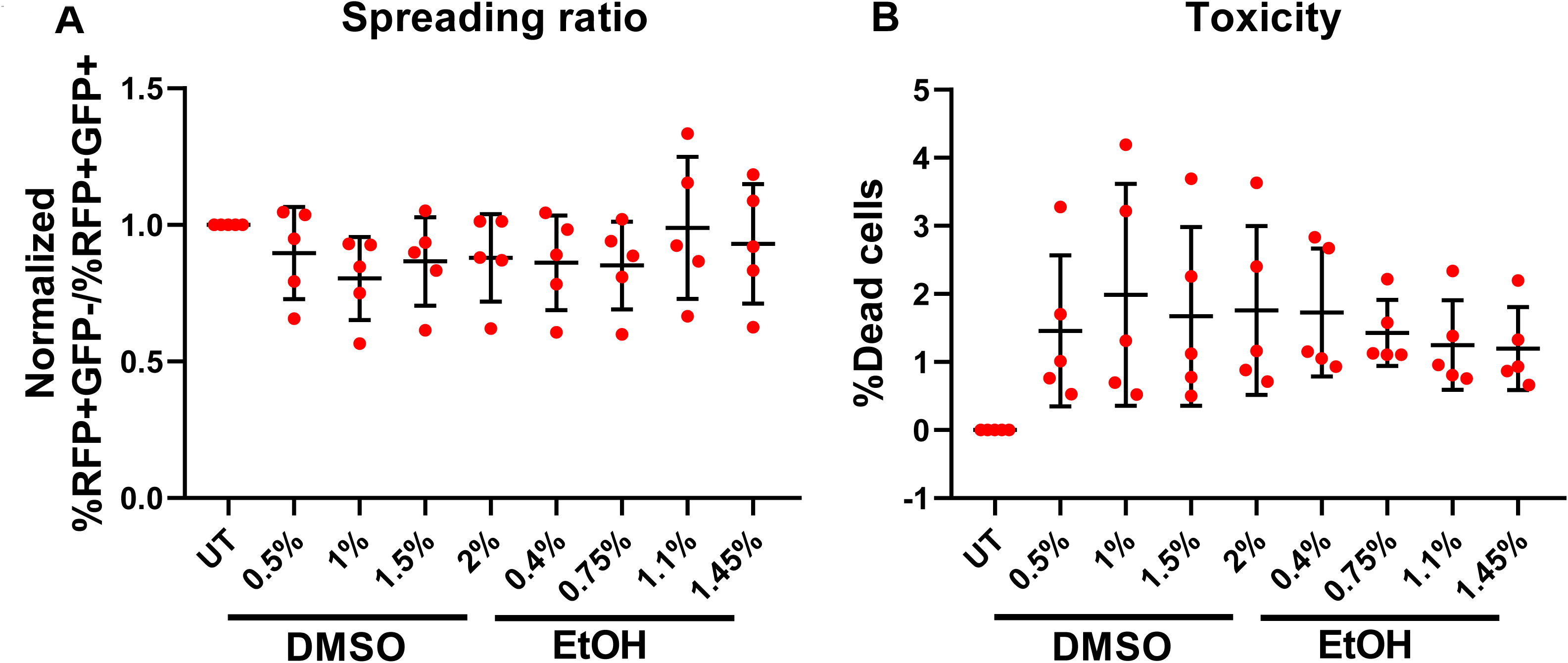
a) Effect of various concentrations of DMSO and ethanol on the cell-to-cell transfer of a-synuclein. Increasing the concentration of the vehicle did not affect the rate of the cell-to-cell transfer of the asyn-RFP fusion protein (i.e. no dose-response effect was seen for the vehicle-only). b) Toxicity levels of various concentrations of the vehicles. No significant difference was observed between concentrations. % indicates v/v% of DMSO or ethanol in the culture medium of the cells. ETOH=ethanol.

**Supplementary figure 5:**
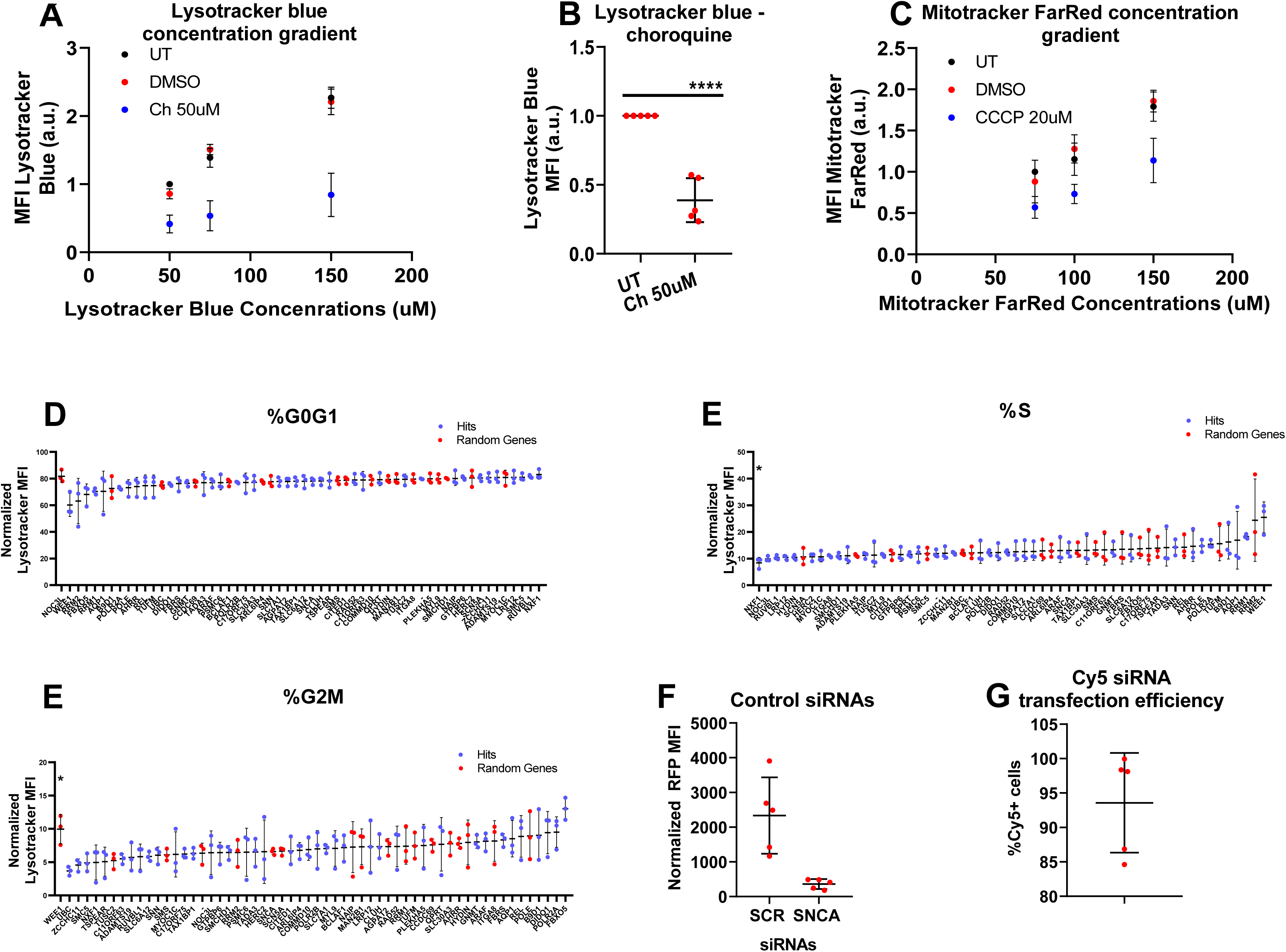
Dyes of different colors were used to confirm the results from the lysotracker and mitotracker assays. a) The mean fluorescence intensity (MFI) of LysoTracker blue showed a linear dose response to the concentration of the dye, and the positive (chloroquine) and negative (untreated) controls were well-separated. b) The positive control (chloroquine) significantly reduced the MFI of LysoTracker blue in comparison to the untreated samples. The experiment was repeated 5 times independently, data was internally normalized to the negative control prior to metaanalysis. Statistical analysis was performed with one sample t test. *=0.05<p-value<0.01, **=0.01<p-value<0.001, ***=0.001<p-value<0.0001, ****=p-value<0.0001. c) The MFI of mitotracker far red showed a linear dose response to the concentration of the dye, and the positive (CCCP) and negative (DMSO) controls were well separated. d,e,f) Percentage of the cells in each phase of the cell cycle. The experiment was performed 3 times independently and the data was normalized internally to the negative control prior to metaanalysis. Statistical analysis was performed with one sample t test, followed by Bonferroni correction for multiple testings. *=0.05<p-value<0.01, **=0.01<p-value<0.001, ***=0.001<p-value<0.0001, ****=p-value<0.0001. Optimizations of siRNA transfections with 3 days’ incubation. g) HEK-aSyn cells were co-transfected with the GRP-2a-aSyn-RFP construct plus 3 pooled siRNAs against *SNCA* or a scrambled control siRNA. The *SNCA* siRNAs resulted in a 83.86+/-3.6% reduction in the MFI of RFP. The experiment was performed 5 times independently. h) HEK-aSyn cells were co-transfected with the GRP-2a-aSyn-RFP construct plus the siRNA-Cy5. Transfection efficiency was 93.6+/-7.2%. The experiment was performed 5 times independently.

**Supplementary figure 6:**
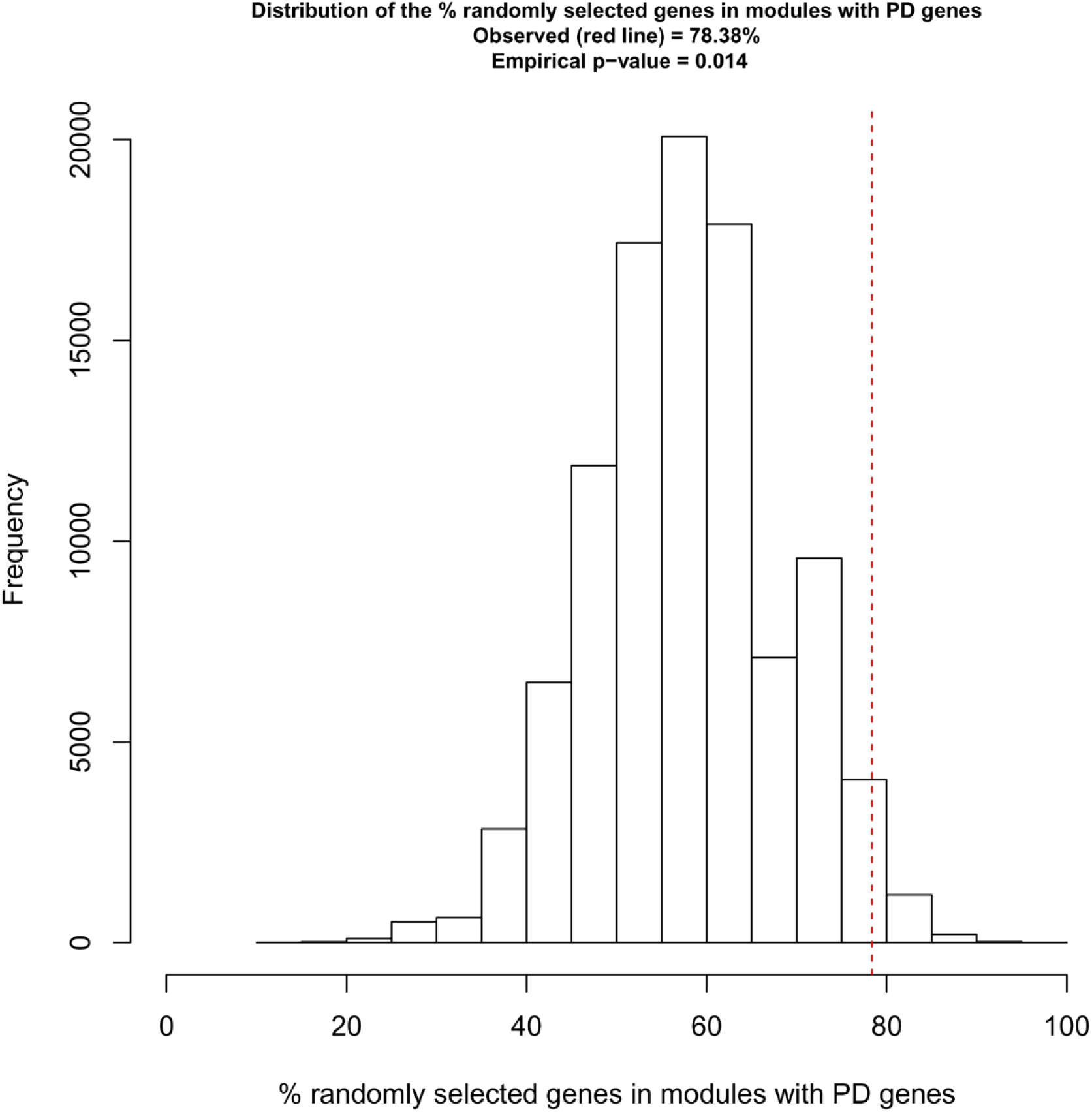
Null distribution of the percentage of genes overlapping with PD genes. 100,000 random simulations were performed, generating 100,000 lists of genes each of which contained 37 genes. Based on the fraction of times that we saw a greater % of overlap of the random gene lists with the PD genes than the % overlap observed for the 37 hits, an empirical p-value was generated. We found that the co-clustering of hits with PD genes occurred more frequently than expected by random chance, with an empirical p-value of 0.014.

**Supplementary figure 7:**
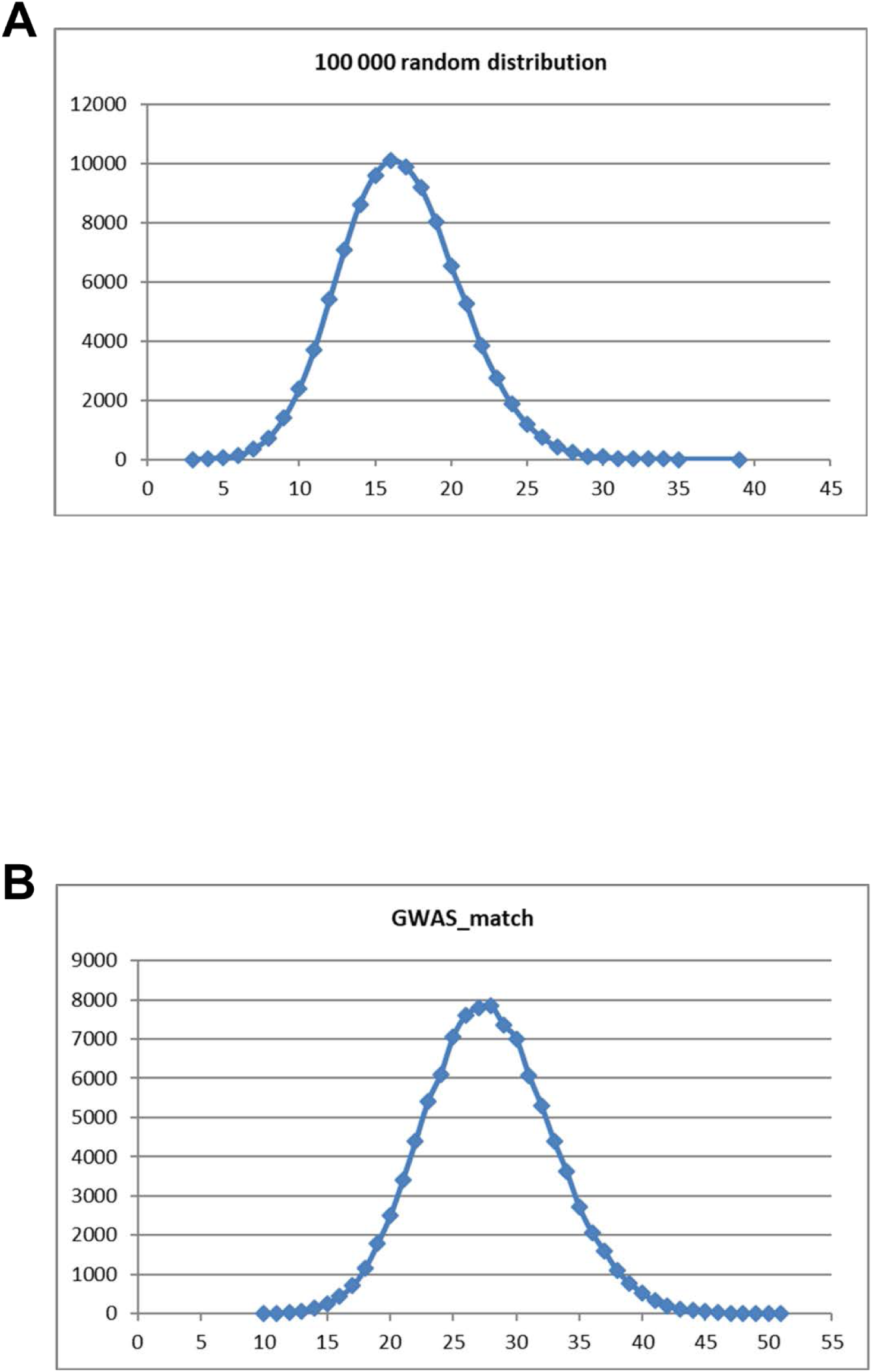
The top panel depicts the normal distribution of the % overlap with Mendelian PD network, while the bottom panel with the PD GWAS risk genes network. a) Random matches (RANDOM) Experimental Network (615 nodes) vs (REAL) PD network (545 nodes) number of real matches = 79; mean of random matches = 16 b) Random matches (RANDOM) Experimental Network (615 nodes) vs (REAL) GWAS network (902 nodes) number of real matches = 132; mean of random matches = 28

## Notes

https://github.com/alecrimi/aSynuclein_siRNA_screen

